# Dermatological plants and their uses in the *Receptarium* of Burkhard III from Hallwyl from 16^th^ century Switzerland – Data mining a historical text and preliminary *in vitro* screening

**DOI:** 10.1101/2024.02.09.579674

**Authors:** Jonas Stehlin, Ina Albert, Thomas Frei, Barbara Frei Haller, Andreas Lardos

## Abstract

**Ethnopharmacological relevance:** Historical texts on *materia medica* can be an attractive source of ethnopharmacological information. In recent decades, various research groups have investigated corresponding resources from Europe and the Mediterranean region, pursuing different objectives. Regardless of the method used in such a work, the indexing of textual information and its conversion into data sets useful for further investigations represents a significant challenge.

**Aim of the study:** First of all, this study aims to systematically catalogue pharmaco-botanical information in the *Receptarium* of Burkhard from Hallwyl (RBH) in order to identify candidate plants in a targeted manner. Secondly, the potential of RBH as a resource for pharmacological investigations will be assessed by means of a preliminary *in vitro* screening.

**Materials and methods:** We developed a relational database for the systematic recording of the parameters composing the medical recipes contained in the historical text. Focusing on dermatological recipes, we explored the mentioned plants and their uses by drawing on specific literature. The botanical identities (candidate species) suggested in the literature for the historical plant names were rated based on their plausibility of being the correct attribution. The historical uses were interpreted by consulting medical-historical and modern clinical literature. For the subsequent *in vitro* screening, we selected candidate species used in recipes directed at the treatment of inflammatory or infectious skin disorders and wounds. Plants were collected in Switzerland and their hydroethanolic crude extracts tested for possible cytotoxic effects and for their potential to modulate the release of IL-6 and TNF in LPS-stimulated whole blood and PBMCs.

**Results:** The historical text analysis points up the challenges associated with the assessment of historical plant names. Often two or more plant species are available as candidates for each of the 161 historical plant names counted in the 200 dermatological recipes in RBH. At the same time, the 56 medicinal uses mentioned in these recipes illustrate, that the details provided in the text about the skin problem in question enable conclusions to be drawn based on which the presumed underlying medical condition and its pharmacological basis can be narrowed down. On this basis, 11 candidate species were selected for *in vitro* screening, four of which were used in RBH in herbal simple recipes and seven in a herbal compound formulation. None of the extracts tested showed a noteworthy effect on cell viability except for the sample of *Sanicula europaea*. Extracts were tested at 50 µg/mL in the whole blood assay, where especially *Vincetoxicum hirundinaria* or *Solanum nigrum* showed inhibitory or stimulatory activities. In the PBMC assay, the root of *Vincetoxicum hirundinaria* revealed a distinct inhibitory effect on IL-6 release (IC_50_ of 3.6 µg/mL).

**Conclusions:** Using the example of RBH, this study illustrates a possible ethnopharmacological path from unlocking the historical text and its subsequent analysis, through the selection and collection of plant candidates to their *in vitro* investigation. Fully documenting our approach to the analysis of historical texts, we hope to contribute to the discussion on solutions for the digital indexing of premodern information on the use of plants or other natural products.

## 1 Introduction

### 1.2 Dermatological uses in historical texts

Skin complaints, wounds or skin manifestations have always plagued humans. Paleodermatological studies of mummies from the 4th millennia BCE to the modern era illustrate that skin lesions, dermatoses, neoplasms or infectious diseases affecting the skin were widespread in all eras and cultures (Lowenstein, 2004). Accordingly, people developed therapeutic measures to counter these health problems early on in history. In the Ebers Papyrus from the 16th century BCE, one of the oldest extant medical texts, numerous dermatological recipes are documented, such as ointments for skin diseases (Eb 104-131), recipes for wound care (Eb 515-542), abscesses (Eb 551-555), ulcers and purulent wounds (Eb 556-591), or leprosy (Eb 773-738) (Popko, 2022). It is therefore of little surprise that in premodern medical texts from Europe and the Mediterranean region dermatological conditions belong to the most important areas of application for medicinal plants (Lardos and Heinrich, 2013; Dal Cero et al., 2023).

An additional circumstance, which makes dermatology particularly attractive in context with the present study, is related to one of the most critical steps in the ethnopharmacological investigation of premodern texts – the interpretation of the medicinal uses mentioned and their translation into modern clinical terminology. In contrast to other health conditions, applications on the skin described in ancient or medieval recipe texts allow us to draw conclusions about the presumed disease being treated in a comparatively straightforward way. As illustrated by a 13th century Byzantine recipe text, the writers of these manuscripts often provide specific details about the skin problem to be treated by a certain recipe (Zipser et al., 2023). Uses such as “if the scar of a wound gets back”, “if it happens in a wound that worms are developing”, “paronychitis, an abscess that arises next to the root of the nail”, point to poorly healing and infected wounds or inflammations of the skin.

Especially since the beginning of the 21st century numerous historical studies have been conducted in ethnopharmacology, some of them with the aim to identify plants or other natural products that could be interesting candidates in the search of new medicines (for an overview of these studies, see Lardos, 2015). More recent studies have also begun to screen plants mentioned in historical sources from Europe for specific activity profiles such as anti-inflammatory, immunomodulatory, anti-infective or antibiotic activities (Wagner et al., 2017; Ulriksen et al., 2022). Among the plants analysed are also such that were used specifically for skin problems including wounds. Only very few studies have so far attempted to reconstruct individual historical recipes to investigate the pharmacological potential of the relevant formulations (Harrison et al., 2015; Connelly et al., 2020). In this context, a multicomponent eye-salve from a 10^th^ century Anglo-Saxon text in fact proved to be a potent antibiotic agent with a specific activity against resistant strains of *Staphylococcus aureus*, but only if the remedy was prepared exactly as described in the medieval text (Harrison et al., 2015). These studies illustrate the potential but also the challenges of working with these resources, on the one hand in the conversion of historical texts into electronic databases amenable to computer-based analyses, or generally in the interpretation and reconstruction of the recipes (Connelly et al., 2020; Harrison and Connelly, 2020).

Beside the above indicated critical step of interpreting historical medicinal uses, an issue of particular difficulty and complexity in the study of premodern texts is the identification of the plants or plant substances mentioned in these resources. As described in the consensus statement on ethnopharmacological field studies (ConSEFS), the commonly applied method in historical studies on herbal *materia medica*, relies on authoritative literature that provides botanical identifications for the plant names in question. At the same time, a critical evaluation of the information provided in these references by taking into account cultural-historical and phytogeographical aspects as well as plant nomenclature is indispensable (Heinrich et al., 2018). However, since relying on existing plant identifications can sometimes be misleading (Riddle, 1996; Raven et al., 2000), alternative methods have recently been proposed (Evergetis and Haroutounian, 2015; Lardos et al., 2024).

### 1.2 Cultural-historical background

During the Renaissance period in Europe, extensive compilations of the knowledge known at the time about the medicinal use of plants and other natural substances were produced. Important examples from Germany are the “Kräuterbuch” by Hieronymus Bock (1539) and that one by Leonhard Fuchs (1543). A contemporary work from Switzerland is Conrad Gesner’s *Historia plantarum*, the incomplete text of which from 1565 was first published in 1750. However, this is less a text with medical- therapeutic content than a botanical encyclopaedia, primarily characterised by its high-quality plant illustrations (Schulze, 2006).

The “Arzneibuch von Hallwyl”, which is the subject of this project, also originates from Switzerland. The oldest known copy of this early New High German text comprising 295 *folia* or 590 manuscript pages is dated to 1580 and was written by Burkhard III from Hallwyl (Frei, 2012). The work can be categorised in the genre of “Hausarzneibücher”, whose target audience consisted primarily of medical laypeople (Fankhauser, 2012). This is also evident from the text’s preface, which mentions that the work is dedicated to the “common man” (Frei Haller, 2005). As with other representatives of this genre, the content is primarily a collection of recipes for the medicinal use of natural substances, especially plants. We refer to the text in this paper as the *Recepatrium* of Burkhard from Hallwyl (RBH).

Little is yet known about the origin of the recipes mentioned in RBH; possible sources include people close to Burkhard III as well as contemporary works with medical content, and some recipes may even have been created by the author himself (Frei Haller, 2005; Fankhauser, 2012). The kind of knowledge contained in RBH is presumably comparable to that of other historical recipe texts from European traditions: These are compilations of earlier and current knowledge that was considered useful based on empirical criteria and within the cultural and historical context (see Horden, 2011 or Nutton, 2013). The fact that up to the 18th century at least 15 manuscript copies were produced and distributed in various regions of Switzerland suggests that RBH must have had a remarkable local significance.

### 1.3 Study objectives

In this study, we are pursuing two major objectives. First, the herbal recipes in RBH, which refer to the treatment of skin complaints or topically manifest disease symptoms (in this paper denoted as dermatological recipes), will be investigated from a botanical and pharmacological point of view. For this purpose, the information contained in the recipes is transferred to an electronic database, which allows qualitative and quantitative data analysis as well as provide the basis for the selection of plant candidates from RBH for subsequent pharmacological investigations. Secondly, to gain a first insight into the potential of the historical text as a resource in this context, a small number of plants from RBH will be selected for preliminary *in vitro* screening in terms of anti-inflammatory or immunomodulatory activities.

## 2 Material and Methods

### 2.1 Original text

For the systematic analysis of the dermatological recipes in the Receptarium of Burkhard III from Hallwyl (RBH) we used the transcription by Fankhauser (2012) of the recipes written by *Hand A* in the manuscript *FA von Hallwyl A 814* housed in the Staatsarchiv Bern (Switzerland) in the family archive “von Hallwyl” under the title “Arzneibuch des Burkhard v. Hallwyl, 1580 Original des Verfassers” (https://www.sta.be.ch/de/start/ueber-uns/staatsarchiv.html).

In addition, we also investigated the famous Hallwyl wound potion (“das recht Hallwylisch wundtrannck”) a compound herbal preparation mentioned on folio 251a of the same manuscript but written by *Hand B* and therefore not included in the transcription by Fankhauser (2012). Our transcription and translation of the text is shown in **Supplementary Material, Table S5**.

### 2.2 Text analysis

In principle we followed the Consensus Statement for Ethnopharmacological Field Studies (ConSEFS) best practice guidelines with special consideration of points relating to historical studies (Heinrich et al., 2018). The main focus of the method used in the text analysis is on the plants and their dermatological uses in RBH. Non-plant ingredients and their uses mentioned in the text were not considered in this study.

#### 2.2.1 Development of the RBH database and data evaluation

For the investigation of the transcribed recipes written by *Hand A* (Fankhauser, 2012) a relational database (MS Access®) was developed, consisting of 20 interlinked tables of one-to-many and one-to-one relationships with a total of 73 data fields (**Supplementary Material, Figure S1**). The database was developed with the aim of qualitatively and quantitatively analysing various components of the recipes and to extract information on plants and their uses in the form of use records. We defined a use record as being a reference within RBH to a specific plant having a particular medicinal use.

To identify dermatological recipes in RBH, Fankhauser’s (2012) transcription was searched manually for recipe headers or relevant passages within the recipes, which mention external applications for the treatment of skin disorders, wounds, injuries, bleedings or other topically manifest disease symptoms as well as internal applications directed at the treatment of these conditions. Each recipe was then assigned a unique signature consisting of the manuscript page (folio), the position of the page in the bound book (a – left side, b – right side), the scan number of the respective digitised manuscript page, the front or back side of the scan printed on both sides (r – recto, v – verso), the recipe number on the page, and, if applicable, the recipe version or supplement. For example, the signature 007b,1r.01 leads to folio 7 on the right side in the bound manuscript (007b), which corresponds to the backside of scan number 1 (1r), the first recipe on that page (01). Recipes can have variants (indicated with “v”, e.g. 007b,1r.01.v01) in terms of ingredients (e.g. substitutes or optional additional ingredients), form of preparation, or medicinal use. They can also have supplements (indicated with “z”, e.g. 007b,1r.02.z01), which mostly concern additional concomitant therapeutic interventions.

From the dermatological recipes first the recipe ingredients (herbal ingredients including the plant part concerned, as well as other ingredients of animal, human or mineral origin) were extracted and recorded in the database. Spelling variants were entered under the same record. For each plant name mentioned in RBH the presumed botanical identity was elucidated following the process described in section 2.2.3. With the aim to separate out non-plant ingredients (i.e. such of mineral, animal or human origin) they were *a priori* assessed mainly with the help of the *Schweizerdeutsches Idiotikon* (Schweizerisches Idiotikon, 2023), which documents and explains terms of the Swiss-German vernacular from the 13th through to the 21st century. Non-plant ingredients were not further considered, because this paper focusses on the plants in RBH.

Next, the recipes were analysed according to their components and the relevant information recorded in separate data fields: recipe type (simple herbal, simple non herbal, compound herbal, compound non herbal, compound mixed, other combination), type of preparation (e.g. decoction, distillation, press-juice), form of administration (e.g. bath, liniment, poultice), way of administration (e.g. oral, topical, rectal), mentioned use, as well as additional possibly relevant details.

In RBH uses referring to the same medical complaint were often reported in differing qualities (spelling or syntax variants) or different therapeutic contexts. To facilitate subsequent data analysis, the mentioned uses were combined and recorded in the data field “Combined use” (e.g., the mentioned uses “für nüw erhabne geschwulst” or “für alle geschwulst” were combined into the use “geschwulst”, or the mentioned uses “wunden die full ist”, “wunden welche full fleisch erzügt” and “alle wunden vnnd schaden […], bewertt das vor fullem fleisch” into the use “wunden – zu fullen wunden, bewertt vor fullem fleisch”).

#### 2.2.2 Interpreting the medicinal uses

For the interpretation of the medicinal uses mentioned in RBH literature documenting and explaining pre-modern and folk terms and concepts of the appropriate cultural and historical background was consulted. We mainly used Höfler’s (1970) *Deutsches Krankheitsnamenbuch* with supporting information from Riecke’s (2004) *Wörterbuch* (*Band 2*) of *Die Frühgeschichte der mittelalterlichen medizinischen Fachsprache im Deutschen,* for the respective German terminology of diseases and symptoms, as well as the *Schweizerdeutsches Idiotikon* for specific terms of the Swiss-German idiom (Schweizerisches Idiotikon, 2023). If there are several possible interpretations which, based on the information in the references, differ in terms of their plausibility, this is indicated accordingly (1 – high, 2 – moderate, 3 – small chance).

Finally, the conclusions were cross-checked with modern clinical tools and the pharmacological interventions for the treatment of the interpreted historical uses were derived. Our key question was, “what are the pharmacological properties of the medication used to treat the dermatological complaint in question?” Using Fitzpatrick’s Dermatology (Kang et al., 2019), we looked up existing pharmacological interventions and compiled the mentioned pharmacological properties of the appropriate medication. The resulting list of pharmacological properties corresponds to the presumed activities required of the associated RBH plants in order for them to be able to exert beneficial effects on the interpreted historical uses. This information serves as a basis for the subsequent selection of test systems for bioactivity investigations (see section 2.3).

For example, one of the interpretations for the historical use “ëyssenn” leads to a purulent abscess, furuncle, or carbuncle. The cross-check from a modern diagnostic perspective reveals that these skin complaints are caused by staphylococcal infections and involve inflammatory conditions sometimes leading to purulent ulcers. As detailed in Kang et al. (2019), Chapter 150 – Superficial cutaneous infections and pyodermas, beside incision and drainage, the pharmacological intervention is based on antibiotic therapy to control infection and support regression of inflammation. Accordingly, the RBH plant(s) used in the recipe(s) for the treatment of “ëyssenn” would have to hold corresponding pharmacological properties in order to exert a beneficial effect on the complaint. The corresponding RBH plant(s) could then be tested for potential antibiotic or anti-inflammatory activities by selection of appropriate pharmacological assays.

#### 2.2.3 Assessing the botanical identity of plant names

For each plant name recorded in RBH database as well as those mentioned in the Hallwyl wound potion, we compiled a list of possible botanical identities previously suggested in the literature. Information was drawn from three different categories of references:

Linguistic and botanical glossaries providing scientific names for vernacular plant names of Swiss-German and other German dialects especially from late Middle Ages to the Renaissance: The online edition of *Schweizerdeutsches Idiotikon* (Schweizerisches Idiotikon, 2023); Marzell’s (1943-1979) *Wörterbuch der deutschen Pflanzennamen*, and Fischer’s (2001) *Mittelalterliche Botanik*.
Pharmaco-botanical reference texts associated with RBH: Leonhard Fuchs’ *New Kreutterbuch* from 1543 and Hieronymus Bock’s *New Kreutterbuch* from 1539 and 1546. The botanical identifications suggested for the plants described in the two herbals were adopted from Dobat and Dressendörfer (2016) and Hoppe (1969), respectively.
Encyclopaedia of the history of pharmaceutical drugs: Volume 5, parts 1 to 3 of Schneider’s (1974) *Lexikon zur Arzneimittelgeschichte* provides pharmacognostic and historical details about herbal drugs and offers a critical evaluation of the proposed botanical identities of these materials.

Any botanical taxon suggested in the literature was included as a candidate if the respective plant name corresponded to the plant name appearing in RBH. The information available in the references was critically evaluated, taking into account: i) whether the references citing the plant were distributed across only one, two or all three of the above categories; ii) cultural-historical aspects (e.g. the location or the time period from which the plant names were reported, based on information in the references used); iii) the distribution area of the candidate plant according to the *Flora Helvetica* (Lauber et al., 2018). Based on this, the respective candidates were rated regarding the plausibility of being the correct attribution (1 – high, 2 – moderate, 3 – small chance,– doubtful). For example, if references from all three categories (A, B, C) suggested the same candidate species for the respective RBH plant name and if that species was also distributed in the region, this was considered a highly possible attribution (score “1”). In general, candidate plants for RBH plant names reported from locations in Switzerland were considered more plausible than those reported for the same plant name from other neighbouring regions. Candidate plants that were only reported in references from one category or for which other uncertainties existed were considered doubtful (“?”).

We employed Kew’s Medicinal Plant Names Services (MPNS, 2023) to validate and harmonise the scientific nomenclature and taxonomy of all reported candidate species names. Scientific names not contained in MPNS were checked with Kew’s Plants of the World Online (POWO, 2023). For each RBH plant name a list of all suggested candidate plants was prepared, using their accepted scientific names. Any differing scientific names reported in the literature used were provided in parentheses after the accepted name.

#### 2.2.4 Analysis of the Hallwyl wound potion

Taking the transcribed text of the Hallwyl wound potion (**Supplementary Material, Document S1**), the recipe was analysed in the same way as the dermatological recipes recorded in the RBH database. The uses of the wound potion were interpreted according to section 2.2.2 and the plants used to prepare it assessed according to section 2.2.3.

### 2.3 Preliminary in vitro screening

#### 2.3.1 Selection of plants

For the preliminary bioactivity screening we selected plants from two resources:

1. Dermatological recipes in the RBH database (section 2.2.1) – Plants from this resource that fulfil the following three criteria were selected: i) RBH plant appears in recipes for the treatment of conditions thought to require medication with anti-inflammatory properties (among others); ii) RBH plant appears in recipes with only one active ingredient of plant origin (herbal simples). This allows to ascribe any potential bioactivity observed to one specific ingredient; iii) RBH plant name leads to candidate species (based on the procedure described in section 2.2.3) known to contain strongly bioactive compounds.
2. Transcription of the Hallwyl wound potion (section 2.2.4) – Only the plants used to prepare the basic remedy of the wound potion were selected.

#### 2.3.2 Plant collecting

Plants selected as described in section 2.3.1 were collected from wild populations in Switzerland between June and October 2022. As far as this was indicated in RBH, the collection was carried out at the recommended collection time and focused on the plant part mentioned in the recipes. Adherence to the local List of Protected Species (Federal Council of Switzerland, 2017) and the due diligence provisions according to the Nagoya Protocol in Switzerland was ensured in all cases (https://www.fedlex.admin.ch/eli/cc/2016/39/en#a8). All plant material was dried at 40°C immediately after collection and stored in air-tight containers until further use. Botanical voucher specimens (physical and/or digital) were collected from all candidate species and deposited in the Herbarium of Natural Product Research and Phytopharmacy Group at the ZHAW – Zurich University of Applied Sciences in Wädenswil (CH). Botanical identities were confirmed with the *Flora Helvetica* (Lauber et al., 2018) and each specimen was assigned a unique voucher number.

#### 2.3.3 Sample preparation

Different hydroethanolic crude extracts were prepared from the plant material collected. Due to the preliminary nature of the study, we were not able to also include pure aqueous extracts or such produced with solvents of lower polarity.

##### 2.3.3.1 Hydroethanolic macerates

Samples of hydroethanolic macerates were prepared from all plant materials collected as well as from the mixture of the plant materials used as ingredients in the Hallwyl wound potion (section 2.3.3.2). Dried plant material was ground using an analytical grinder (IKA, Staufen, Germany) and added to the tenfold amount of 80% ethanol (m/m). The mixture was stirred at room temperature for two hours and filtered. Ethanol was removed from the filtrate by rotary evaporation at 40°C and 50 mBar (Büchi, Flawil, Switzerland). The remaining aqueous liquid was frozen at -20°C and lyophilized using an Alpha 2-4 LSCPlus device (Martin Christ, Osterode am Harz, Germany). The dried extract was ground to a fine powder and stored at -20°C.

##### 2.3.3.2 Hallwyl wound potion

According to the text of the Hallwyl wound potion (**Supplementary Material, Document S1**), the herbal ingredients of the basic remedy, dried, powdered and in equal parts, shall be boiled together in wine for as long as it takes “to hard-boil an egg”. In an approximation of the historical preparation, first a hydroethanolic decoction was prepared: the mixture of equal parts of the dried powdered herbs was added to the tenfold amount of 10% (m/m) of ethanol and then simmered for 10 minutes. In addition to the decoction, the Hallwyl wound potion was also prepared as a hydroethanolic macerate following the procedure described in section 2.3.3.1.

##### 2.3.3.3 Pure compounds and positive control

To compare possible effects of the extract samples with the activity of pure plant compounds, corresponding substances with reported anti-inflammatory effects in cell-based *in vitro* models were also tested: silymarin, quercetin, imperatorin, glycyrrhetinic acid (Chang et al., 2013; Gugliandolo et al., 2020; Guo et al., 2012; Wang et al., 2011). Dexamethasone was used as the positive control.

#### 2.3.4 Stock solutions

Extract samples were dissolved using pure DMSO, or 10% (v/v) DMSO in PBS in samples prepared by hydroethanolic decoction (10% ethanol (m/m)), at a concentration of 100 mg/mL. Pure compounds and control were dissolved in pure DMSO at a concentration of 10 mM. Stock solutions were stored at -20°C. Before use, stock solutions were diluted with PBS to the appropriate concentration and filtrated using a 0.22 μm PES filter. No precipitation was observed upon dilution with PBS for any of the samples.

#### 2.3.5 MTT Assay

Cytotoxic effects of all samples were determined using the MTT-assay in the NIH-3T3 murine fibroblast cell line (DSMZ, ACC59). The cells were cultured in DMEM-HG (BioConcept, Allschwil, Switzerland) with 10% heat inactivated FBS (Fischer Scientific AG, Reinach, Switzerland) by volume at 37°C and 5% CO_2_. Cells were only used until the 15^th^ passage.

Cultured cells were trypsinized and 100 µL of the suspension was seeded into flat-bottom 96-well plates at a density of 1 × 10^4^ cells/well and subsequently incubated for 24h. To each well, 10 μL of diluted sample was added and the treated plates were again incubated for 24h. Afterwards, the medium was aspirated and exchanged for 100 µl of DMEM-HG containing MTT (Sigma-Aldrich, St. Louis, MO, USA) at a concentration of 0.5 mg/mL. The plates were then incubated for further 4h. Medium was gently removed and 200 µl of DMSO added to dissolve the precipitate, aided by a plate shaker. Finally, 25 µl of Sorensen’s Glycine Buffer (0.1 M NaCl, 0.1 M glycine, 0.1 M NaOH) was added into each well. Absorbance was measured at 640 and 570 nm using a Spark 20M plate reader (Tecan, Männedorf, Switzerland). The 640 nm measurements served as the reference and were subtracted from the individual 570 nm measurements.

#### 2.3.6 Whole blood assay and PBMC assay

Following largely the procedure described in Elisia et al. (2018), a whole blood assay was conducted with the aim to assess the potential of the test samples to modulate the release of the inflammatory mediators IL-6 and TNF.

Anonymised blood samples collected from healthy volunteers, were obtained from Blutspende Zürich (https://www.blutspendezurich.ch/) in heparin coated tubes and stored at 4°C for a maximum of 24 hours before use. The procedure was confirmed by the Ethics Committee of the Canton of Zurich (BASEC Nr. Req-2022-0139).

Samples were added to the wells of a flat-bottom 96-well plate at a volume of 10 μL. Subsequently, 50 μL of human whole blood were added into each well. The plate was then pre-incubated at 37°C and 5% CO_2_ for one hour. Afterwards 10 μL of PBS as control or PBS containing LPS (Sigma-Aldrich, St. Louis, MO, USA) were added, resulting in a final concentration of 10 ng/mL of LPS per well. The plate was then incubated for six hours at 37°C and 5% CO_2_. Supernatants were removed after centrifugation at 400g and 4°C for 5 minutes and stored at -80°C until further use.

As a confirmatory study of the whole blood assay, PBMCs were isolated from whole blood via density-gradient centrifugation using Histopaque-1077 according to manufacturer’s instruction. PBMCs were maintained in RPMI-1640 medium (BioConcept, Allschwil, Switzerland) supplemented with 10% FBS and 50 U/mL of penicillin and streptomycin, each at 37°C and 5% CO_2_. Subsequently cells were seeded into round-bottom ultra-low attachment 96-well plates at 5 × 10^5^ cells/mL. The plates were incubated for 14 hours at 37°C and 5% CO_2_ and then 10 μL of diluted samples were added. After 1 hour of incubation LPS was added to a final concentration of 10 ng/mL. After 6 hours supernatants were removed and stored at -80°C until further use.

#### 2.3.7 Haemolytic activity

To determine the haemolytic activity of the extract samples, the optical density of all supernatants taken from the whole blood assay was measured at 540 nm using a plate reader. Triton X-100 (1 %) was added instead of LPS in PBS to act as haemolysis positive control.

#### 2.3.8 ELISA

Determination of the cytokine levels in the supernatants was carried out using commercial ELISA kits for human IL-6 and TNF (BioLegend, San Diego, US). ELISA was carried out according to the manufacturer’s instructions.

#### 2.3.9 Data analysis

All experiments were carried out with n = 3. When human primary cells were used, replicates corresponded to individual donors. Statistical tests were done using GraphPad Prism version 9.5.0 for Windows (GraphPad Software, Boston, USA).

## 3 Results

### 3.1 Pharmaco-botanical analysis of the historical text

#### 3.1.1 Dermatological recipes in the RBH database

In Fankhauser’s (2012) transcription (see section 2.1) of RBH we were able to distinguish 196 recipes linked with dermatological uses that contained at least one plant ingredient. All 196 recipes were recorded in the RBH database.

Each recipe provides information about the symptom or disease treated, the ingredients to be used, the method of preparation and how to apply the medicine. Most of the recipes deal with the preparation of medicines for external use (182 recipes). However, there are 14 recipes in which some kind of drink is prepared that is taken orally but is nevertheless aimed at the treatment of some topically manifest symptom. Sometimes these preparations are addressed as “trunck” or “tranck” (potion) and often prepared in a similar way to the famous Hallwyl wound potion (“das recht Hallwylisch wundtrannck”), in which the pounded herbs were boiled in wine.

Analysis of our RBH database reveals that the preparations most frequently mentioned in the 196 recipes are ointments (53 recipes), different kinds of so-called “waters” which also include aqueous or alcoholic distillates (31), decoctions in water, wine or vinegar (22), and cataplasms (22). In 19 recipes the preparation used is not clearly stated.

Depending on the list of ingredients, the medicines prepared in the recipes can be classified into compound recipes containing either, plant and non-plant ingredients (83 recipes), or only plant ingredients (36), as well as herbal simples, i.e., preparations that contained only one plant ingredient (77). Liquids used as extraction solvents (e.g. water, wine, vinegar) or different kinds of flour used in plasters were regarded as technical ingredients and not considered in this classification (**Supplementary Material – Table S1)**.

In consideration of the focus of this paper, we limit further analysis of the dermatological recipes in RBH to the plants mentioned therein and their uses.

#### 3.1.2 Medicinal plant uses in the RBH database

Combining the mentioned RBH uses referring to the same medical complaint (see section 2.2.1) allows us to distinguish 52 different medicinal uses that are mentioned in the 196 recipes. For each of the combined uses one or more interpretations are available. In four cases we were not able to provide a more detailed interpretation than that the mentioned uses seem to refer to some skin disorder. The cross-check of the interpreted uses with modern diagnostic tools (see section 2.2.1) reveals that most of the uses (47) are related to poorly healing or infected wounds, inflammations of the skin, burns, infectious diseases of bacterial, fungal, mycobacterial, parasitic or viral origin, as well as neoplastic skin chances. A small number of uses (5) deals with stopping of acute bleeding or dissolving haematomas. In most of the underlying medical conditions possible pharmacological interventions are largely based on medication with antibiotic, anti-inflammatory or immunomodulatory properties (**Table 1**).

**Table 1.**
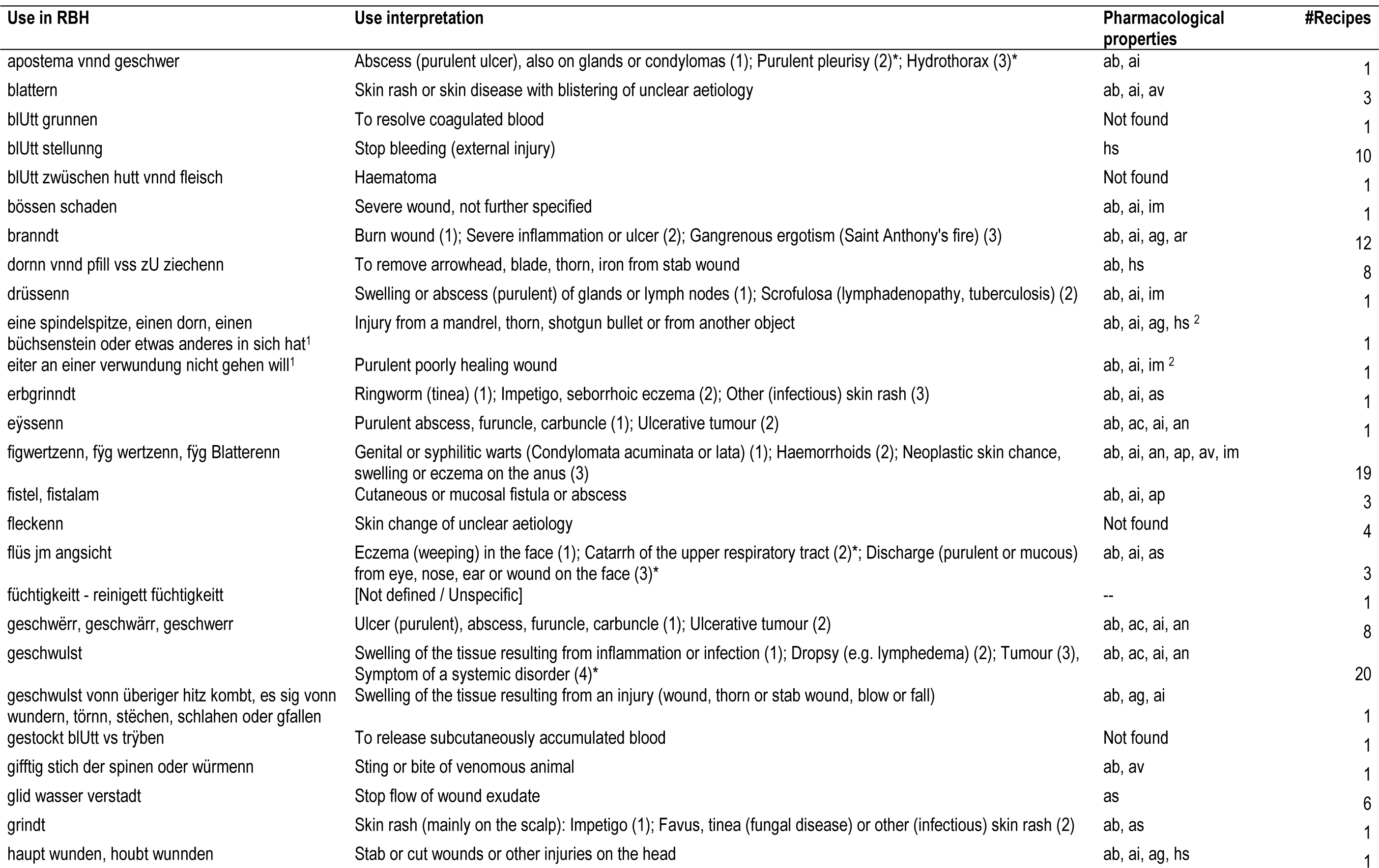
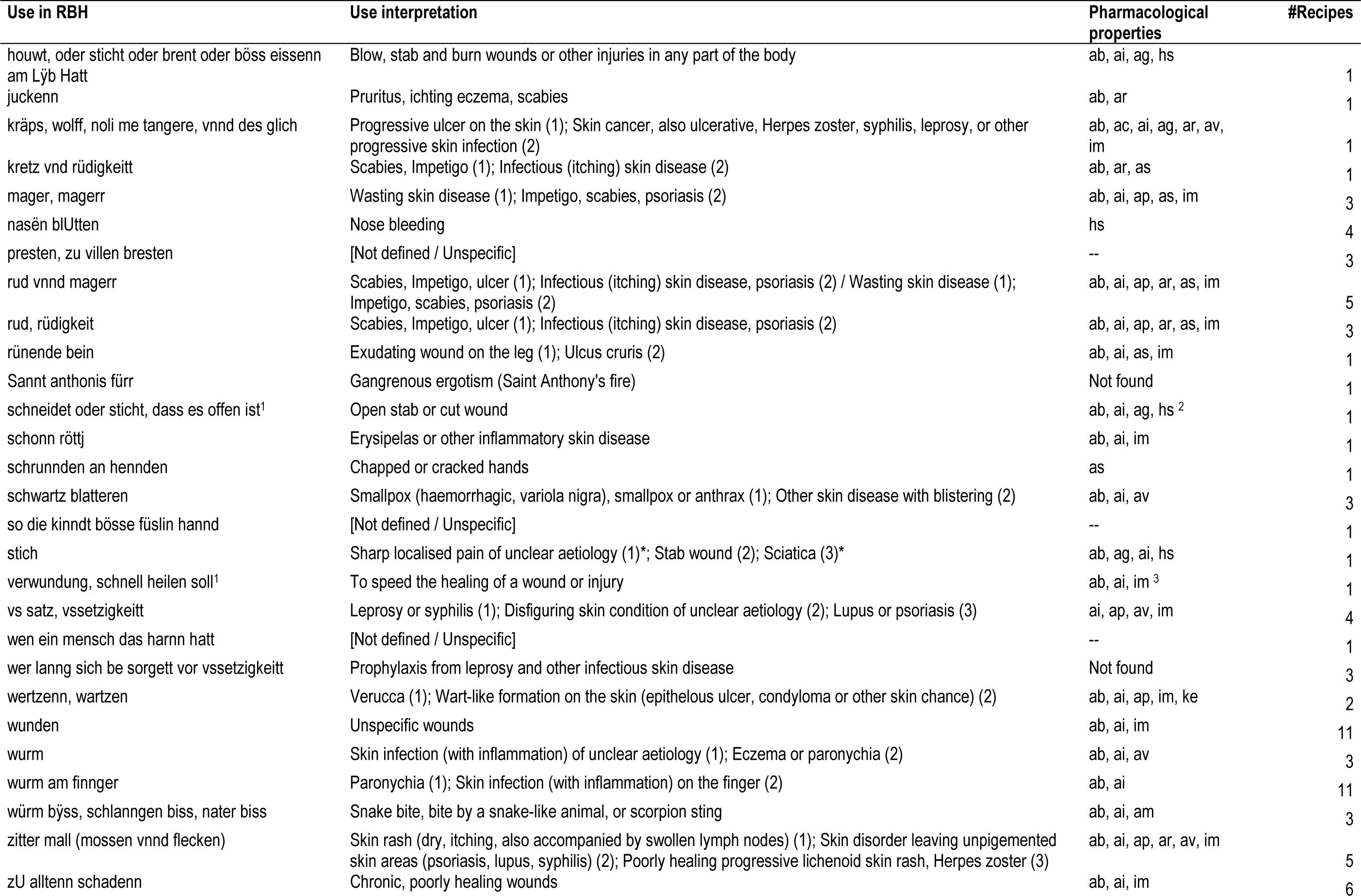
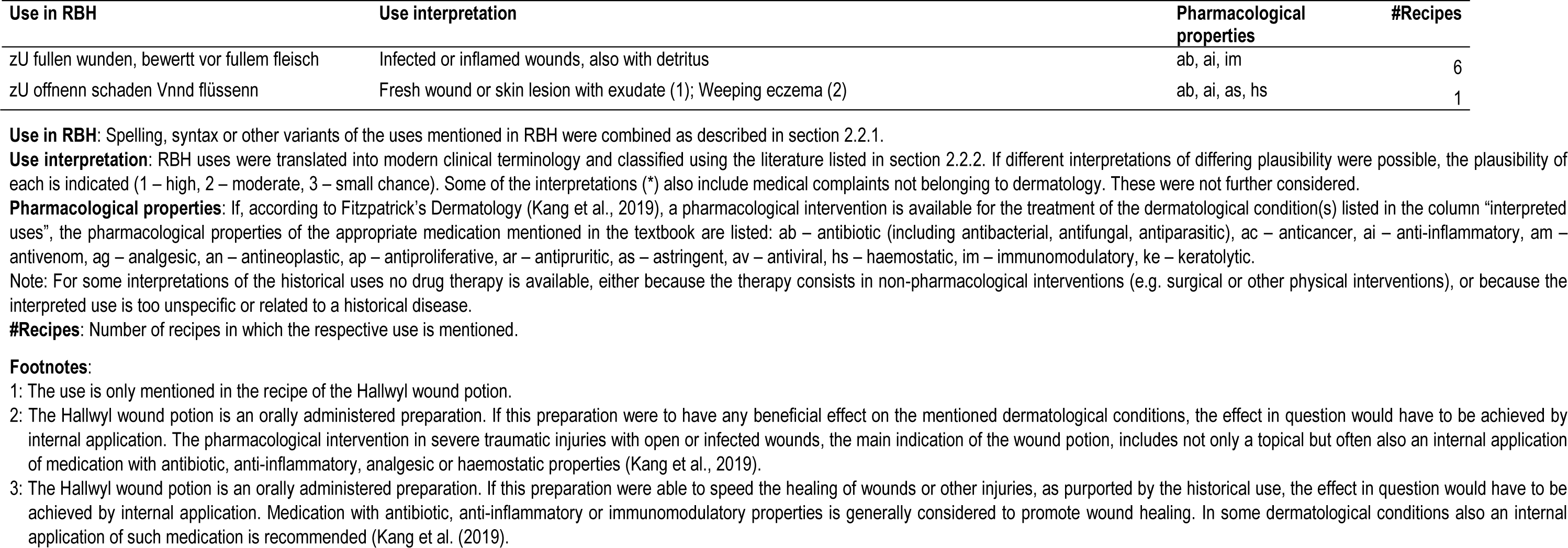
Interpretation of the medicinal uses mentioned in the investigated dermatological recipes of the *Receptarium* of Burkhard from Hallwyl (RBH).

The most frequently mentioned uses in the 196 recipes refer to different kinds of wounds, such as “wunden” (11 recipes), “zU alltenn schadenn” (6), “zU fullen wunden” (6), mentioned in altogether 23 recipes and which were interpreted as unspecific or infected and inflamed wounds, as well as chronic or poorly healing wounds. Further important uses are: “gschwulst” – swelling of the tissue as a result of inflammation, tumour, dropsy or an infectious disease (20 recipes); “figwertzen” – genital warts (condylomas) or perhaps haemorrhoids, cancerous skin change or ulcers on the anus (19); “branndt” – burn wound, violent inflammation of unclear aetiology, or gangrenous ergotism (Saint Anthony’s fire) (12); “wurm am finger” – paronychia or another skin inflammation on the finger (11); “blutt stellunng” – stop bleeding in the event of an external injury (10); “dornn vnnd pfill vss zU ziechenn“ – remove an arrowhead, blade, thorn or another iron from a stab wound (8); “geschwërr, geschwärr” – ulcer (purulent), abscess, furuncle, carbuncle, or ulcerative tumour (8) (**Table 1**).

#### 3.1.3 Plants in the RBH database

In the 196 recipes we counted 159 plant names, representing either generic names of plants, dedicated names of plant substances (e.g. “mastix”, “olibanum”, “terpenntin”), spices (e.g. “macis”, “saffrann”, “zimett”) or products of plant origin such as “baum öll”.

For each of the 159 plant names a list of the potential botanical identities mentioned in the literature (candidate plants) had been prepared as described in section 2.2.3. In each case, the information available in the literature was critically reviewed to establish the attribution of a particular scientific name (**Supplementary Material, Tables S2 and S3**). The botanical identity of 4 plant names could not be established based on the literature used. For the remaining 155 plant names one or more candidate plants were available (**Table 2**).

**Table 2.**
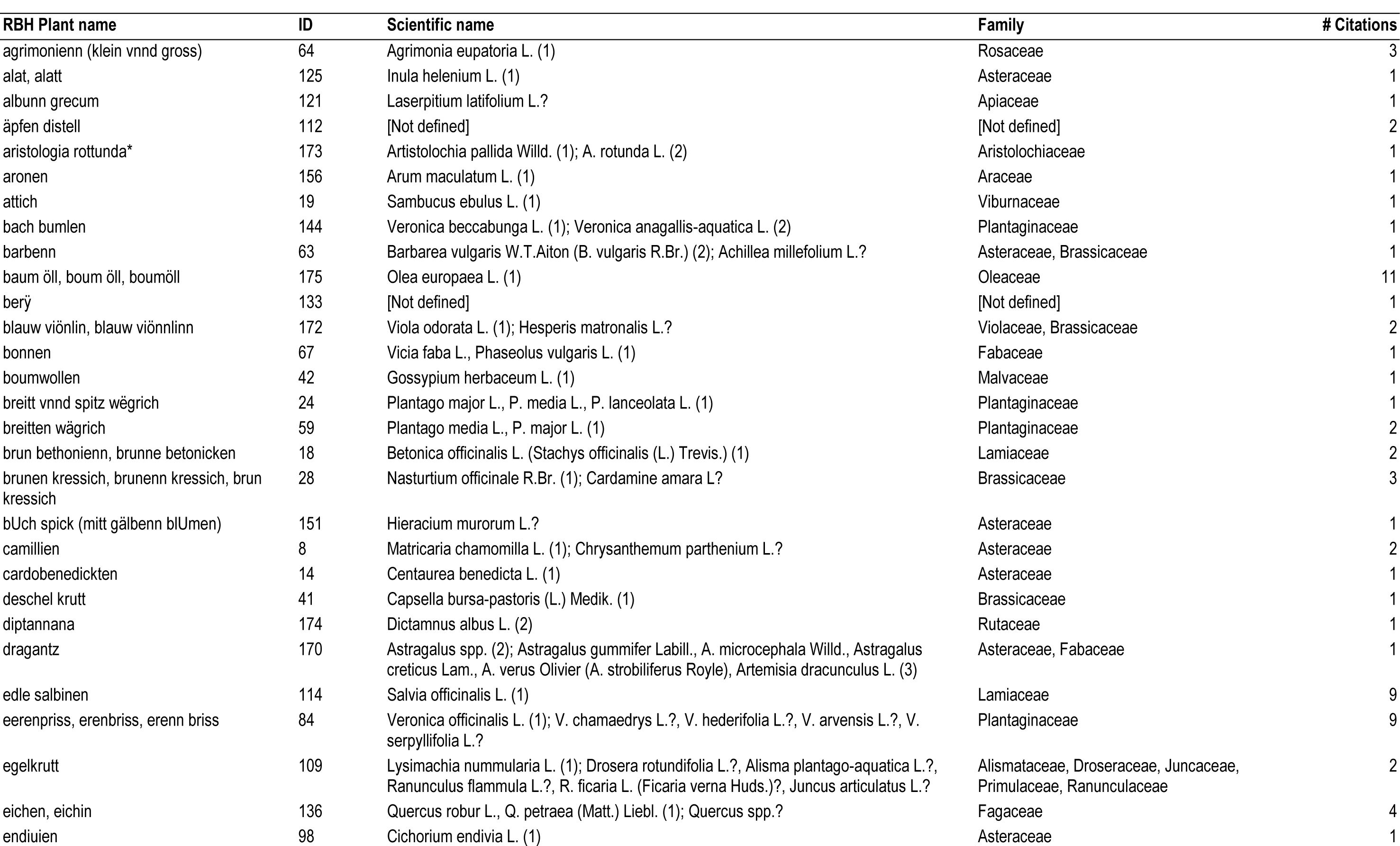
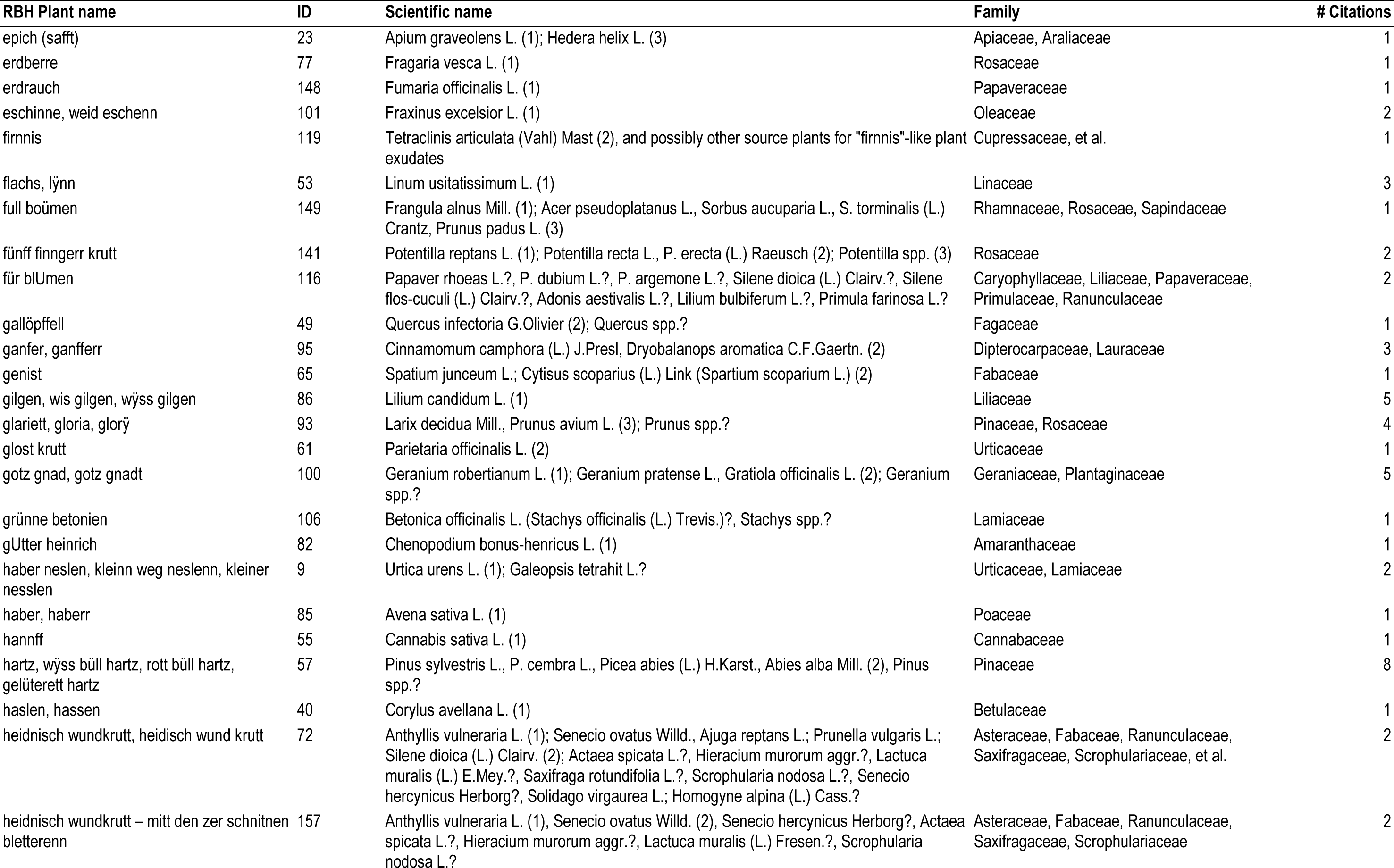
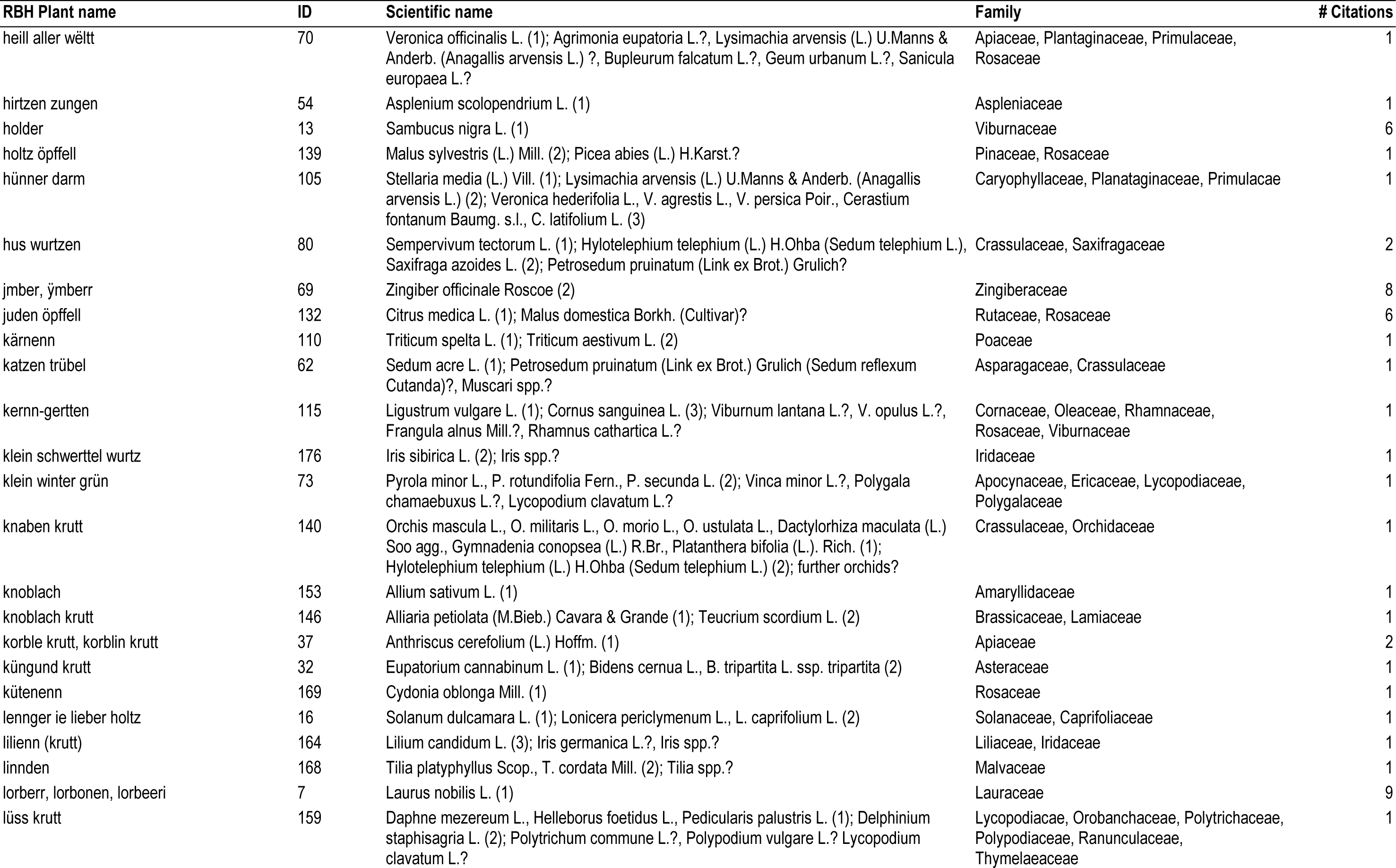
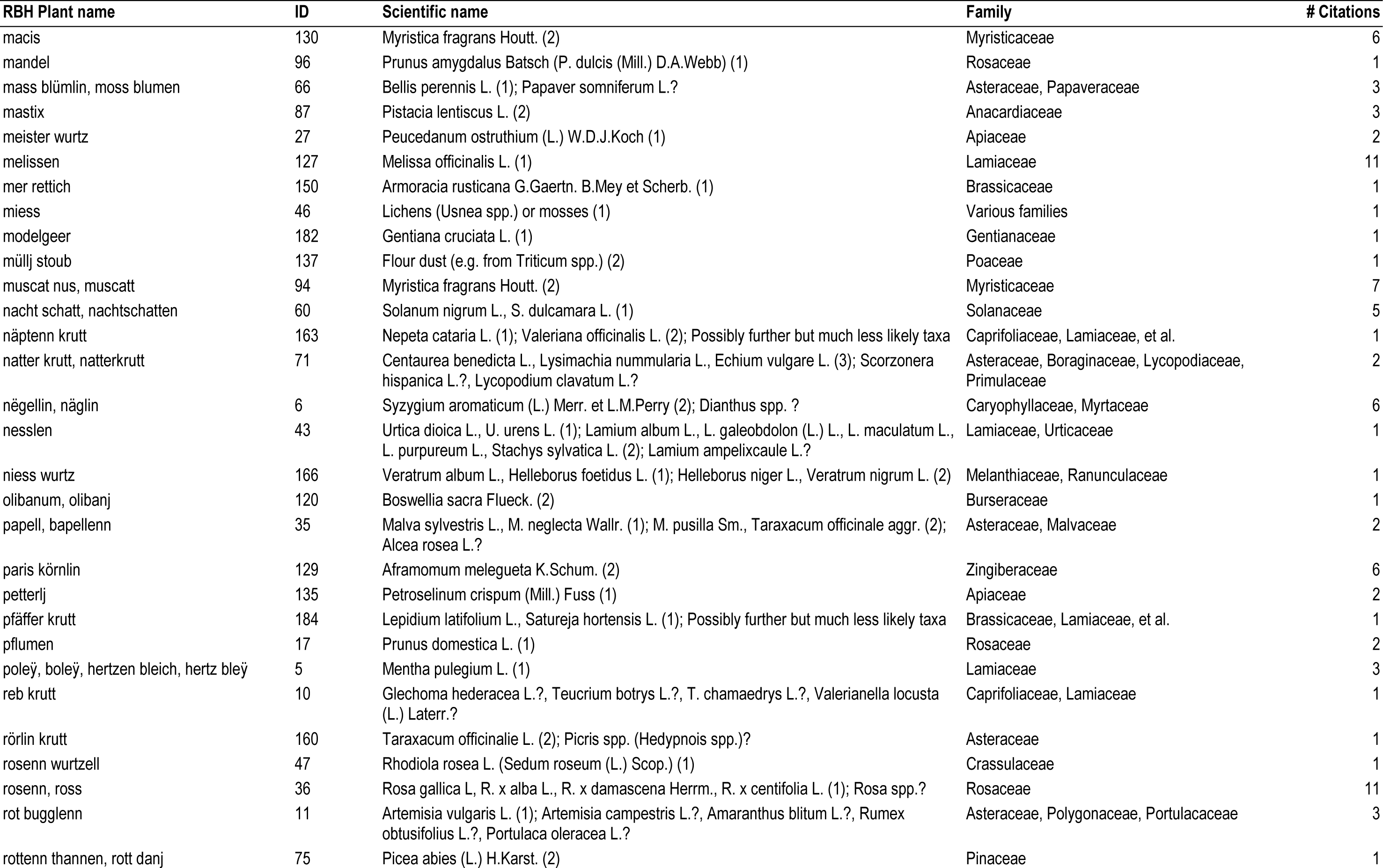
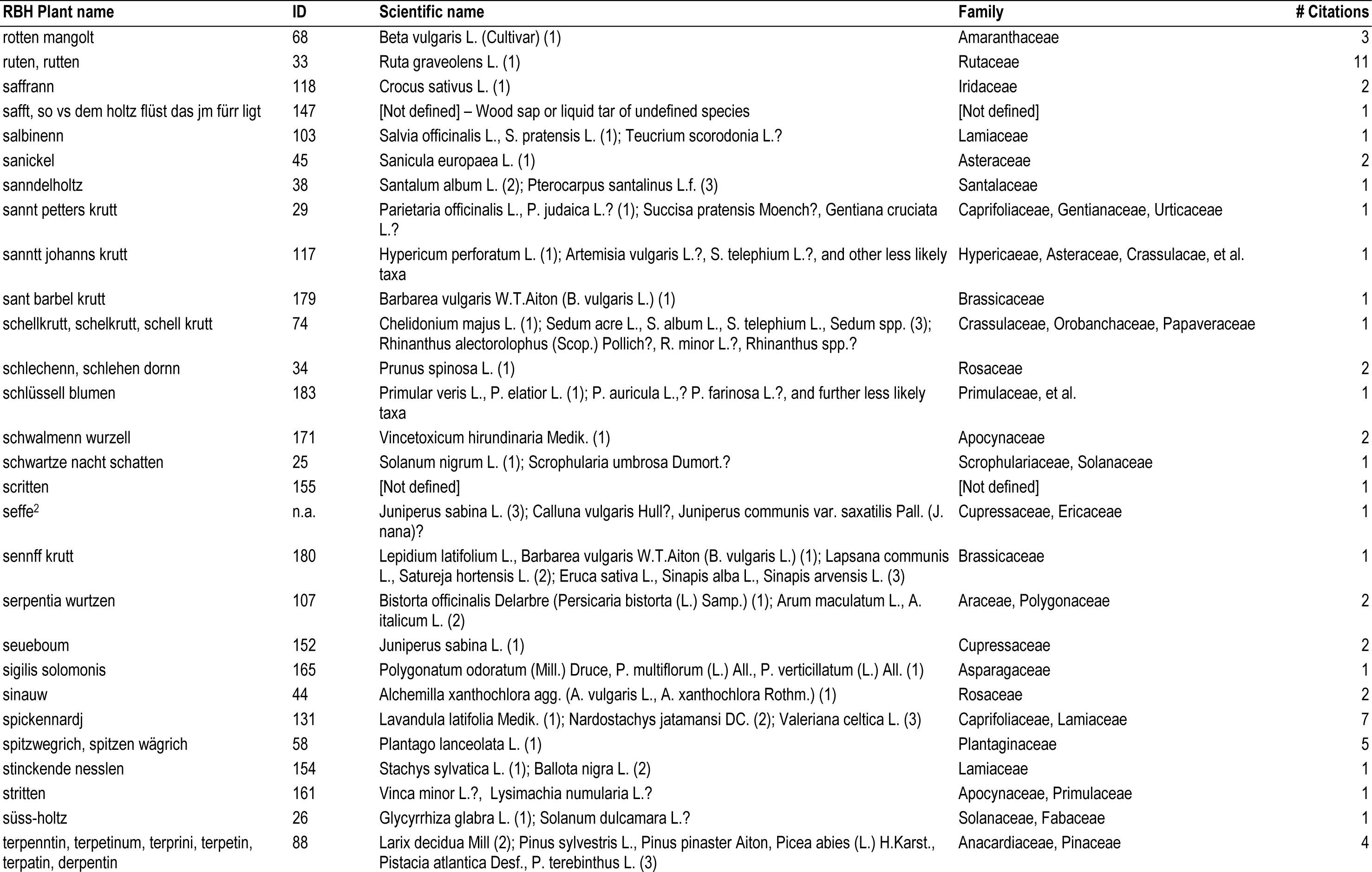
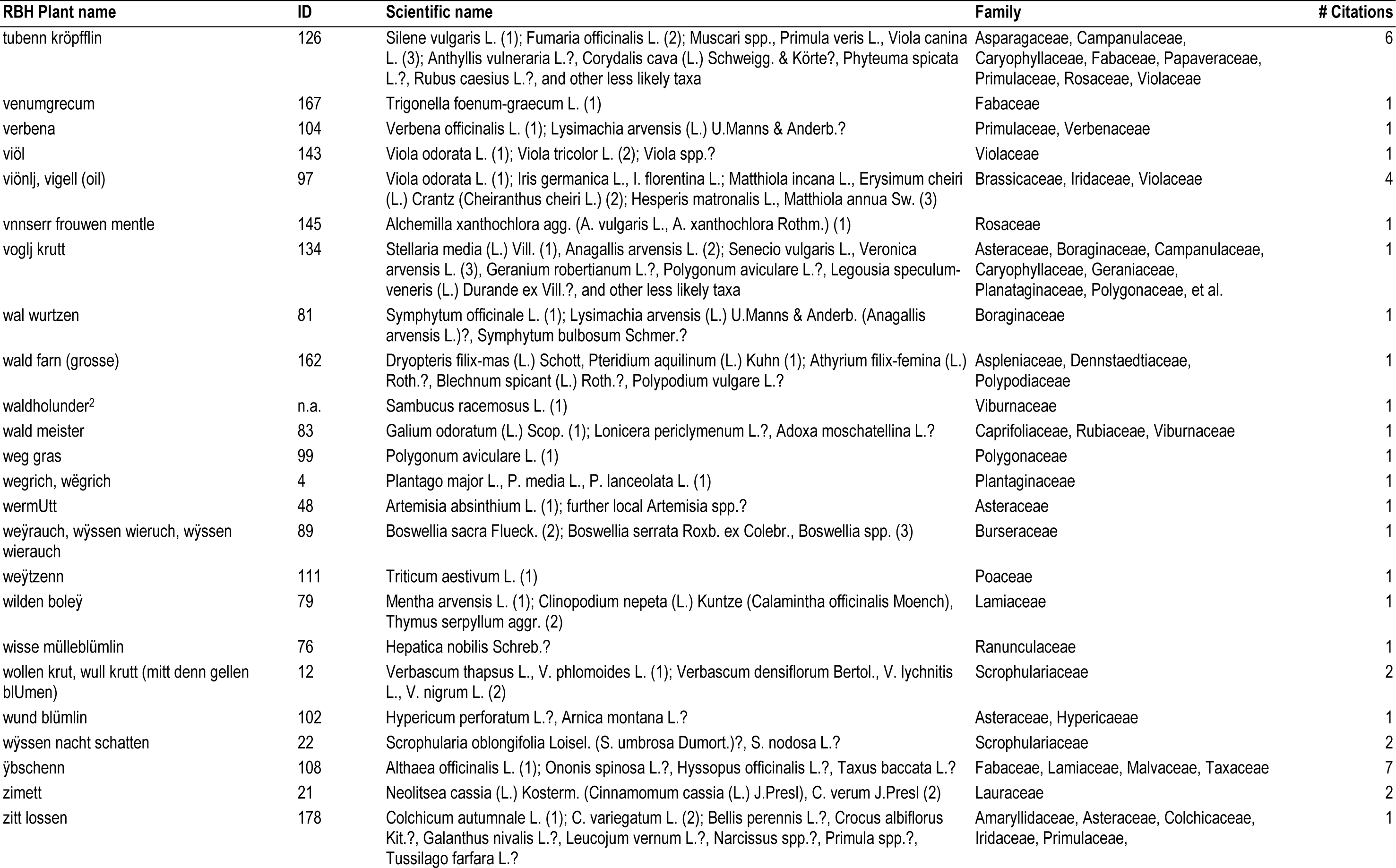
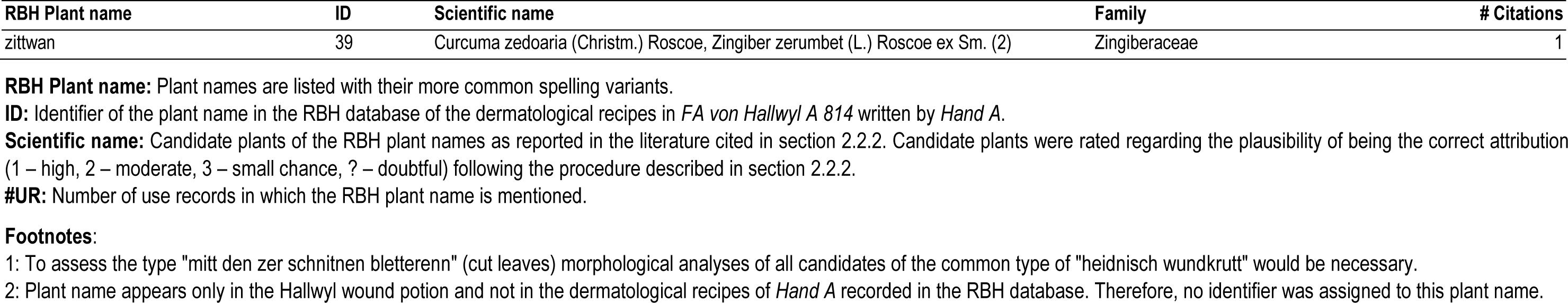
Plant names in the investigated dermatological recipes of the *Receptarium* of Burkhard from Hallwyl (RBH) and the associated candidate species.

The results show that 53 (34%) of these 155 plant names lead to only one single candidate species with the highest chance of being the correct attribution (score “1”). In these cases, at least one reference from each of the three categories comes to the same conclusion, with no references having divergent views. The botanical identity is also supported when assessed according to the criteria plant distribution and cultural-historical context. In all other cases, the association between plant name and candidate species is less consistent. For the majority of the assessed RBH plant names (102 cases, 66%) two or more candidate species are available, often from different genera or families. In numerous of these cases the assignment remains doubtful (cases marked with “?”) (**Table 2**), because information from only one category of references was available or because of other uncertainties (**Supplementary Material, Tables S2 and S3**).

Further analysis of the 159 RBH plant names reveals that they are involved in 362 unique plant use records (URs) (**Supplementary Material, Table S4**). Among the most frequently mentioned plant names are: “melissen” consistently attributed to *Melissa officinalis* and in RBH mainly used in form of liquid preparations (11 URs); “rosenn” or “ross” – garden roses from different *Rosa* species used for the preparation of rose oil, rose water or vinegar (11 URs); “ruten” – *Ruta graveolens* (11 URs); “edle salbinen” – *Salvia officinalis* (9 URs); “eerenpriss” – *Veronica officinalis* and perhaps other *Veronica* species, also mainly used in form liquid preparations (9 URs); “lorberr” – *Laurus nobilis*, with laurel oil or fruits (9 URs); Different qualities of “hartz” – resins from conifers from the genera *Abies*, *Picea* and *Pinus* (8 URs); “jmber” or “ÿmberr” – *Zingiber officinale* (8 URs). Another frequently mentioned herbal ingredient is “boum öll”, olive oil from *Olea europaea* appearing in 11 URs. However, it is not always clear to what extent this is an active component or rather a technical aid or carrier, as for example in its use as an extraction solvent in the preparation of oily macerates (**Table 2**).

#### 3.1.4 Plants and their uses in the Hallwyl wound potion

Text analysis of the Hallwyl wound potion reveals that one basic remedy and three additional variants, each with one supplementary plant, are described (**Supplementary Material, Table S5**). According to our counting method (see section 2.2.1) this corresponds to four recipes with four individual uses, all of which are not mentioned in the recipes recorded in the RBH database (section 3.1.2). The mentioned uses relate to the treatment of traumatic wounds, including purulent, poorly healing wounds or other skin injuries. Possible pharmacological interventions rely on medication with antibiotic, anti-inflammatory, analgesic, immunomodulatory or haemostatic properties (**Table 1**, see footnotes 1-3).

The text gives a detailed description of preparing a decoction with white wine using the plants mentioned. As explained in section 2.3.1, six plants are used for the preparation of the basic remedy. Of the three additional plants used for the preparation of the variants of the remedy, two – “seffe” and “waldholunder” – are not mentioned in recipes recorded in the RBH database and thus only appear here in the wound potion (**Table 2**, see footnote 2).

Including data from the Hallwyl wound potion increases the total number of recipes analysed in this study to 200, plant names to 161 and medicinal uses to 56.

### 3.2 Results of the preliminary *in vitro* screening

#### 3.2.1 Selection of plants and test samples

We collected 13 different plant materials from 11 candidate species identified by applying the selection criteria described in section 2.3.1 (**Table 3**). Four species originate from the RBH database and seven species from the Hallwyl wound potion:

**Table 3.**
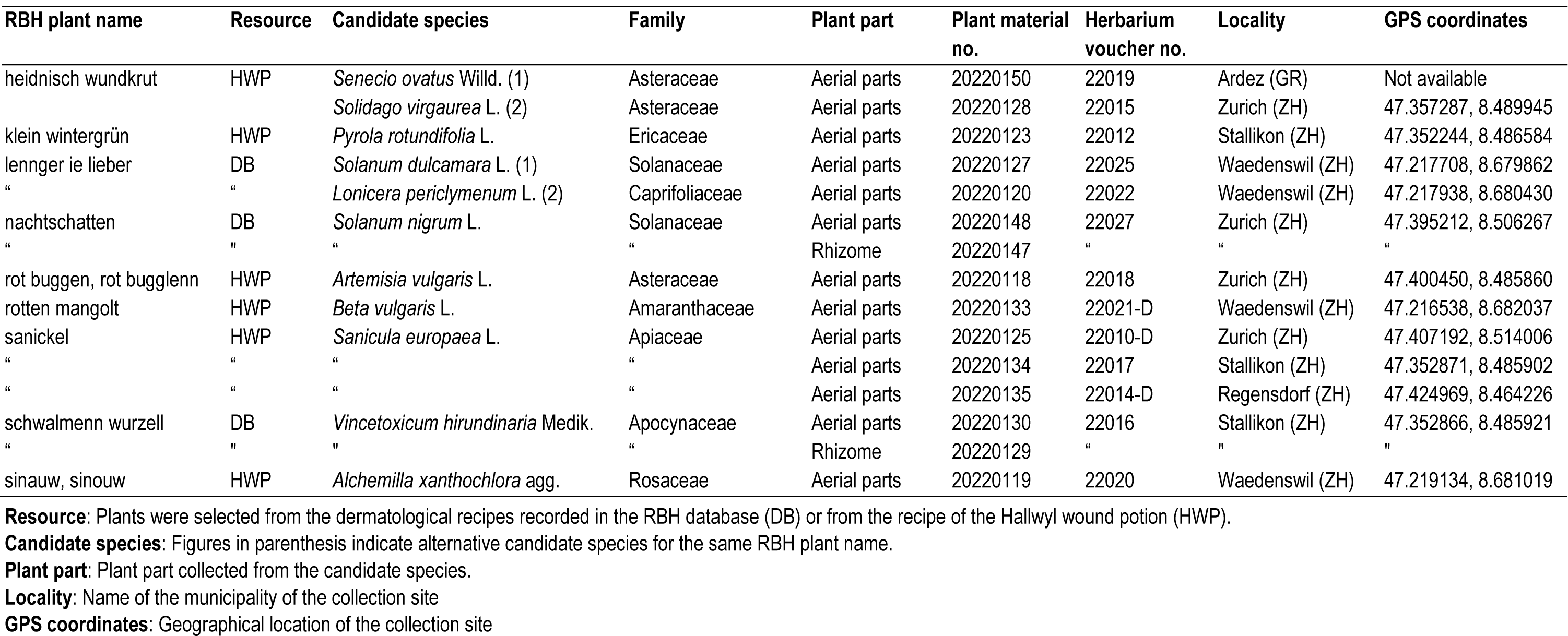
Details of the plant material collected from the 11 candidate species for preliminary in vitro screening.

| RBH plant name | Resource | Candidate species | Family | Plant part | Plant material no. | Herbarium voucher no. | Locality | GPS coordinates |
| --- | --- | --- | --- | --- | --- | --- | --- | --- |
| heidnisch wundkrut | HWP | *Senecio ovatus* Willd. (1) | Asteraceae | Aerial parts | 20220150 | 22019 | Ardez (GR) | Not available |
|  |  | *Solidago virgaurea* L. (2) | Asteraceae | Aerial parts | 20220128 | 22015 | Zurich (ZH) | 47.357287, 8.489945 |
| klein wintergrün | HWP | *Pyrola rotundifolia* L. | Ericaceae | Aerial parts | 20220123 | 22012 | Stallikon (ZH) | 47.352244, 8.486584 |
| lennger ie lieber | DB | *Solanum dulcamara* L. (1) | Solanaceae | Aerial parts | 20220127 | 22025 | Waedenswil (ZH) | 47.217708, 8.679862 |
| " | " | *Lonicera periclymenum* L. (2) | Caprifoliaceae | Aerial parts | 20220120 | 22022 | Waedenswil (ZH) | 47.217938, 8.680430 |
| nachtschatten | DB | *Solanum nigrum* L. | Solanaceae | Aerial parts | 20220148 | 22027 | Zurich (ZH) | 47.395212, 8.506267 |
| " | " | " | " | Rhizome | 20220147 | " | " | " |
| rot buggen, rot bugglenn | HWP | *Artemisia vulgaris* L. | Asteraceae | Aerial parts | 20220118 | 22018 | Zurich (ZH) | 47.400450, 8.485860 |
| rotten mangolt | HWP | *Beta vulgaris* L. | Amaranthaceae | Aerial parts | 20220133 | 22021-D | Waedenswil (ZH) | 47.216538, 8.682037 |
| sanickel | HWP | *Sanicula europaea* L. | Apiaceae | Aerial parts | 20220125 | 22010-D | Zurich (ZH) | 47.407192, 8.514006 |
| " | " | " | " | Aerial parts | 20220134 | 22017 | Stallikon (ZH) | 47.352871, 8.485902 |
| " | " | " | " | Aerial parts | 20220135 | 22014-D | Regensdorf (ZH) | 47.424969, 8.464226 |
| schwalmenn wurzell | DB | *Vincetoxicum hirundinaria* Medik. | Apocynaceae | Aerial parts | 20220130 | 22016 | Stallikon (ZH) | 47.352866, 8.485921 |
| " | " | " | " | Rhizome | 20220129 | " | " | " |
| sinauw, sinouw | HWP | *Alchemilla xanthochlora* agg. | Rosaceae | Aerial parts | 20220119 | 22020 | Waedenswil (ZH) | 47.219134, 8.681019 |

**Resource:** Plants were selected from the dermatological recipes recorded in the RBH database (DB) or from the recipe of the Hallwyl wound potion (HWP).
**Candidate species:** Figures in parenthesis indicate alternative candidate species for the same RBH plant name.
**Plant part:** Plant part collected from the candidate species.
**Locality:** Name of the municipality of the collection site
**GPS coordinates:** Geographical location of the collection site

In the dermatological recipes recorded in the RBH database we identified three RBH plants (“lennger ie lieber”, “nachtschatten”, “schwalmenn wurzell”) mentioned in four recipes leading to candidate species from the Apocynaceae and Solanaceae which best met the selection criteria described in section 2.3.1 (**Supplementary Material, Table S6**). From these recipes, altogether four candidate species were selected: *Solanum dulcamara*, *Solanum nigrum*, *Vincetoxicum hirundinaria* and *Lonicera periclymenum*. The latter species of the Caprifoliaceae was also included because it is one of two secondary alternatives (candidates with score “2”) for “lennger ie lieber”, which in the first place leads to *Solanum dulcamara* (score “1”) (**Table 3**).

From the transcription of the Hallwyl wound potion we selected the six plants used to prepare the basic remedy of the herbal compound preparation: “rotten mangolt”, “heydnisch wundkrut”, “klein wintergrün”, “rot buggen”, “sinouw”, “sanickel” (**Supplementary Material, Table S5**). Following the procedure described in section 2.2.3, the six RBH plant names lead to more than six candidate species, because in two cases (“heidnisch wundkrutt” and “klein winter grün”) more than one candidate is possible. Focusing on the most plausible and available candidates, altogether seven different species were selected (**Table 3**).

From the 13 plant materials consisting of aerial parts and roots 17 different samples of hydroethanolic crude extracts were prepared (**Table 4**). Four of the samples represent different reconstructions of the Hallwyl wound potion – two macerates (samples 4001, 4002) and, in approximation of the historical recipe, two decoctions (samples 4003, 4004). In the case of both, macerate and decoction, one sample each was prepared with one of two alternatives for the RBH plant “heidnisch wundkrutt” – *Senecio ovatus* or *Solidago virgaurea*.

**Table 4.**
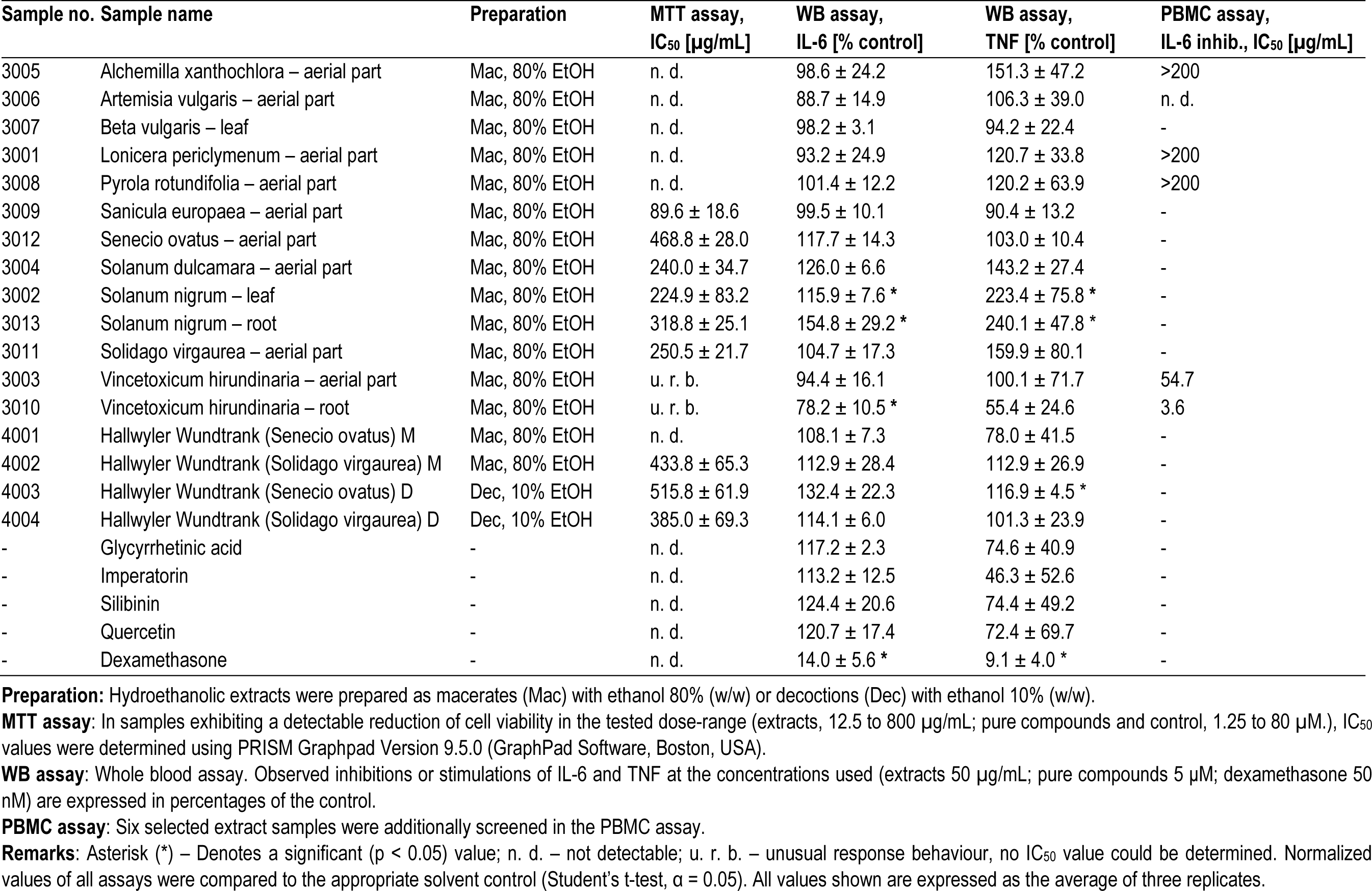
Test samples and assay results. Cytotoxic effects in the MTT assay and immunomodulatory effects in the LPS-stimulated whole blood assay and PBMC assay.

#### 3.2.2 Cytotoxic effects

Effects of the samples on the ability of NIH-3T3 cells to metabolize MTT into its purple formazan salt were equated with the general cell viability and regarded as a measure of any potentially relevant cytotoxic effect. Extract samples were tested at 800, 200, 50 and 12.5 μg/mL, pure compounds and control at 80, 20, 5, 1.25 μM (**Table 4**).

Values of IC_50_ under 100 μg/mL were considered as cytotoxic and corresponding samples excluded from further investigation. *Sanicula europaea* (sample 3009) is the only such candidate with a calculated IC_50_ of 89.55 ± 18.64 μg/mL. Extracts of both aerial parts and roots of *Vincetoxicum hirundinaria* (samples 3003, 3010) showed uniform suppressive effects on cell viability over the tested dose range. These unusual dose-response effects made curve fitting unfeasible and no IC_50_ values could be determined. Therefore, a second experiment with eight instead of four dose points was carried out, revealing a constant reduction in cell viability to around 50% across all doses from 6.3 to 1600 μg/mL (**Supplementary Material, Figure S2**). All other extract samples as well as pure compounds and control either did not exhibit a detectable reduction of cell viability or did so only with an IC_50_ above our stated threshold (100 μg/mL) (**Table 4**).

#### 3.2.3 Modulation of the inflammatory mediators IL-6 and TNF

Whole blood assay was used to screen all samples for their potential to modulate the release of the inflammatory mediators IL-6 and TNF after stimulation with LPS Extract samples were screened first directly in whole blood, at a concentration of 50 μg/mL, pure compounds at 5 μM and dexamethasone at 50 nM. Normalized values were compared to the appropriate vehicle control.

Of all the extract samples, only *Vincetoxicum hirundinaria* root (hydroethanolic macerate) (sample3010) shows a significant inhibitory effect with a reduction of LPS induced IL-6 release in whole blood to 78.1% compared to vehicle control. On the other hand, significant stimulatory effects can be observed with *Solanum nigrum* leaves and roots (sample 3002, 115.9%; sample 3013, 154.8%). All other extracts show varying degrees of inhibition or stimulation on LPS-induced IL-6 release, albeit with no significant effects.

The results on the modulation of TNF are largely analogous to those of IL-6. Again, the root of *V. hirundinaria* (sample 3010) shows the most prominent reduction of LPS induced release (55.4%), whereas *Solanum nigrum* leaves and roots (samples 3002, 3013) show significant stimulatory effects of over 100 % compared to vehicle control.

As expected, dexamethasone shows strong and significant inhibitory effects, suppressing IL-6 release almost completely. However, none of the pure compounds exhibit a significant effect, although a weak stimulation of IL-6 and a weak to moderate inhibition of TNF could be observed (**Table 4**). Here and elsewhere, comparatively large standard deviations in the measurement of TNF were observed in many samples, indicating larger variation of response in the donor blood samples (see section 2.3.5). No haemolytic activity beyond the threshold value of 5% compared to control (1% Triton X-100) was found for any of the extract samples tested in the whole blood assay.

Six selected extract samples were additionally screened in the PBMC assay with the aim to confirm the effects observed in the whole blood assay and to establish dose-response curves (**Supplementary Material, Figure S3**). TNF was not measured, because effects were expected to be analogous to IL-6, as previously observed in the whole blood assay. Samples were tested at the concentrations 200, 50, 12.5, 3.1, 0.8 and 0.2 μg/mL. Vehicle controls were also tested at the relevant concentrations. Both plant parts of *Vincetoxicum hirundinaria* show a distinct and significant inhibition of LPS-induced IL-6 release (samples 3003, 3010). Especially the root, with an IC_50_ value of 3.62 ± 0.54 μg/mL, reveals a remarkably strong effect (**Figure 1**). In all other samples, IC_50_ values are above 200 μg/mL or cannot be calculated (out of range value) as in the case of *Artemisia vulgaris* (sample 3006), despite the plant’s observed tendency of an inhibitory activity in the whole blood assay (**Table 4**).

**Figure 1.**
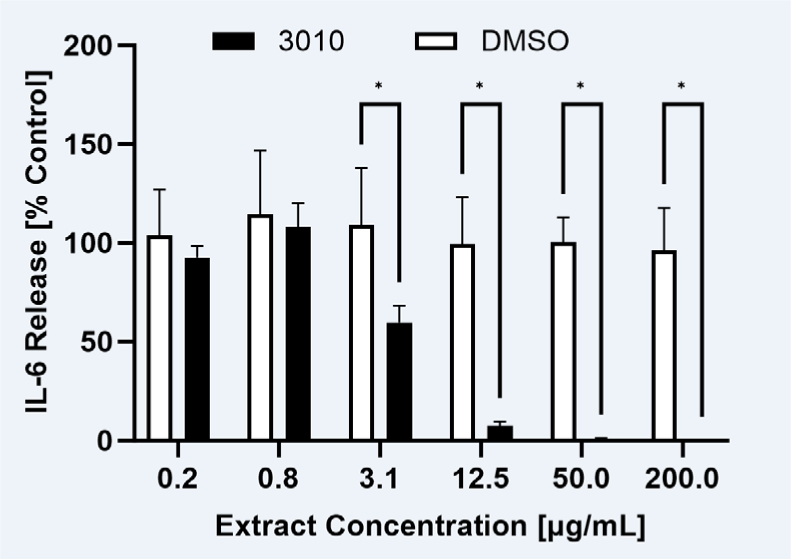
IL-6 release in LPS stimulated PBMCs incubated with *Vincetoxicum hirundinaria* root (sample 3010). The sample was tested at the concentrations 200, 50, 12.5, 3.1, 0.8, 0.2 µg/mL and a IC_50_ value of 3.62 ± 0.54 µg/mL was calculated. The dose-dependent activity was compared to the respective DMSO solvent control (two-way ANOVA with multiple comparisons, corrected for using Bonferroni’s method), revealing statistically significant activity at a dose of 3.1 µg/mL. Pairs with p < 0.05 were considered statistically significant and are marked with an asterisk (*).

## 4 Discussion

### 4.1 Assessment of the plants mentioned in RBH

In the first place, the investigation of the dermatological recipes in RBH shows the complexity behind the objective of analysing the plants and their uses in historical texts. In this study, we relied on previously suggested candidate plants to establish the botanical identities for the plant names mentioned in the historical text.

On the one hand, our method allowed us to pin down the botanical identity to one defined candidate species with a high chance of being the correct attribution (score “1”) in one third of the RBH plant names. Assuming historical consistency in the relevant plant names, the validity of many of these conclusions is supported by the fact that the respective candidate species is still referred to by the same plant name today. Often these are well-known herbs or medicinal plants, as for example “edle salbinen” for *Salvia officinalis*, “hannff” for *Cannabis sativa*, “holder” for *Sambucus nigra*, “lorberr” for *Laurus nobilis*, or “melissen” for *Melissa officinalis*.

However, in the remaining majority of the RBH plant names for which botanical identities could be established, the situation is complex. We exclude here the discussion of the well-established concepts ethnotaxa or plant complexes (see Leonti et al., 2015 and Linares and Bye, 1987), represented in RBH for example by the case of “wollen krut” which included various *Verbascum* species. What we observed is, that depending on the type of references used (glossaries, edited herbals, encyclopaedias) and the geographical and temporal context of the information, the different references often come to different conclusions or offer several different candidates for a particular plant name. In plant names with more than one possible candidate plants, usually one or few of them appear to have a higher chance of being the correct attribution (score “1”), while several additional attributions are less likely (scores “2” or “3”) or principally doubtful (“?”) (**Table 2**).

In the case of “heidnisch wundkrutt”, one of the plants contained in the Hallwyl wound potion, we found *Anthyllis vulneraria* and *Senecio ovatus* to be the candidates with the highest chance (score “1”). Support for this conclusion comes from Dobat and Dressendörfer (2016) as well as Hoppe (1969) who identified the plants in the 16th century herbals of Hieronymus Bock and Leonhard Fuchs, which are culturally and historically related to RBH. Further support with corresponding records from places in Switzerland is provided by Schweizerisches Idiotikon (2023) or Marzell’s lexicon of German plant names (1943-1979) and both species are also widespread in the region (Lauber et al., 2018). All the remaining candidates (*Actaea spicata, Hieracium murorum, Lactuca muralis, Saxifraga rotundifolia, Scrophularia nodosa, Senecio hercynicus, Solidago virgaurea*) are doubtful (“?”), because they are only mentioned by Marzell (1943-1979) and either “heidnisch wundkrutt” does not seem to be the name by which these species are addressed in the first place, or the corresponding records are not from Switzerland but from other German-speaking areas. As shown by Marzell (1943-1979), even in one and the same language region, the plant name in question is often only associated with a particular species in certain places, or the name refers to different species depending on the location. The case of “heidnisch wundkrutt” is further complicated by the fact that in some recipes a certain type of the plant, namely one with “cut” leaves (“mitt den zer schnitnen bletterenn”) was to be used (**Table 2**). Possible candidates for this type from the pool of candidate plants for “heidnisch wundkrutt” could only be determined by using morphological criteria such as the shape of the leaves. Apart from the inherent uncertainties in adopting existing identifications, this case also highlights the limitations of a method that relies on suggested candidate plants.

In response to this, recent historical studies on herbal *materia medica* in texts from the Mediterranean region have explored methods based on plant morphology to establish botanical identities or re- assess previous identifications. Using images and text from the historical source, Evergetis and Haroutounian (2015) constructed plant descriptions and compared these with information available in modern floristic works. In an interdisciplinary approach, Lardos et al. (2024) developed a multi-stage methodology that relies on comparative analyses including statistical evaluation of botanical and medicinal information drawn from both historical and modern sources.

### 4.2 Interpretation of the dermatological uses in RBH

In the majority of the cases, the vocabulary used in RBH to describe the skin problems to be treated by the recipes allowed us to draw possible conclusions about the underlying medical complaint. In this respect, our findings are in line with observations from other historical recipe books (see e.g., Zipser et al., 2023): The writers of these manuscripts often provide detailed descriptions of the skin problem in question or use specific disease terms on which an interpretation can be based.

For the interpretation of the medicinal uses, we relied on relevant literature on historical medical terms in German or the Swiss German idiom (Höfler, 1970; Riecke, 2004; Schweizerisches Idiotikon, 2023). The information contained in the references was critically assessed and in order to allow for all possible interpretations, care was taken to apply as broad a scope of interpretation as possible. Therefore, in most cases we offer more than one possible conclusion and at the same time assess these according to their presumed plausibility on the basis of the information available (**Table 1**). The interpretations developed also reflect some of the prevailing health problems and the epidemiological situation in Switzerland during the 16th century. Various uses of the recipes seem to indicate the presence of leprosy, smallpox and syphilis. That these serious infectious diseases were actually present in the region at the time is supported by corresponding historical studies (Furrer, 2022).

Although, we do not claim to have the final word on the interpretations of these historical medical uses, we consider the accumulated dataset to be a potentially helpful starting point with regard to the direction of any subsequent pharmacological studies of the plants in question. As pointed out in section 3.2.1, most of the uses are related to different kinds of wounds, inflammations of the skin, infectious diseases or neoplastic skin chances. In order to have any effect on these complaints, the plants used in the related recipes would have to have corresponding pharmacological activities. These include, for example, anti-inflammatory, immunomodulatory, antiproliferative, antineoplastic as well as antibiotic (including antibacterial, antifungal, antiparasitic) or antiviral effects.

The Hallwyl wound potion is an orally administered preparation. If this preparation were to have any beneficial effect on the mentioned dermatological conditions, the effect in question would have to be achieved by internal application. The pharmacological intervention in severe traumatic injuries with open or infected wounds, the main indications of the wound potion, includes not only a topical but often also an internal application of medication with antibiotic, anti-inflammatory, analgesic or haemostatic properties (Kang et al., 2019). It remains an open question, whether the wound potion could actually have developed its effect via these presumed pharmacological properties, or whether other factors based on traditional concepts for the perception or interpretation of an effect are decisive here.

### 4.3 Preliminary *in vitro* screening

In our preliminary screening endeavour with plants from RBH, we have investigated 17 crude extracts; 13 samples were prepared with one single plant and 4 samples were herbal mixtures of different reconstructions of the Hallwyl wound potion. Prior to screening the extract samples regarding their immunomodulatory potential, they were tested for possible cytotoxic effects in the MTT assay. None of the extracts showed a notable effect on cell viability except the extract of *Sanicula europaea*. Using the same assay, the same plant too was the source of the sample exerting the strongest cytotoxic effect in a screening of 23 plants of the Scandinavian folk medicine conducted by Ulriksen et al. (2018).

With the aim to use a cell-based *in vitro* assay for screening the crude extracts regarding their potential to modulate the release of LPS-induced IL-6 and TNF, we have investigated the suitability of a whole blood assay. In contrast to widely used PBMC based assays to explore immune modulation, the whole blood assay has the advantage of more closely mimicking *in vivo* conditions (Coch et al., 2013). Moreover, in whole blood, cells and soluble proteins influencing pharmacokinetics and the inflammatory response are present in unchanged form (Ushiyama et al., 2008). To avoid screening artifacts due to the use of unphysiologically high concentrations in *in vitro* assays as pointed out by Gertsch (2009), extract samples were tested at a concentration of 50 μg/mL and pure compounds at 5 μM. We observed mixed results ranging from inactivity to low or moderate inhibition (e.g. *Vincetoxicum hirundinaria* root) or to strong increase of cytokine release (e.g. *Solanum nigrum* roots and leaves, *Solanum dulcamara* aerial parts). In none of the extracts, except for the sample of *Sanicula europaea*, the concentration at which a cytotoxic effect could be detected, if at all, was in the range of the extract concentration of 50 µg/mL used in the whole blood assay (**Table 4**). In view of the results on the modulation of cytokine release, it should also be noted that various factors could have an influence on the readouts of the whole blood assay, such as oestrogen or cortisol levels in the donor blood (Schmid et al., 2009; Ritchie et al., 2011), or the high affinity of several plant compounds to bind to serum proteins such as albumin (Boulton et al., 1998; López-Yerena et al., 2020). It was not possible to pursue these issues in the scope the present study.

However, the tendencies observed with our extract samples in the whole blood assay are in line with Elisia et al. (2018), who also found either inhibitory or stimulatory effects on the release of IL-6 for the plant extracts they tested.

A stimulation of pro-inflammatory cytokines might in fact be indicated for the use described in one of the relevant RBH recipes (121b,116r.02): The juice of the roots of “nachtschatten”, most likely *Solanum nigrum*, is applied as a poultice in cases of “figwertzenn” which is primarily interpreted as genital or syphilitic warts (**Supplementary Material, Table S1**). Immunostimulatory compounds such as imidazoquinolines inducing secretion of TNF and other cytokines, have been used clinically for the treatment of external genital warts and topically manifested viral infections (Hengge and Cusini, 2003). From this perspective, the distinct stimulation of cytokine release observed in whole blood for the extract of *Solanum nigrum* roots at 50 µg/mL (IL-6, 154%; TNF, 240%) would support the validity of the historical use.

In any case, plants with either immunosuppressant or immunostimulant activity could be of interest in dermatology. Especially in the context of wound healing, the possibility to modulate the immune response is becoming increasingly important, both to slow down excessive immune reactions in chronic inflammatory skin disorders or to enhance immune reactions in order to stimulate granulation tissue as for example in leg ulcers (ulcus cruris) (Ramm-Fischer, 2021).

The data of the six extract samples investigated in the PBMC assay, partly resonate but also diverge from the ones of the whole blood assay. Elisia et al. (2018) even found a general lack of correlation between the readouts of the whole blood assay and the PBMC assay regarding the modulation of cytokine release. In our results on the PBMC assay the distinct and significant inhibitory effects on IL-6 release of the two extract samples of *Vincetoxicum hirundinaria* (aerial parts and root) stand out. The root of the plant (sample 3010) revealed an IC_50_ of 3.6 µg/mL (**Table 4**) and was able to almost block the release of the cytokine with statistical significance at a concentration of 3.1 µg/mL (**Figure 1**).

The medicinal properties of the species are often associated with the phenanthroindolizidine alkaloids (PIA) found in plants of the genus *Vincetoxicum* as well as in other genera. *In vitro* studies found potent anti-inflammatory activities in RAW264.7 mouse macrophage cell line for different PIAs (Yang et al., 2006; Chou et al., 2017; Reimche et al., 2022). However, PIAs are also known to be strongly cytotoxic and have therefore also been explored in terms of their anticancer properties. PIAs isolated from *Vincetoxicum hirundinaria* Medik. (*Cynanchum vincetoxicum* (L.) Pers.) and *Vincetoxicum mukdenense* Kitag. (*Cynanchum paniculatum* (Bunge) Kitag.) showed pronounced inhibitory activities in numerous cancer cell lines (Wiart, 2013). Although we were not able to find studies conducted with extracts of *Vincetoxicum hirundinaria* in this context, the activities reported for the PIAs seem to correlate with the use of the relevant RBH recipe, such as different inflammatory conditions or perhaps also tumours (**Supplementary Material, Table S6**).

In our attempt to approximate the historical preparation of the Hallwyl wound potion, we were not always able to follow the details described in the recipe, such as type of solvent or reaction vessels, or important information such as was missing (e.g. correct dilutions) or left a wide space of interpretation (extraction time, quantities). For example, the recipe of the wound potion calls for the use of “new glazed pottery” as a reaction vessel, which raises the question whether the lead silicate glaze used at the time (Bouquillon et al., 2017) had any influence on the decoction prepared in it, especially when using an acidic solvent like white wine as prescribed in the recipe (**Supplementary material, Table S5**).

Regarding the results of the whole blood assay for the different samples of the Hallwyl wound potion, only the version approximating the historical preparation with *Senecio ovatus* as the alternative candidate for the RBH plant “heidnisch wundkrutt” exhibited a noteworthy effect in form of a weak stimulation of the release of IL-6 and TNF (132.4%, p = 0.062; 116.9% %, p = 0.028). Apart from *Alchemilla xanthochlora* as to the release of TNF, none of the other plant ingredients of the wound potion (*Artemisia vulgaris, Beta vulgaris, Pyrola rotundifolia, Sanicula europaea, Senecio ovatus*), when tested on their own, exhibited a comparable effect. This could indicate the presence of additive or perhaps even synergistic effects in the herbal mixture (**Table 4**). Other combinations of alternative candidates for the relevant herbal ingredients used to prepare the mixture (see **Table 2**) might lead to immunomodulatory effects differing from the ones observed in this study. All this offers various possibilities for further research into the Hallwyl wound potion.

## Conclusions

Using the example of RBH, this study illustrates a possible ethnopharmacological path from unlocking the historical text and its subsequent analysis, through the selection and collection of plant candidates to their investigation in a preliminary *in vitro* screening.

The pharmaco-botanical focus in the text analysis points up the challenges associated with the assessment of historical plant names. Our case study shows that often two or more plant species are available as candidates for one historical plant name. This fact should be considered for subsequent investigations. At the same time, the dermatological plant uses mentioned in RBH illustrate that in many cases the information provided in the text about the skin problem in question enables conclusions to be drawn based on which the presumed underlying medical condition and its pharmacological basis can be narrowed down. This consequently facilitates decision-making when selecting and designing suitable biological test systems. Fully documenting our approach to the analysis of historical texts, we hope to contribute to the discussion on solutions for the digital indexing of premodern information on the use of plants or other natural products.

For the preliminary *in vitro* screening, we selected 11 candidate species, four of which were used in RBH in herbal simple recipes and seven in a herbal compound formulation. None of the crude extracts prepared from these plants, except for the sample of *Sanicula europaea*, showed a noteworthy effect on cell viability and critical concentrations, if detectable at all, were not in the range of the extract concentration (50 µg/mL) used in the whole blood assay. In the PBMC assay, the roots of *Vincetoxicum hirundinaria* revealed a distinct inhibitory effect on IL-6 release (IC_50_ of 3.6 µg/mL). Although the plant or its compounds would likely be sorted out during dereplication, this finding together with the results on cell viability, point to the potential value of RBH as a resource for further pharmacological investigations. From a methodological perspective, the whole blood assay, being fast and comparatively cheap, could be an interesting alternative for testing herbal substances. A wider application in this context would be needed to clarify its potential. Finally, the observations made during this preliminary study indicate the need for defined protocols for the *in vitro* screening of plants in ethnopharmacology.

## CRediT author statement

JS: Methodology, Formal analysis, Investigation, Resources, Data curation, Writing – Original Draft, Writing – Review & Editing

IA: Methodology, Investigation, Writing – Review & Editing

TF: Conceptualisation, Resources, Writing – Review & Editing, Funding acquisition

BFH: Conceptualisation, Methodology, Resources, Writing – Review & Editing, Supervision, Project administration, Funding acquisition

AL: Conceptualisation, Methodology, Formal analysis, Investigation, Resources, Data curation, Writing – Original Draft, Writing – Review & Editing, Visualization, Supervision, Project administration, Funding acquisition

## Supporting information

Supplemental Figues S1 to S3

## Acknowledgements

We thank Evelyn Wolfram, Nina Vahekeni and Samuel Peter from the ZHAW (Zurich University of Applied Sciences) for advice in the planning and realisation of the laboratory-specific work.

## Appendix A. Supplementary Material

See separate data files

## Glossary

|  |  |
| --- | --- |
| DMEM-HG | Dulbecco's modified Eagle's medium – High Glucose |
| DMSO | Dimethoxysulfoxid |
| EtOH | Ethanol |
| ELISA | Enzyme-linked Immunosorbent Assay |
| FBS | Fetal Bovine Serum |
| IL-6 | Interleukin 6 |
| LPS | Lipopolysaccharide |
| MTT | Tetrazolium Dye |
| OD | Optical Density |
| PBMC | Peripheral Blood Mononuclear Cells |
| PBS | Phosphate-buffered Saline |
| TNF | Tumor Necrosis Factor |

