## Supplemental Figues S1 to S3 for "Dermatological plants and their uses in the *Receptarium* of Burkhard III from Hallwyl from 16^th^ century Switzerland – Data mining a historical text and preliminary *in vitro* screening"

**SM - Figure S1**. Illustration of the relationships between the 20 tables of the MS Access® database of the Receptarium of Burkhard from Hallwyl (RBH).

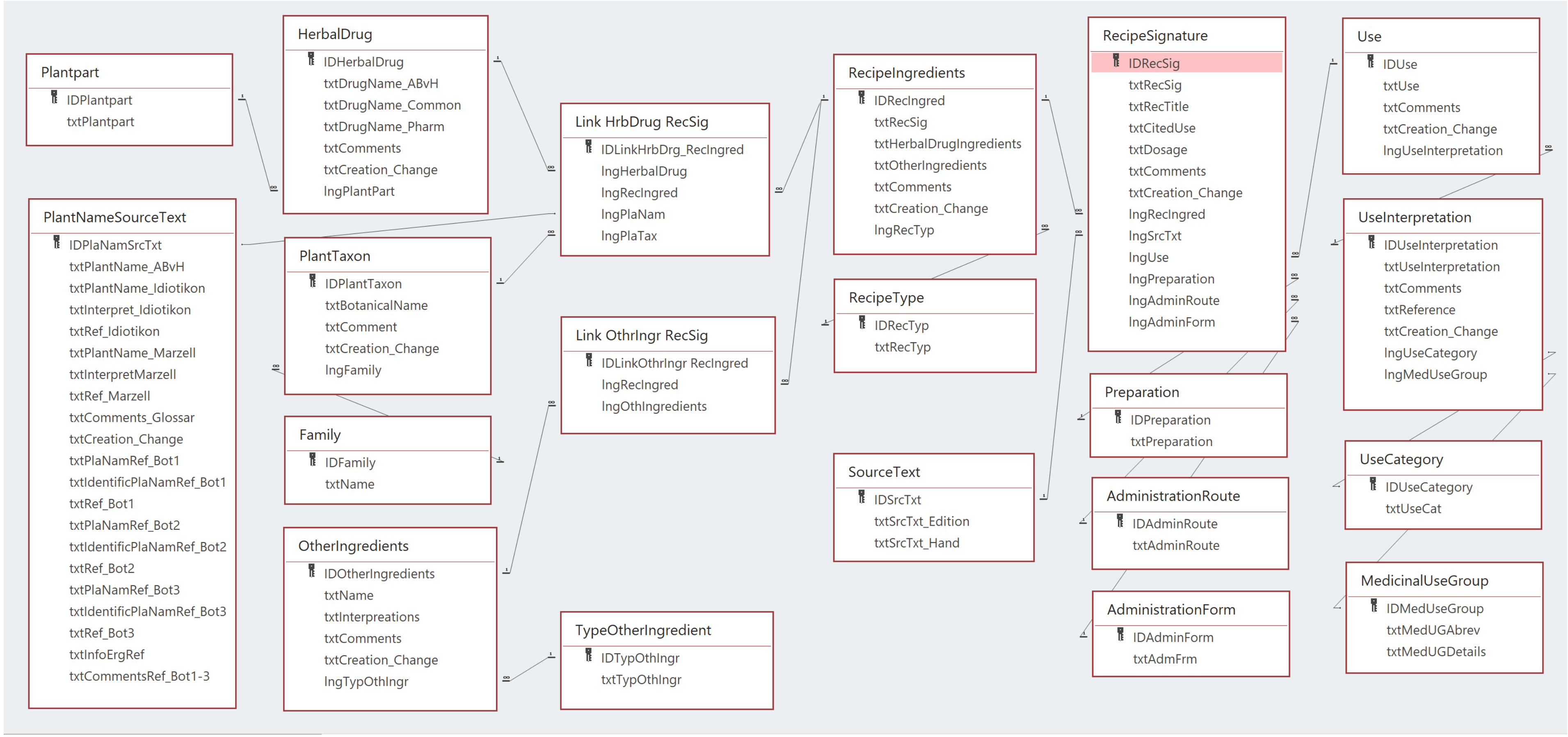


**Dermatological plants in the *Receptarium* of Burkhard III from Hallwyl from 16th century Switzerland – Data mining a historical text and preliminary *in vitro* screening**

February 2024


*Jonas Stehlin, Ina Albert, Thomas Frei, Barbara Frei Haller, Andreas Lardos*

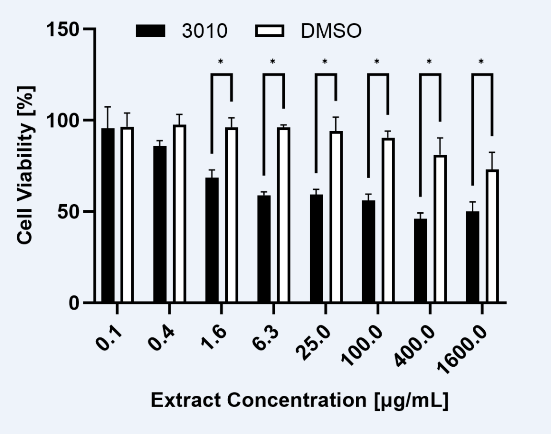


**SM – Figure S2.** Results of the extended MTT experiment with eight dose points with the ethanolic macerate of *Vincetoxicum hirundinaria* roots (sample 3010) in NIH-3T3 murine fibroblast cell line. The cell viability at individual dose points was compared to the relevant solvent control using two-way ANOVA with multiple comparisons corrected for using Bonferroni's method. Pairs with $p < 0.05$ were considered statistically significant and are marked with an asterisk (*).

**Dermatological plants in the *Receptarium* of Burkhard III from Hallwyl from 16th century Switzerland – Data mining a historical text and preliminary *in vitro* screening**

February 2024

*Jonas Stehlin, Ina Albert, Thomas Frei, Barbara Frei Haller, Andreas Lardos*

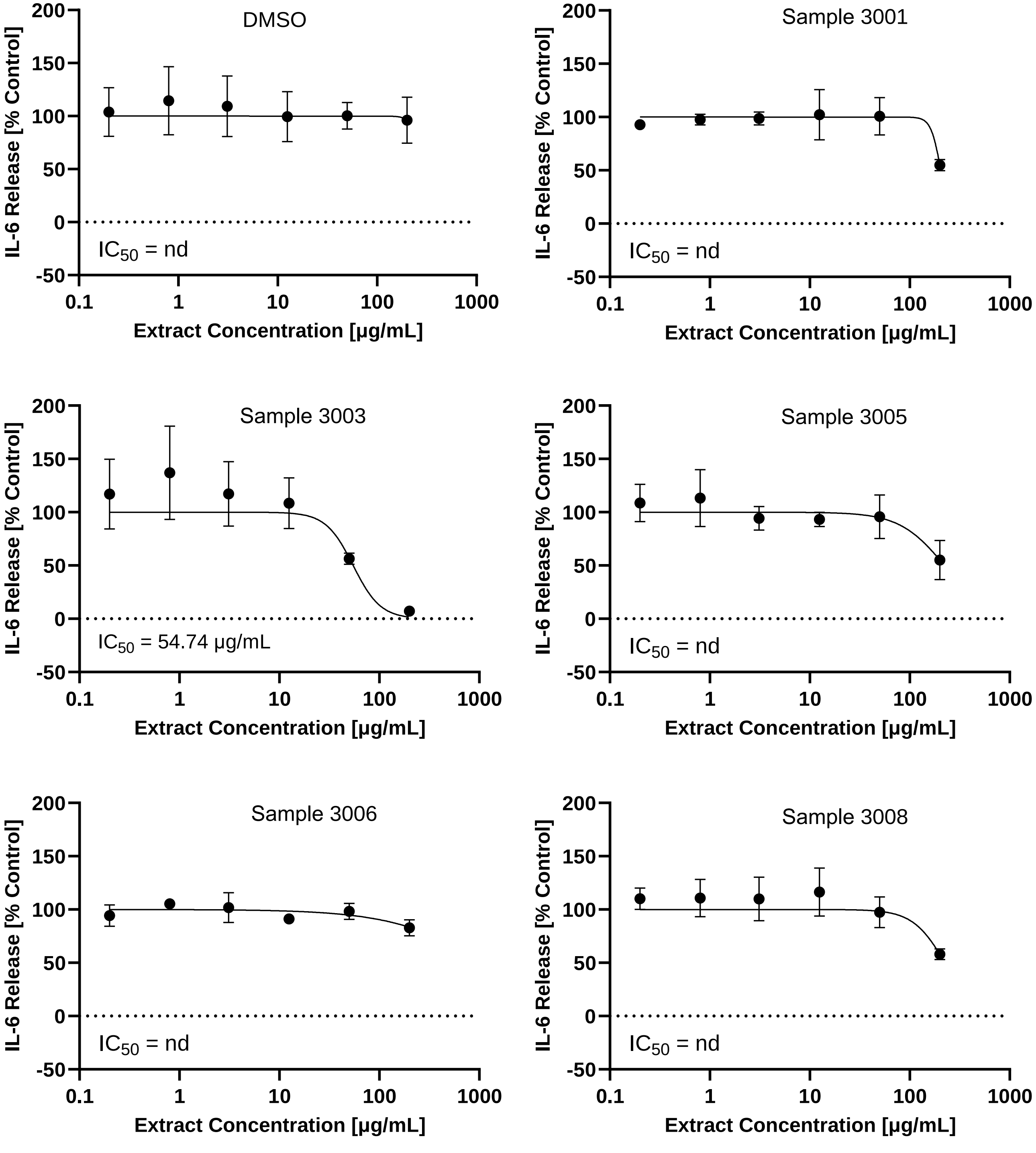


**Supplementary Material - Figure S3**. Dose response curves of five of the six samples of crude extracts (3001, 3003, 3005, 3006, 3008) investigated in the PBMC assay (Details about the samples, see Table 4; Data of sample 3010, see Figure 1). All samples including the vehicle control DMSO were tested at the concentrations 200, 50, 12.5, 3.1, 0.8 and 0.2 µg/mL. Except for sample 3003 (and 3010 - see Figure 1), in all samples IC50 values are above 200 µg/mL or cannot be calculated due to out of range values (nd = not determined).

**Dermatological plants and their uses in the *Receptarium* of Burkhard III from Hallwyl (RBH) from 16th century Switzerland – Data mining a historical text and preliminary in vitro screening –**

*Jonas Stehlin, Ina Albert, Thomas Frei, Barbara Frei Haller, Andreas Lardos* - February 2024
