## Supplemental Tables S1 to S6 for "Dermatological plants and their uses in the *Receptarium* of Burkhard III from Hallwyl from 16^th^ century Switzerland – Data mining a historical text and preliminary *in vitro* screening"

**SM – Table S1 – Recipes** (Extract from the RBH database on dermatological recipes)

| txtRecSig | txtRecTyp | txtCitedUse | txtCombinedUse | txtAdminRoute |
|---|---|---|---|---|
| 051b,45r.01 | Simple herbal | geschwulsten vnnd wie die zU vertribenn, Erstlich ann eim schenckel | geschwulst | Topical (external) |
| 051b,45r.02 | Composed herbal | Für die geschwulst | geschwulst | Topical (external) |
| 051b,45r.02.z01 | Composed mixed | Für die geschwulst | geschwulst | Oral (ingested) |
| 051b,45r.03.v01 | Composed herbal | Für alle geschwulst | geschwulst | Oral (ingested) |
| 051b,45r.03.v02 | Composed herbal | Für alle geschwulst | geschwulst | Oral (ingested) |
| 051b,45r.04 | Composed mixed | Für nüw erhabne geschwulst | geschwulst | Oral (ingested) |
| 051b,45r.05 | Composed mixed | Für nüw erhabne geschwulst, Einn annders | geschwulst | Topical (external) |
| 051b,45r.06 | Composed herbal | Für nüw erhabne geschwulst | geschwulst | Topical (external) |
| 052a,45v.01 | Simple herbal | zertheillt vnnd nider geleidt werden, vnnd zur heillunng komen alle geschwulst vnnd knollen | geschwulst | Not mentioned |
| 052a,45v.02 | Composed mixed | Für gschwulst so vonn überiger hitz kombt es sig vonn wundern. törnn. stëchen schlahen oder gfallen | geschwulst vonn überiger hitz kombt, es sig vonn wundern, törnn, stëchen, schlahen oder gfallen | Topical (external) |
| 052a,45v.03 | Simple herbal | für alle gschwulst gUtt | geschwulst | Not mentioned |
| 052a,45v.04.v01 | Composed mixed | bösse hitzige gesch wulsten schennckel | geschwulst | Topical (external) |
| 052a,45v.04.v02 | Composed mixed | schonn röttj | schonn röttj | Topical (external) |
| 052a,45v.05.v01 | Simple herbal | für die gschwulst, der beinenn vnnd schëncklen | geschwulst | Topical (external) |
| 052a,45v.05.v02 | Simple herbal | für die gschwulst, der beinenn vnnd schëncklen | geschwulst | Topical (external) |
| 052b,46r.02 | Simple herbal | die bosen schenckell, vertribt die gschwulst, so vonn keltte vnnd müde kompt | geschwulst | Topical (external) |
| 052b,46r.03 | Composed herbal | gschwulst | geschwulst | Topical (external) |
| 052b,46r.05 | Simple herbal | Für alle geschwulst | geschwulst | Topical (external) |
| 078b,72r.01 | Simple herbal | blUtt stellunng | blUtt stellunng | Topical (external) |
| 078b,72r.06 | Composed mixed | Gestockt blUtt vs trÿben | Gestockt blUtt vs trÿben | Oral (ingested) |
| 079a,72v.01 | Simple herbal | blUtt verstellen | blUtt stellunng | Topical (external) |
| 079a,72v.03 | Simple herbal | so aber das blUtt selber kompt von der nasen | Nasën blUtten | Topical (external) |
| 079a,72v.03.z01 | Simple herbal | so aber das blUtt selber kompt von der nasen | Nasën blUtten | Oral (ingested) |
| 079a,72v.04 | Simple herbal | legs vff das ver wundt orth, so hörtt es blUtten | blUtt stellunng | Topical (external) |

**SM – Table S1 – Recipes** (Extract from the RBH database on dermatological recipes)

| txtRecSig | txtRecTyp | txtCitedUse | txtCombinedUse | txtAdminRoute |
|---|---|---|---|---|
| 079a,72v.05 | Composed herbal | thUn das vnder ein anderen thU das buluer in die wunden es stadt für warr [blUtt ze stellen] | blUtt stellunng | Topical (external) |
| 079a,72v.06.v01 | Composed herbal | so aber das blUtt selber kompt von der nasen | Nasën blUtten | Topical (external) |
| 079a,72v.07 | Simple herbal | nasën blUtten | Nasën blUtten | Topical (external) |
| 079a,72v.08 | Composed herbal | blUtt ze stellen | blUtt stellunng | Topical (external) |
| 079a,72v.09 | Simple herbal | gestellt das blUtt gar wunder barlichen | blUtt stellunng | Not mentioned |
| 079a,72v.10 | Simple herbal | Einn anders gwüs blUtt zestellen | blUtt stellunng | Not mentioned |
| 079a,72v.11 | Simple herbal | gwüs blUtt zestellen | blUtt stellunng | Topical (external) |
| 079b,73r.01 | Composed herbal | edle blUtt stellung | blUtt stellunng | Not mentioned |
| 079b,73r.04 | Composed herbal | Wem blUtt zwüschen hutt vnnd fleisch | blUtt zwüschen hutt vnnd fleisch | Topical (external) |
| 079b,73r.05 | Simple herbal | blUtt stellenn | blUtt stellunng | Topical (external) |
| 094b,89r.05 | Composed herbal | Für denn Stich | stich | Topical (external) |
| 103b,98r.04 | Composed mixed | zU villen presten | presten, zu villen bresten [äusserlich behandelt] | Topical (external) |
| 103b,98r.04.z01 | Simple herbal | zU villen presten | presten, zu villen bresten [äusserlich behandelt] | Topical (external) |
| 104a,98v.02 | Composed mixed | heilett brannd vnnd erfrores | Branndt | Topical (external) |
| 104a,98v.03 | Composed mixed | bruch die zum schaden, fürr den Branndt | Branndt | Topical (external) |
| 104b,99r.01 | Composed mixed | wund salben | wunden | Topical (external) |
| 105a,99v.02 | Composed mixed | Blattern | Blattern | Topical (external) |
| 105a,99v.03 | Composed mixed | Wunnd salb | wunden | Topical (external) |
| 105a,99v.05 | Composed mixed | zU alltenn schadenn | zU alltenn schadenn | Topical (external) |
| 105b,100r.01 | Composed mixed | für alerleÿ flüs im anngesicht | flüs jm angsicht | Topical (external) |
| 105b,100r.02 | Composed mixed | so hast ein gUtte brannd salben | Branndt | Topical (external) |
| 105b,100r.03 | Composed mixed | zU offnenn schaden Vnnd flüssenn | zU offnenn schaden Vnnd flüssenn | Topical (external) |
| 106a,100v.01 | Composed mixed | zU frischen wunndenn | wunden | Topical (external) |
| 106a,100v.02 | Composed mixed | Uber alle geschwärr | geschwërr, geschwärr, geschwerr | Topical (external) |
| 106a,100v.03 | Simple herbal | Wunnden zU samen ziechen | wunden | Topical (external) |

**SM – Table S1 – Recipes** (Extract from the RBH database on dermatological recipes)

| txtRecSig | txtRecTyp | txtCitedUse | txtCombinedUse | txtAdminRoute |
|---|---|---|---|---|
| 106a,100v.05 | Composed mixed | wunnd salben | wunden | Topical (external) |
| 106b,101r.01 | Composed mixed | Houwt, oder sticht oder brent oder böss eissenn am Lÿb Hatt | Houwt, oder sticht oder brent oder böss eissenn am Lÿb Hatt | Topical (external) |
| 106b,101r.02 | Composed mixed | zU allerlei Wunnden vnnd annderem | wunden | Topical (external) |
| 106b,101r.03 | Simple herbal | haupt wunden, ist nitt also heillsam. sonnder hefftett die selben zU samen jnn kurtzerr zitt, vnnd zücht auch vs die sprisenn der hirnn schallen | haupt wunden, Houbt wunnden | Topical (external) |
| 106b,101r.04.v02 | Simple herbal | sÿ süberett auch wunden | wunden | Topical (external) |
| 106b,101r.04.v03 | Simple herbal | verthribt auch fleckenn | fleckenn | Topical (external) |
| 106b,101r.05.v01 | Composed mixed | wunden | wunden | Topical (external) |
| 106b,101r.05.v02 | Composed mixed | allerleÿ geschwulst | geschwulst | Topical (external) |
| 106b,101r.06.v01 | Composed herbal | Altt schadenn, heillett wunderlichen | zU alltenn schadenn | Topical (external) |
| 106b,101r.06.v02 | Composed herbal | heillett wunder-lichen, auch fistalam | fistel, fistalam | Topical (external) |
| 107a,101v.01 | Composed mixed | Schwartz blatteren | Schwartz blatteren | Topical (external) |
| 107a,101v.02 | Composed mixed | für die schwartzen Blatterenn | Schwartz blatteren | Topical (external) |
| 107a,101v.03 | Composed mixed | einn ietliche blatterr | Blattern | Topical (external) |
| 107a,101v.05 | Simple herbal | gschwärr | geschwërr, geschwärr, geschwerr | Topical (external) |
| 107a,101v.06.v01 | Composed herbal | geschwerr | geschwërr, geschwärr, geschwerr | Topical (external) |
| 107a,101v.06.v02 | Composed herbal | geschwërr | geschwërr, geschwärr, geschwerr | Topical (external) |
| 107a,101v.07 | Composed mixed | geschwërr | geschwërr, geschwärr, geschwerr | Topical (external) |
| 107a,101v.08.v01 | Composed mixed | eÿssenn | eÿssenn | Topical (external) |
| 107a,101v.08.v02 | Composed mixed | gUtt blatterenn | Blattern | Topical (external) |
| 107a,101v.09 | Composed mixed | Schrunnden an hennden | Schrunnden an hennden | Topical (external) |
| 107b,102r.01 | Composed herbal | wunnden | wunden | Topical (external) |
| 107b,102r.02.v01 | Composed herbal | wunden welche full fleisch erzügt, vertribt auch die hitz | zU fullen wunden, bewertt vor fullem fleisch | Topical (external) |
| 107b,102r.03 | Composed mixed | geschwerren | geschwërr, geschwärr, geschwerr | Topical (external) |
| 107b,102r.04 | Composed mixed | wunden oder ein eissen | wunden | Topical (external) |
| 108a,102v.01.v01 | Composed mixed | wunden | wunden | Topical (external) |

**SM – Table S1 – Recipes** (Extract from the RBH database on dermatological recipes)

| txtRecSig | txtRecTyp | txtCitedUse | txtCombinedUse | txtAdminRoute |
|---|---|---|---|---|
| 114b,109r.01 | Composed mixed | zU alttenn schaden, es reiniget zücht fleisch trücknett vnnd rimpfft zU samen | zU alltenn schadenn | Topical (external) |
| 114b,109r.02.v01 | Composed mixed | zU altenn scheden welchs etzt vnnd züch zU eitter reinigt | zU alltenn schadenn | Topical (external) |
| 114b,109r.04 | Composed mixed | ZU allten schädenn | zU alltenn schadenn | Topical (external) |
| 114b,109r.05.v01 | Composed mixed | allt schaden, allte ruden | zU alltenn schadenn | Topical (external) |
| 114b,109r.05.v02 | Composed mixed | als kräps, wolff, nolime tangere vnnd des glich | kräps, wolff, noli me tangere, vnnd des glich | Topical (external) |
| 114b,109r.05.v03 | Composed mixed | rünende bein | rünende bein | Topical (external) |
| 116b,111r.01.v01 | Simple herbal | gUtt den menschen, so sich förchten vor der vss setzig keitt | vssetzigkeitt, wer lanng sich be sorgett vor ... | Oral (ingested) |
| 116b,111r.01.v02 | Simple herbal | für alle kretz vnd rüdigkeitt | kretz vnd rüdigkeitt | Oral (ingested) |
| 116b,111r.01.v05 | Simple herbal | flüs jm angsicht | flüs jm angsicht | Topical (external) |
| 116b,111r.01.v06 | Simple herbal | so einner grunenn blUtt bÿ jm hatt | blUtt grunnen | Not mentioned |
| 117a,111v.01.v01 | Simple herbal | alle wunden, vnnd schaden, mitt dissem wasser gewäschen reinigett vnnd bewertt das vor fullem fleisch, vnnd ist gUtt für rünende wunden | zU fullen wunden, bewertt vor fullem fleisch | Topical (external) |
| 117a,111v.01.v02 | Simple herbal | fistell | fistel, fistalam | Topical (external) |
| 117a,111v.01.v03 | Simple herbal | wer lanng sich be sorgett vor vssetzigkeitt der bruch des wasserrs | vssetzigkeitt, wer lanng sich be sorgett vor ... | Oral (ingested) |
| 117a,111v.01.v04 | Simple herbal | werr den vs satz mitt dissem wasserr wescht offt vnnd dick es hinndertt vnnd ver-hütett den lanng | vs satz, vssetzigkeitt | Topical (external) |
| 117a,111v.02.v01 | Simple herbal | alle wunden, vnnd schaden, mitt dissem wasser gewäschen reinigett vnnd bewertt das vor fullem fleisch, vnnd ist gUtt für rünende wunden | zU fullen wunden, bewertt vor fullem fleisch | Topical (external) |
| 117a,111v.02.v02 | Simple herbal | fistell | fistel, fistalam | Topical (external) |
| 117a,111v.02.v03 | Simple herbal | wer lanng sich be sorgett vor vssetzigkeitt | vssetzigkeitt, wer lanng sich be sorgett vor ... | Oral (ingested) |
| 117a,111v.02.v04 | Simple herbal | werr den vs satz mitt dissem wasserr wescht offt vnnd dick es hinndertt vnnd verhütett den lanng | vs satz, vssetzigkeitt | Topical (external) |
| 117b,112r.01.v02 | Simple herbal | reinigett alle bösse füchtigkeitt | füchtigkeitt - reinigett füchtigkeitt | Topical (external) |
| 117b,112r.01.v03 | Simple herbal | vor alle apostema vnnd geschwer | apostema vnnd geschwer | Oral (ingested) |
| 117b,112r.01.v05 | Simple herbal | gschwulst alls büllen | geschwulst | Oral (ingested) |
| 117b,112r.02.v02 | Composed mixed | Uber allen prestenn gelegt | presten, zu villen bresten [äusserlich behandelt] | Topical (external) |
| 117b,112r.02.v03 | Composed mixed | zU fullen wunden | zU fullen wunden, bewertt vor fullem fleisch | Topical (external) |

**SM – Table S1 – Recipes** (Extract from the RBH database on dermatological recipes)

| txtRecSig | txtRecTyp | txtCitedUse | txtCombinedUse | txtAdminRoute |
|---|---|---|---|---|
| 117b,112r.02.v05 | Composed mixed | für fläcken jnn angesicht | fleckenn | Topical (external) |
| 117b,112r.02.v12 | Composed mixed | verthribt alles geschwerr | geschwërr, geschwärr, geschwerr | Not mentioned |
| 117b,112r.02.v15 | Composed mixed | büllen, vnnd geschwulst | geschwulst | Topical (external) |
| 117b,112r.02.v17 | Composed mixed | es thribt den vssatzt | vs satz, vssetzigkeitt | Topical (external) |
| 118a,112v.02.v07 | Simple herbal | zU allen wunden | zU fullen wunden, bewertt vor fullem fleisch | Topical (external) |
| 118a,112v.02.v08 | Simple herbal | bössenn schaden | bössenn schaden | Topical (external) |
| 118a,112v.02.v18 | Simple herbal | alle geschwärr heilett vnnd weicht es | geschwërr, geschwärr, geschwerr | Topical (external) |
| 120b,115r.01 | Composed herbal | figwertzenn | figwertzenn, fÿg wertzenn, fÿg Blatterenn | Topical (external) |
| 120b,115r.02 | Composed mixed | figwertzenn die zU vertribenn | figwertzenn, fÿg wertzenn, fÿg Blatterenn | Rectal |
| 120b,115r.03.v01 | Composed mixed | figwertzenn die zU vertribenn | figwertzenn, fÿg wertzenn, fÿg Blatterenn | Topical (external) |
| 120b,115r.03.v02 | Composed mixed | wen ein mensch das harnn hatt | wen ein mensch das harnn hatt | Topical (external) |
| 120b,115r.05.v03 | Composed mixed | fÿg wertzenn | figwertzenn, fÿg wertzenn, fÿg Blatterenn | Topical (external) |
| 121a,115v.01 | Composed herbal | Für fÿgwertzenn | figwertzenn, fÿg wertzenn, fÿg Blatterenn | Topical (external) |
| 121a,115v.02 | Simple herbal | Für fÿgwertzenn | figwertzenn, fÿg wertzenn, fÿg Blatterenn | Topical (external) |
| 121a,115v.03 | Composed mixed | Für fÿgwertzenn | figwertzenn, fÿg wertzenn, fÿg Blatterenn | Topical (external) |
| 121a,115v.04 | Simple herbal | heilett gruntlich die fÿgwërtzen | figwertzenn, fÿg wertzenn, fÿg Blatterenn | Oral (ingested) |
| 121a,115v.05 | Simple herbal | fÿg Blatterenn | figwertzenn, fÿg wertzenn, fÿg Blatterenn | Topical (external) |
| 121a,115v.06 | Simple herbal | fÿg Blatterenn | figwertzenn, fÿg wertzenn, fÿg Blatterenn | Topical (external) |
| 121b,116r.01 | Composed mixed | fÿg blatterenn | figwertzenn, fÿg wertzenn, fÿg Blatterenn | Topical (external) |
| 121b,116r.02 | Simple herbal | Man sol wüssenn, das drierlein wertzen sinnd, das manns nitt mitt einerlei vertribt, die ersten vertribt man mitt ... | figwertzenn, fÿg wertzenn, fÿg Blatterenn | Topical (external) |
| 121b,116r.03 | Simple herbal | Man sol wüssenn, das drierlein wertzen sinnd, das manns nitt mitt einerlei vertribt, ... die anderen vertribt man mitt ... | figwertzenn, fÿg wertzenn, fÿg Blatterenn | Topical (external) |
| 121b,116r.03.z01 | Composed mixed | Man sol wüssenn, das drierlein wertzen sinnd, das manns nitt mitt einerlei vertribt, ... die annderen vertribt man...: wen ein mensch da gschwollen das er nitt kan zU stUll gang | figwertzenn, fÿg wertzenn, fÿg Blatterenn | Topical (external) |
| 121b,116r.05 | Simple herbal | fig wertzen | figwertzenn, fÿg wertzenn, fÿg Blatterenn | Topical (external) |
| 121b,116r.06 | Simple herbal | fig wertzen | figwertzenn, fÿg wertzenn, fÿg Blatterenn | Topical (external) |

**SM – Table S1 – Recipes** (Extract from the RBH database on dermatological recipes)

| txtRecSig | txtRecTyp | txtCitedUse | txtCombinedUse | txtAdminRoute |
|---|---|---|---|---|
| 122a,116v.01 | Simple herbal | fig wertzenn | figwertzenn, fÿg wertzenn, fÿg Blatterenn | Topical (external) |
| 122a,116v.02 | Simple herbal | Für fig wertzenn | figwertzenn, fÿg wertzenn, fÿg Blatterenn | Topical (external) |
| 122a,116v.02.z01 | Composed mixed | Für fig wertzenn | figwertzenn, fÿg wertzenn, fÿg Blatterenn | Topical (external) |
| 122a,116v.04 | Simple herbal | Wertzen | Wertzenn, wartzen | Topical (external) |
| 122a,116v.05 | Simple herbal | Wertzen | Wertzenn, wartzen | Topical (external) |
| 122a,116v.06 | Composed herbal | Drüssenn vertribenn | Drüssenn | Topical (external) |
| 122a,116v.07 | Composed mixed | ist gUtt für fläcken | fleckenn | Topical (external) |
| 127a,121v.01 | Simple herbal | für die rud Vnnd denn magerr | rud vnnd magerr | Topical (external) |
| 127a,121v.02 | Composed mixed | Für denn magerr | rud vnnd magerr | Topical (external) |
| 127a,121v.03 | Composed mixed | Für denn bössenn magerr | rud vnnd magerr | Topical (external) |
| 127a,121v.04 | Simple herbal | Für das juckenn | juckenn | Topical (external) |
| 127a,121v.05 | Composed mixed | Für die rud | rud vnnd magerr | Topical (external) |
| 127a,121v.07 | Composed mixed | Für denn magerr, rud vnnd annderr boss schadenn | rud vnnd magerr | Topical (external) |
| 127b,122r.01 | Composed mixed | für denn magerr | mager, magerr | Topical (external) |
| 127b,122r.02 | Composed mixed | für den magerr | mager, magerr | Topical (external) |
| 127b,122r.03 | Simple herbal | Einn annders für denn magerr | mager, magerr | Topical (external) |
| 127b,122r.05 | Composed mixed | für die rud | rud | Topical (external) |
| 127b,122r.06 | Composed mixed | Für die rud | rud | Topical (external) |
| 127b,122r.07 | Composed mixed | so die kinndt bösse füslin hannd | so die kinndt bösse füslin hannd | Topical (external) |
| 128b,123r.01 | Composed mixed | einn angesicht als obs vss setzig sige | vs satz, vssetzigkeitt | Topical (external) |
| 128b,123r.02 | Simple herbal | Flüss jm anngesicht | flüs jm angsicht | Topical (external) |
| 131a,125v.01 | Composed herbal | wurm am finnger | wurm am finnger | Topical (external) |
| 131a,125v.02 | Composed mixed | Wurms annfanng; wurm am finnger | wurm am finnger | Topical (external) |
| 131a,125v.09 | Simple herbal | wurm am finngerr | wurm am finnger | Topical (external) |
| 131a,125v.10 | Composed herbal | wurm am finngerr | wurm am finnger | Topical (external) |
| 131b,126r.01 | Composed herbal | wurm am finngerr | wurm am finnger | Topical (external) |
| 131b,126r.02 | Simple herbal | wurm am finngerr | wurm am finnger | Topical (external) |

**SM – Table S1 – Recipes** (Extract from the RBH database on dermatological recipes)

| txtRecSig | txtRecTyp | txtCitedUse | txtCombinedUse | txtAdminRoute |
|---|---|---|---|---|
| 131b,126r.04 | Simple herbal | wurm am finngerr | wurm am finnger | Topical (external) |
| 131b,126r.06 | Simple herbal | für denn wurm | wurm am finnger | Topical (external) |
| 131b,126r.07 | Composed herbal | für denn wurm, So er nitt sterben will | wurm am finnger | Topical (external) |
| 131b,126r.08 | Composed herbal | für denn wurm | wurm am finnger | Topical (external) |
| 131b,126r.09 | Composed herbal | für denn wurm | wurm am finnger | Topical (external) |
| 131b,126r.10 | Composed herbal | zum wurm | wurm [allgemein] | Topical (external) |
| 131b,126r.11 | Composed mixed | für denn wurm | wurm [allgemein] | Topical (external) |
| 132a,126v.02.v01 | Composed mixed | wurm | wurm [allgemein] | Topical (external) |
| 132a,126v.04 | Simple herbal | Würm bÿss zeheillenn | Würm bÿss, schlanngen biss, nater biss | Topical (external) |
| 132a,126v.05 | Simple herbal | Würm bÿss, nater biss | Würm bÿss, schlanngen biss, nater biss | Topical (external) |
| 134a,128v.03 | Simple herbal | zitter mall: mossen vnnd flecken | zitter mall (mossen vnnd flecken) | Topical (external) |
| 134a,128v.05 | Simple herbal | zitter mall | zitter mall (mossen vnnd flecken) | Topical (external) |
| 134a,128v.06 | Composed mixed | zitter mall | zitter mall (mossen vnnd flecken) | Topical (external) |
| 134a,128v.08.v01 | Composed mixed | zitter möller vnnd fläckenn | zitter mall (mossen vnnd flecken) | Not mentioned |
| 134a,128v.08.v02 | Composed mixed | rud | rud | Not mentioned |
| 134a,128v.08.v03 | Composed mixed | verthribt all ... gifftig stich der spinen oder würmenn | gifftig stich der spinen oder würmenn | Not mentioned |
| 134a,128v.11 | Composed mixed | Böss fläckenn zU vertriben | zitter mall (mossen vnnd flecken) | Topical (external) |
| 134b,129r.01 | Composed herbal | Fläckenn | fleckenn | Topical (external) |
| 134b,129r.02 | Simple herbal | wunden die full ist, die macht sÿ rein | zU fullen wunden, bewertt vor fullem fleisch | Topical (external) |
| 136a,130v.01 | Composed herbal | Branndt Löschenn | Branndt | Topical (external) |
| 136a,130v.02.v01 | Simple herbal | Sannt anthonis fürr | Sannt anthonis fürr | Topical (external) |
| 136a,130v.02.v02 | Simple herbal | schwartzen blatteren | Schwartz blatteren | Topical (external) |
| 136a,130v.02.v03 | Simple herbal | heillet schlanngen vnnd scorpionn biss | Würm bÿss, schlanngen biss, nater biss | Topical (external) |
| 136a,130v.03 | Simple herbal | branndt | Branndt | Topical (external) |
| 136a,130v.04 | Composed mixed | branndt, es sige mitt wasserr, oder mitt fürr | Branndt | Topical (external) |
| 136a,130v.05 | Composed herbal | branndt | Branndt | Topical (external) |
| 136a,130v.08 | Composed mixed | branndt | Branndt | Topical (external) |

**SM – Table S1 – Recipes** (Extract from the RBH database on dermatological recipes)

| txtRecSig | txtRecTyp | txtCitedUse | txtCombinedUse | txtAdminRoute |
|---|---|---|---|---|
| 136b,131r.01 | Composed mixed | grosse hitz zU eim schadenn schlecht die der hitz hefftig werdt | Branndt | Topical (external) |
| 136b,131r.02 | Composed herbal | Branndt | Branndt | Topical (external) |
| 136b,131r.03 | Simple herbal | für alle gschwulst | geschwulst | Topical (external) |
| 136b,131r.04 | Simple herbal | hitziger schadenn | Branndt | Topical (external) |
| 136b,131r.05 | Simple herbal | hitziger schadenn | Branndt | Topical (external) |
| 156a,150v.01 | Composed herbal | Dornn vnnd pfill vss zU ziechenn | Dornn vnnd pfill vss zU ziechenn | Oral (ingested) |
| 156a,150v.02 | Composed herbal | Dornn vnnd pfill vss zU ziechenn | Dornn vnnd pfill vss zU ziechenn | Topical (external) |
| 156a,150v.03 | Composed herbal | Dornn vnnd pfill vss zU ziechenn | Dornn vnnd pfill vss zU ziechenn | Topical (external) |
| 156a,150v.06 | Simple herbal | Pfill vs Ziechenn | Dornn vnnd pfill vss zU ziechenn | Topical (external) |
| 156a,150v.08 | Composed mixed | dornn vnnden in fUss | Dornn vnnd pfill vss zU ziechenn | Topical (external) |
| 156a,150v.09 | Simple herbal | ein herrlich atraitium dörnn sprisen, spinlen spitz vs ze züchen | Dornn vnnd pfill vss zU ziechenn | Topical (external) |
| 156b,151r.02 | Simple herbal | an einn ÿssenn trädten, es zücht herus | Dornn vnnd pfill vss zU ziechenn | Topical (external) |
| 156b,151r.03 | Simple herbal | Ob einner an einn ÿssenn trädten, es zücht herus | Dornn vnnd pfill vss zU ziechenn | Topical (external) |
| 160a,154v.01 | Composed herbal | Erbgrinndt | erbgrinndt | Topical (external) |
| 160a,154v.02 | Composed mixed | rott louffennde grinnd | grindt | Topical (external) |
| 161b,156r.01 | Composed mixed | das eim das glid Wasser Verstadt | glid Wasser Verstadt | Topical (external) |
| 161b,156r.02 | Composed mixed | das eim das glid Wasser Verstadt | glid Wasser Verstadt | Topical (external) |
| 161b,156r.03.v01 | Composed mixed | das eim das glid Wasser | glid Wasser Verstadt | Topical (external) |
| 161b,156r.03.v02 | Composed herbal | das eim das glid Wasser Verstadt | glid Wasser Verstadt | Topical (external) |
| 161b,156r.06 | Composed mixed | das eim das glid Wasser Verstadt | glid Wasser Verstadt | Topical (external) |
| 161b,156r.07 | Composed mixed | das eim das glid Wasser Verstadt | glid Wasser Verstadt | Topical (external) |

**Dermatological plants and their uses in the *Receptarium* of Burkhard III from Hallwyl (RBH) from 16th century Switzerland – Data mining a historical text and preliminary *in vitro* screening –** Jonas Stehlin, Ina Albert, Thomas Frei, Barbara Frei Haller, Andreas Lardos – February 2024.

**SM – Table S2 – Assessment of RBH plant names, category A** (Extract from the RBH database on dermatological recipes / Reference list, see last page)

| Plant name RBH | ID | Plant name Idiotikon | Sci name Idiotikon | Reference Idiotikon | Plant name Marzell | Sci name Marzell | Reference Marzell | Plant name Fischer | Sci name Fischer |
|---|---|---|---|---|---|---|---|---|---|
| agrimonienn (klein vnnd gross) | 64 | agermönli, argemoni | Agrimonia eupatoria L. | 1.127 | Argemoni, Agrimoni | Agrimonia eupatoria L. | 1.139 | agrimonia | Agrimonia eupatoria L. |
| alat, alatt | 125 | [Not found] | -- | -- | Alant, Alat, Alatwurz | Inula helenium L. | 2.1012 | alantwurz | Inula helenium L. |
| albunn grecum | 121 | Perhaps corresponding to "weyss hirtzwurtz" = album graecum | Laserpitium latifolium (Libanotis alba) | 16.1734 | Weisse Hirschwurz? | Laserpitium latifolium L. (Libanotis alba, Cervariae albae, Weisser Enzian) | 2.1180 | [Not found] | -- |
| äpfen distell | 112 | [Not found] | -- | -- | [Not found] | -- | -- | [Not found] | -- |
| Aristologia rottunda | 173 | [Not found] | -- | -- | [Not found] | -- | -- | aristologia rotunda | Artistolochia pallida Willd. |
| aronenn | 156 | Aron, aronen | 1. gefleckter Aron, arum maculatum, sehr beliebtes Volksheilmittel gegen Lungenkrankheiten; 2. Alpenhahnenfuss, ranunculus alpestris | 1.388 | aron, aurone, arone; arone | Arum maculatum L.; Ranunculus alpestris L. ? | 1.443; 3.1238 | aaron, aranea | Arum maculatum L. |
| attich | 19 | Attichwurz, Attichholder | Zwergholunder, Sambucus ebulus | 16.1726 | Attich | Sambucus ebulus L. | 4.58 | attich | Sambucus ebulus L. |
| bach bumlen | 144 | Bachbumbele, Bachbumele | 1) Veronica beccabunga L. 2) Caltha palustris L., 3) Trollius europaeus L., 4) Schoenoplectrus lacustris (syn. Scirpus lacustris) - nur 1 Eintrag | 4.1259 | Bachbummel, Bachbumbel, Bachbumele | 1) Veronica beccabunga L. 2) Caltha palustris L., Geum rivale L., Petasites hybridus L., Trollius europaeus L. (als Übertragung von V. beccabunga) | 1.750, 2.681, 4.830, 1058 | bungen | Veronica beccabunga L. |
| barbenn | 63 | [Not found] | -- | -- | Barbenkraut, Barbara-, barbune; Barbenkraut, Barbara- | Achillea millefolium L. (1); Barbarea vulgaris R.Br. ? (2) | 1.84, 538 | [Not found] | -- |
| berÿ | 175 | [Not found] | -- | -- | [Not found] | -- | -- | [Not found] | -- |
| blauw Viönnlinn | 133 | Violen; Viöndli | Veilchen (Produkte aus deren Blütenblätter und Kraut, Art nicht näher definiert); Iris florentina L.: Es sind blaue | 16.1726 | Blauveilchen, Blauer Veil; Blaue Viole | Hesperis matronalis L.; Viola odorata L. | 2.842; 4.1169 | uioln | Viola odorata L. |

**SM – Table S2 – Assessment of RBH plant names, category A** (Extract from the RBH database on dermatological recipes / Reference list, see last page)

| Plant name RBH | ID | Plant name Idiotikon | Sci name Idiotikon | Reference Idiotikon | Plant name Marzell | Sci name Marzell | Reference Marzell | Plant name Fischer | Sci name Fischer |
|---|---|---|---|---|---|---|---|---|---|
| | | | und weisse Variätenten erwähnt, von beiden wird der Wurzelstock verwendet (vieliwurtz). Ein Synonym ist Vigelwurtz* | | | | | | |
| bonnen | 172 | Bonen-bluest /Bonen-bluest-wasser | Bohnen: Bohnenblüten / Bohnenblütenextrakt oder Aqua Hyssopi | 4.1315/16.1827 | Bohnen | Phaseolus vulgaris L., Vicia faba L. | 4.1122 | bonen | Vicia faba L. |
| boum öll, baum öll, boumöll | 67 | Baumöl | Keine Angaben zu Identität bzw. Stammpflanze | 1.182 | Ölbaum | Olea europaea L. | 3.378 | [Not found] | -- |
| boumwollen | 42 | baumwulle | Baumwolle, gossipion | 15.1377 | Baumwolle | Gossypium spp. | 2.736 | baumwolle | Gossypium herbaceum L., gossypium bombax |
| breitt vnnd spitz wëgrich | 24 | Breit-Wegerich / Spitz-Wegerich | Plantago major bzw. media / Plantago lanceolata | 15.951 | Breitwegerich; Spitzwegerich | Plantago major L., P. media L.; P. lanceolata L. | 3.806, 815, 835 | wegebreite; spitzer wegereich | Plantago media L., P. major L.; Plantago lanceolata L. |
| breitten wëgrich | 59 | Breit-Wegerich | Plantago major bzw. media | 15.951 | Breit-Wegerich | Plantago major L., P. media L. | 3.816, 836 | wegebreite | Plantago media L., P. major L. |
| Brun bethonienn, Brunne Betonicken | 18 | Brune Bettonien (siehe unter "brun") | Betonica off. | 5.647 | Braun Betonien, Betonica | Stachys officinalis (L.) Trevis. (S. betonica Benth.) | 4.461 | rote und braune bethonien | Stachys officinalis (L.) Trevis. (Betonica officinalis L.), Stachys alopecuros (L.) Benth. (Betonica alopecurus L.) |
| brunen kressich, brunenn kressich, brun kressich | 28 | Brunne-Chresse | Brunnenkresse, nast. offic. | 3.851 | Brunnenkresse | Nasturtium officinale R.Br.; Cardamine amara L. (auch wilde oder bittere Brunnenkresse) | 3.302 | brunnkress | Nasturtium officinale R.Br. |
| bUch spick (mitt gälbenn blUmen) | 151 | Buechspick | eine Art Wald-Habichtskraut, zunächst Mauer-Habichtskraut, Hieracium murorum | 10.100 | Buchspick | Heracium murorum L. (Name bezieht sich darauf, dass die Art oft in Buchenwäldern wächst) | 2.858 | [Not found] | -- |
| camillien, cammillien | 8 | chamille, kamille | Gemeine Kamille, Matric. cham. | 3.256 | Kamille | 1) Matricaria chamomilla L., 2) | 3.66, 1.975 | camillenblumen | Matricaria chamomilla L. |

**SM – Table S2 – Assessment of RBH plant names, category A** (Extract from the RBH database on dermatological recipes / Reference list, see last page)

| Plant name RBH | ID | Plant name Idiotikon | Sci name Idiotikon | Reference Idiotikon | Plant name Marzell | Sci name Marzell | Reference Marzell | Plant name Fischer | Sci name Fischer |
|---|---|---|---|---|---|---|---|---|---|
| | | | | | | Chrysanthemum parthenium L. | | | |
| cardobenedickten | 14 | Karde-Benedikte | Kardobenediktenkraut, Cnic. bened. | 4.1289 | Kardobenedikten-kraut | Centaurea benedicta L. (Cnicus benedictus L.) | 1.1062 | cardus benedictus, benedicte | Centaurea benedicta L. (Cnicus benedictus L.) |
| deschel krutt | 41 | Täschli-Chrut | Capsella bursa pastoris L., Hirtentäschel | 12.1873 | Täschel-kraut, teschelkrut | Capsella bursa pastoris L. | 1.788 | deschenkraut, taschelkhrawt | Capsella bursa-pastoris (L.) Medik. |
| diptannana | 174 | Diptam | Dictamnus albus L. | 14 | Diptam, dictamme | Dictamnus albus L. | 2.125 | diptan, diptamnum | Dictamnus albus L., Origanum dictamnus L. |
| dragantz | 170 | Tragant | Tragant (auch Wirbelkraut), Astragalus, auch der daraus gewonnene Saft | 14.610 | [Not found] | -- | -- | dragant | Astragalus creticus Lam., A. strobiliferus Royle; Artemisia dracunculus L. |
| edle salbinen | 114 | Edel-Salbinen | Salvia off. | 7.819 | Edel-Salbey | Salvia officinalis L. | 4.44 | salveye | Salvia officinalis L. |
| eerenpriss, erenbriss, erenn briss | 84 | Ere(n)bris | Ehrenpreis - Veronica off., Veronica cham (1); Augentrost - Euphrasia off. (2) | 5.795 | Ehrenpreis, Erenpris | Veronica officinalis L. (1); V. chamaedrys L. (2) | 4.1078 | erenbris | Veronica officinalis L., V. hederaefolia L., V. arvensis L. |
| egelkrutt | 109 | Egel-Chrut | 1) Rundblättriger Sonnentau, Dros. rot.; 2) Pfennigkraut, Lys. num. | 3.887 | Egelkraut; Egelchrut | Lysimachia nummularia L., Ranunculus flammula L., Drosera rotundifolia L., Alisma plantago-aquatica L.; Juncus articulatus L. (J. lampocarpus), Ficaria verna Huds. (Ranunculus ficaria L.) | 1.192, 2.170, 1070, 1506, 4.1257, 1263 | egilgras | Lysimachia nummularia L., Polygonum aviculare L.* |
| eichen, eichin | 136 | Eiche | Eiche | 1.71 | Eiche | Quercus | 3.1209 | Eich | Quercus robur L. (Q. penduculata), Q. petraea (Matt.) Liebl. (Q. sessiliflora) |
| endiuien | 98 | endivie; endivienwasser | Endivie, Cichorie, Cichorium endivia; Endivienwasser = Arznei aus der Endivie | 1.312; 16.1807 | Endivie; Wilde Endivie | Cichorium endivia L.; Cichorium intybus L. | 1.988, 997 | endiuie | Cichorium endivia L. |

**SM – Table S2 – Assessment of RBH plant names, category A** (Extract from the RBH database on dermatological recipes / Reference list, see last page)

| Plant name RBH | ID | Plant name Idiotikon | Sci name Idiotikon | Reference Idiotikon | Plant name Marzell | Sci name Marzell | Reference Marzell | Plant name Fischer | Sci name Fischer |
|---|---|---|---|---|---|---|---|---|---|
| epich (safft) | 23 | eppich; eppichwurtz | Epheu, Hedera helix; Knolle des Sellerie, Apium graveolens | 1.365 / 16.1725 | Eppich; Eppich, Epich | Apium graveolens L.; Hedera helix L. | 1.355 | ephich, epphi; ebich, epich | Apium graveolens L.; Hedera helix L. |
| erdberre | 77 | Erdbeere, ersperi | Walderdbeere, Frag. vesca | 4.1463 | Erdbeere | Fragaria vesca L. | 2.459 | ertberi, ertberechrawt | Fragaria vesca L. |
| erdrauch | 148 | Erdrauch | Fumar. off. B. | 6.97 | Erdrauch | Fumaria officinalis L. | 2.506 | erdrauch | Fumaria offcinalis L. |
| Eschinne | 101 | Esch, Esche | Eschenbaum, Fraxinus excelsior | 1.568 | Esche | Fraxinus excelsior L. | 2.468 | esche | Fraxinus excelsior L. |
| firnnis | 119 | firnissieren | Etwas mit Firnis anstreichen | 1.1019 | Sandarak | Tetraclinis articulata (Vahl) Mast (Callitris quadrivalvis Vent.) | 1.729 | vernix, sandaraca | Tetraclinis articulata (Vahl) Mast (Callitris sp.) |
| flachs, lÿnn | 53 | Flachs, Lein | Falchs - die Kulturpflanze; "Lein-" für Produkte aus Flachs | 1.1165, 7.935 | Flachs, Lein | Linum usitatissimum L. | 2.1334 | flachs, lein | Linum usitatissimum L. |
| full boümen | 149 | Fulbaum, Faulbaum | a) Elsbeerbaum, Sorbus torm. b) Pfeifenstrauch, Philad. coron. c) Ahlkirsche, Prun. prad. d) Faulbaum, Rhamn. frang. e) Waldrebe, clem. vitalba | 4.1237 | Fulbaum, Faulbaum | Acer pseudoplatanus L.; Frangula alnus Mill.; Prunus padus L.; Sorbus aucuparia L.; Sorbus torminalis (L.) Crantz | -- | fulbaum | Frangula alnus Mill. (Rhamnus frangula L.) |
| fünnff fingerr krutt | 141 | Fünffingerchrut; Wasserfünffingerkraut | versch. Potentilla, mit 5-zähl. Bl., bes. kriech. F., P. reptans L., Frühlings-F., P. verna L. (P. neumanniana Rchb. od. P. grandiflora L.), goldblüt. F., P. aurea (sub-/alpin), Alpen-F., P. alpestris Haller f. (P. alpina) (sub-/alpin) ; Comarum pal. | 3.890; 3.890 | Fünffingerkraut | 1) Potentilla reptans L.; 2) Potentilla erecta (L.) Raeusch.; 3) Comarum palustre L. (Potentilla palustris (L.) Scop.) = Rotes- / Wasser-Fünffingerkraut | 3.1015, 1023, 1.1114 | Fünffingerkraut | Potentilla reptans L., P. recta L., P. anserina L. |
| für blUmen | 116 | Fürbluem, Fürblüemli, Fürblume | 1) Klatschrose, Papver rhoeas; 2) Sandmohn, Papaver aregmone; 3) Feuerlilie, Lilium bulbiferum; 4) Mehlige Schlüsselblume, Prim. far. | 5.72 | Feuerblume, Fürblume | Papaver rhoeas L., P. dubium L., P. argemone L. (1); Silene dioica (L.) Clairv. (Melandrium rubrum), Silene flos-cuculi (L.) Clairv. (Lychnis flos-cuculi) (2); Adonis aestivalis | 1.119, 2.1450, 1292, 3.112, 538 | [Not found] | -- |

**SM – Table S2 – Assessment of RBH plant names, category A** (Extract from the RBH database on dermatological recipes / Reference list, see last page)

| Plant name RBH | ID | Plant name Idiotikon | Sci name Idiotikon | Reference Idiotikon | Plant name Marzell | Sci name Marzell | Reference Marzell | Plant name Fischer | Sci name Fischer |
|---|---|---|---|---|---|---|---|---|---|
| | | | | | | L., Lilium bulbiferum L., Primula farinosa L. (3) | | | |
| Gallöpffell | 49 | Gallepfel | Keine Angaben | 1.381 | Galläpfel-Eiche | Quercus infectoria G.Olivier | 3.1211 | [Not found] | -- |
| ganfer, ganfferr | 95 | Gamfer | Kampher | 2.310 | Kampfer, Kampferbaum | Cinnamomum camphora (L.) J.Presl, Dryobalanops aromatica C.F.Gaertn. | 1.1006, 2.174 | camffer | Cinnamomum camphora (L.) J.Presl (C. laura L.), Dryobalanops aromatica C.F.Gaertn. (D. camphora Colebr.) |
| genist blUmen | 65 | [Not found] | -- | -- | Genister, Gester | Cytisus scoparius (L.) Link | 4.112 | Genista | Spatium junceum L.; Cytisus scoparius (L.) Link (Spartium scoparium L.) |
| Gilge, wis gilge, wÿss Gilge | 86 | Wiss-ilge, Wissi ilie, gilge, Gilgenöl | Weisse Lilie, Lilium candidum; | 1.179, 181 | Gilge, Wiss Gilgen, Wissi Ilge, Wissilge | Lilium candidum L. | 2.1298 | liligen, weiss lilien | Lilium candidum L. |
| glariett, gloria, glorÿ | 93 | Gloriat, Glorie, Glorja, Glori | 1) Lerchenharz, Resina Laricea, vulgo Terebinthina; 2) "Harz" von Kirschenbäumen, "Gummi" | 2.642 | gloriet, loriet, gloriet-baum | Lerchenharz von Larix decidua Mill. | 2.1178 | [Not found] | -- |
| glost krutt | 61 | [Not found] | -- | -- | Glaskraut | Parietaria officinalis L. | 3.572 | [Not found] | -- |
| Gotz gnad, Gotz gnadt | 100 | Gott(e)sgnad; Gottes-Gnaden | 1) Stinkender Storchenschnabel, Ger. robert.; 2) Gratiola off.; 3) Corydalis cava (in Standort und Geruch ähnlich Ger. robert.); 3) Knautia longifolia (Scabiosa longifol.)*; Storchenschnabel-Arten, bes. Wald- und Wiesen-St., ger. silv., ger. prat.; | 2.661, 3.893 | Gott(e)sgnad, Gottesgnad | Gratiola officinalis L.; Geranium robertianum L.; Geranium pratentis L. (Übertragung von G. robert.); Corydalis cava (Übertragung von G. robertianum da ähnlicher Standort und Geruch); Scabiosa columbaria L. oder Sedum acre L. (als Übertragung ersterer). | 1.1197, 2.658, 667, 738, 4.152 | gottesgnad | Geranium robertianum L., G. sanguineum L., G. pratense L., G. molle L. |

**SM – Table S2 – Assessment of RBH plant names, category A** (Extract from the RBH database on dermatological recipes / Reference list, see last page)

| Plant name RBH | ID | Plant name Idiotikon | Sci name Idiotikon | Reference Idiotikon | Plant name Marzell | Sci name Marzell | Reference Marzell | Plant name Fischer | Sci name Fischer |
|---|---|---|---|---|---|---|---|---|---|
| grünne betonien | 106 | [Not found] | -- | -- | betonien | Stachys officinalis (L.) Trevis. (S. betonica Benth.) | 4.462 | betonien | Stachys officinalis (L.) Trevis. (Betonica officinalis L.), Stachys alopecuros (L.) Benth. (Betonica alopecurus L.) |
| gUtter heinrich | 82 | [Not found] | -- | -- | Guter Heinrich | Chenopodium bonus-henricus L. | 1.937 | guet heinrich | Chenopodium bonus-henricus L. |
| haber neslen, kleinn weg neslenn, kleiner nesslen | 9 | gross nesslen und klein, habernesslen | Urtica | 4.806 | Habernessel, Habernesslen, kleine brennende Nessel | 1) Urtica urens L., 2) Galeopsis tetrahit L. | 2.550 | habernessel, cleyn brenndende nessel | Urtica urens L. |
| haber, haberr | 85 | haber | Hafer, Avena sativa | 2.930 | haber | Avena sativa L. | 1.531 | habero | Avena sativa L. |
| hannff | 55 | Hanf | Hanfpflanze | 2.1437 | Hanf | Cannabis sativa L. | 1.775 | hanf | Cannabis sativa L. |
| hartz, wÿss büll hartz, rott büll hartz, gelüterett hartz* | 57 | Harz; Bül-Harz | Harz, Stammpflanze nicht definiert; Beulharz: Harzbeulen an den Stämmen von "Tannen", besonders von "Weisstannen". | 2.1654; 2.1655 | Harz; Beulenharzbaum | Harzbaum, Harztanne: Pinus sylvestris, Picea abies (P. excelsa); Edler Harz: Pinus cembra | 5.195 | [Not found] | -- |
| haslen | 40 | hasle, hasel, hassel | hasel, hasle = Haselnussstrauch | 3.1673 | Hasel, Hasle | Corylus avellana L. | 1.1199 | hasel | Corylus avellana L. |
| heidisch wund krutt | 157 | wundchrut, heidnisch | Wundklee, Anthyllis vulneraria | 3,951 | Heidnisch Wundkraut, Heydnisch Wundkraut, Heidnisch Wundchrut | Anthyllis vulneraria L., Solidago virgaurea L., Senecio ovatus Willd., Actaea spicata L., Hieracium murorum aggr., Mycelis muralis (L.) Dumort (Lactuca muralis (L.) Fresen.), Saxifraga rotundifolia L., Scrophularia nodosa L., Senecio hercynicus Herborg (S. nemorensis L.) | 1.114, 343; 2.859, 1145; 4.144, 188, 268, | -- (keine Einträge für "heidnisch"); I58wundcrut; wuntcruth; wundchrawt | -- / Aguja reptans L.; Prunella vulgaris L.; Silene dioica (L.) Clairv. (Melandryum rubrum Garcke.) |
| heidnisch wundkrutt (mitt den | 72 | [Not found] | -- | -- | [Not found] | -- | -- | [Not found] | -- |

**SM – Table S2 – Assessment of RBH plant names, category A** (Extract from the RBH database on dermatological recipes / Reference list, see last page)

| Plant name RBH | ID | Plant name Idiotikon | Sci name Idiotikon | Reference Idiotikon | Plant name Marzell | Sci name Marzell | Reference Marzell | Plant name Fischer | Sci name Fischer |
|---|---|---|---|---|---|---|---|---|---|
| zer schnitnen bletterenn)[1] | | | | | | | | | |
| Heill aller wëltt | 70 | [Not found] | -- | -- | Heil aller Welt, Heyl aller Welt | Agrimonia eupatoria L.; Anagallis arvensis L., Bupleurum falcatum L., Geum urbanum L., Sanicula europaea L., Veronica officinalis L. | 1.142, 258, 697; 2. 685; 4.102, 1080 | hayl aller welt | Agrimonia eupatoria L. |
| hirtzen zungen | 54 | [Not found] | -- | -- | Hirz-zunge, Hirschzunge | Asplenium scolopendrium L. (Phyllitis scolopendrium, Scolopendrium vulgare) | 3.704 | hirtzzungen | Scolopendrium vulgare Smith (= vrmtl. Scolopendrium officinare >> Aslpendium scolopendrium L.) |
| holder | 13 | holder | Gemeiner Holunder, Samb. nigra | 2.1183 | Hölder, Holder | Sambucus nigra L. | 4.64 | holder | Sambucus nigra L. |
| holtz öpffell | 139 | holzepfel, holzöpfel | 1) Frucht des wilden Apfelbaumes; 2) verschiedene veredelte Apfelsorten | 1.370 | holzapfel, holzöpfel; holzappel | 1) Malus communis ssp. silvestris; 2) Die Zapfen von Picea abies (L.) H.Karst (Picea excelsa, P. abies), unsicher, nur 2 Einträge aus D. | 3.25; 3.739 | holzapfel | Malus sylvestris Mill. (Pirus acerba H.C.) |
| hünner darm | 105 | hüenerdarm | ein kraut, alsine, corchorus; a) Stellaria media (alsine media), b) Anagallis arvensis (corchorus), c1) Veronica arvensis (Ehrenpreis), c2) Veronica agrestis (Eisenkraut), d) Cerastium latifolium | 13.1602 | Hühnerdarm, hüenerdarm | 1) Stellaria media (L.) Vill.; 2) Anagallis arvensis L. (syn. Lysimachia arvensis (L.) U.Manns & Anderb.); 3) Veronica hederifolia L., V. agrestis L., V. persica Poir.; Cerastium fontanum Baumg. s.l. (incl. C. caespitosum Gilb.) | 1.257, 897, 4.498 | hunesdarm, gruen hünerdärm; hunerdärm | Stellaria media (L.) Vill.; Anagallis avensis L. |

**SM – Table S2 – Assessment of RBH plant names, category A** (Extract from the RBH database on dermatological recipes / Reference list, see last page)

| Plant name RBH | ID | Plant name Idiotikon | Sci name Idiotikon | Reference Idiotikon | Plant name Marzell | Sci name Marzell | Reference Marzell | Plant name Fischer | Sci name Fischer |
|---|---|---|---|---|---|---|---|---|---|
| hus wurtzen | 80 | huswurz | Dachhauswurz, Sempervivum tect. (1); Trauben-Steinbrech, Saxifraga aizoon (2) | 16.1735 | hauswurtz, huswurz | Sempervivum tectorum L. (1), Sedum telephium L., Saxifraga azoides L. (2) | 4.135, 228, 246 | huswurtz | Sempervivum tectorum L. |
| jmber, ÿmberr | 69 | imber | Ingwer, amomum zingiber offic., bzw. dessen Wurzelstöcke | 1.236 | Ingwer, imber | Zingiber officinale Roscoe | 4.1244 | ymber | Zingiber officinale Roscoe (Zingiber amomum L. ?) |
| juden öpffell | 132 | Judenepfel, Judenapfel | auch Lederapfel: "verschiedene Rainettenarten mit lederartiger Haut" (Apfelsorte) | 1.370, 372 | Judenäpfel | Citrus medica L. | 1.1032 | Judenapfel | Citrus medica L. |
| kärnenn | 110 | Chërne | Dinkel | 3.466 | Kern, Cherne | Triticum aestivum subsp. spelta (L.) Thell. (Triticum spelta L.), Dinkel | 4.817 | [Not found] | -- |
| katzen trübel | 62 | Chatzen-Trübel | Sedum, kleine Hauswurz, Petrosedum pruinatum (Link ex Brot.) Grulich (Sedum reflexum Cutanda) | 14.215 | Katzentreubel (männlein, weiblein); Katzentraube, Katzenträubel | Sedum acre L., S. album L. (1); Muscari racemosum (L.) Mill., M. neglecta aggr., M. botryoides (L.) Mill. ?, M. comosum (L.) Mill. ? (2) | 3.225, 4.202, 216 | Katzentrübel | Sedum acre L. |
| kernn gertten | 115 | Chern-Gerte, kerngerten | Verschiedene Sträucher, grösstenteils mit zähen Zweigen, Hartrigel/Cornus, Ligustrum | 2.441 | Kerngerte, Kelgerte, Chorngerte (rot, weiss, schwarz) | Ligustrum vulgare L., Cornus sanguinea L. (1); Viburnum lantana L., V. opulus L., Frangula alnus Mill. Rhamnus cathartica L. (2) | 1.1179, 2.483, 1285, 4.1099, 1111, 1307 | [Not found] | -- |
| klein schwerttel wurtz | 176 | Schwertelwurz | Knolle verschiedener (Schwert-)Lilien-Arten | 16.1753 | [Not found] | -- | -- | [Not found] | -- |
| klein winter grün | 73 | Wintergrün | Verschiedene im Winter grüne Pflanzen: a) Immergrün, Vinca minor; b) Bärlapp; c) Mistel, Viscum album; d) Gnaphalium marg.; e) | 2.753 | Wintergrün | Pyrola minor L., Pyrola spp., Vinca minor L. (1); Polygala chamaebuxus L., Lyopodium clavatum | 3.868, 1195; 4.1144 | wintergruen der chlanz | Pyrola rotundifolia L., P. secunda L. |

**SM – Table S2 – Assessment of RBH plant names, category A** (Extract from the RBH database on dermatological recipes / Reference list, see last page)

| Plant name RBH | ID | Plant name Idiotikon | Sci name Idiotikon | Reference Idiotikon | Plant name Marzell | Sci name Marzell | Reference Marzell | Plant name Fischer | Sci name Fischer |
|---|---|---|---|---|---|---|---|---|---|
| | | | Mercurialis; f) Limonium, pirola | | | L. (2) - all are "small" plants referred to as "Wintergrün", there are however no specific entries for "Kleines Wintergrün". | | | |
| knaben krutt | 140 | Knabenkraut, Chnabechrut | Orchis | 3.898 | Knabenkraut, knabenkrut; Knabenkraut | Orchis mascula L., O. militaris L., Dactylorhiza maculata (O. maculata L.), D. incarnata ssp. incarnata (O. latifolia L.), O. morio L., O. ustulata L., Orichis spp.; Gymnadenia conopsea (L.) R.Br.; Platanthera bifolia (L.). Rich., Sedum telephium L. | 3.422; 4.229 | Knabenkraut | Orchis spp. |
| knoblach | 153 | Chnoblauch, Knoblouch | 1. Knoblauch 2. Trauben-Bisamhyacinthe, musc. rac. {Moschustraubenhyazinthe} | 3.1006 | knoblauch; wilder knoblauch, chnoblich | Allium sativum L. (1); Allium ursinum L. (2) | 1.204, 211 | chlobluch, knoblauch | Allium sativum L. |
| knoblach krutt | 146 | Knob-Lauch-Chrut, Chnoblauchchrut | Knoblauchsgamander, Teucr. scord. (unter "Fieber-Chrut", 2. Synonym) | 3.899 | Knoblauchkraut, Chonblauch-Chrut | Alliaria petiolata (M.Bieb.) Cavara & Grande (A. officinalis Andrz.); Teucrium scordium L. | 1.194 | [Not found] | -- |
| korble krutt, korblin krutt | 37 | Chröblichrut, körble-, körbli-, körblinkrut | Gartenkerbel, Anthr. caer. | 3.897 | Körblinkraut, Körbelkraut, Körblekraut, Kerbel | Anthriscus cerefolium (L.) Hoffm. | 1.330 | chervenchraut | Anthriscus cerefolium (L.) Hoffm. |
| küngund krutt | 32 | [Not found] | -- | -- | Kunigundenkraut, Kündgundkraut | 1) Eupatorium cannabinum L. (Kundigundkraut menlin); 2) Bidens cernuus L. (Kunigundkraut weible) oder B. | 1.603, 2.360 | chumguntkraut | Eupatorium cannabinum L. |

**SM – Table S2 – Assessment of RBH plant names, category A** (Extract from the RBH database on dermatological recipes / Reference list, see last page)

| Plant name RBH | ID | Plant name Idiotikon | Sci name Idiotikon | Reference Idiotikon | Plant name Marzell | Sci name Marzell | Reference Marzell | Plant name Fischer | Sci name Fischer |
|---|---|---|---|---|---|---|---|---|---|
| | | | | | | tripartitus L., wachsen am gleichen Standort wie Eupatorium cannabinum. | | | |
| kütenenn | 169 | Chütenen-, Chüten-, Küttenen-, Kütten- | Quitten- | 4.1240 | Chutina, Kutina, Quitte | Cydonia oblonga Mill. | 1.1289 | chutten, quitten, kütten | Cydonia oblonga Mill. (C. vulgaris Pers.) |
| lennger ie lieber | 16 | Längeri lieber (Je länger je lieber) | 1) Bittersüss, Sol. dulc.; 2) Geissblatt: a) Lonic. capr./pericl.; b) Sauerdorn, Berb. vulg. | 3.991 | Je länger je lieber | 1) Solanum dulcamara L., 2) Lonicera periclymenum L., L. caprifolium L. | 2.1376, 4.358 | je länger je lieber | Solanum dulcamara L.; Veronica chamaedrys L. |
| liliенn (krutt) | 164 | Lilie / Ilgen-wasser, Lilien-wasser | Lilie: 1. Lilie 2. Bergveilchen, bzw. gesporntes V., viola alp. et calc. 3. Frühlingsenzian, gent. verna / Lilienwasser: Arznei aus Blütenblättern der Lilie; in Essig eingelegt Blütenblätter der Lilie, die auf Wunden gelegt wurden | 3.1260 / 16.1807 | Lilie, Lilge; Lilie; (Berg-)Lilie | Lilium candidum (1); Iris germanica (2); Viola calcarata (3): unwahrscheinlich, da Alpenpflanze und meist als "Berg-Lilie/-Veilchen angesprochen. | 2.1296 / 2.1026 / 4.xxx | lilge, lilium | Lilium candidum L. |
| linnden | 168 | Linde | die Linde, tilia, und zwar sowohl die grossblättrige, t. grand., als die kleinblättrige, t. parv., meist als «Summer-» und «Winter-L.» unterschieden | 3.1319 | Linde | Tilia spp. | 4.718 | linte | Tilia spp. |
| Lor-, lorberr | 7 | Lorbone (Lor-) | Lorbeer | 4.1313 | Lorbe, Lorbone | Laurus nobilis L. | 2.1209 | lorboum, lorpir | Laurus nobilis L. |
| lüss krutt | 159 | Lüs-chrut | Sumpf-Läusekraut, ped. pal.; Niesswurz, bes. hell. vir. und hell. foet.; scharfer oder Läuse-Rittersporn, delph. staph.; Haarmoos, pol. vul. | 3.900 | Läusekraut | Delphinium staphisagria L. (nicht in CH), Daphne mezereum L., Helleborus foetida L. (hist. Texte); Lycopodium clavatum L. / selago L. (in CH selten, v.a. Alpentäler), Pedicularis palustris | -- | lusskrut | Delphinium staphisagria L., Pedicularis sp. |

**SM – Table S2 – Assessment of RBH plant names, category A** (Extract from the RBH database on dermatological recipes / Reference list, see last page)

| Plant name RBH | ID | Plant name Idiotikon | Sci name Idiotikon | Reference Idiotikon | Plant name Marzell | Sci name Marzell | Reference Marzell | Plant name Fischer | Sci name Fischer |
|---|---|---|---|---|---|---|---|---|---|
| | | | | | | L., Polytrichium commune L., Polypodium vulgare L. (nx CH, D, CS) | | | |
| macis | 130 | Mazis | macis, Muskatblüte | 4.601 | Macisnuss | Myristica fragrans Houtt. | 3.264 | macis | Myristica fragrans Houtt. (M. moschata Thunb.) |
| mandel | 96 | mandel | Mandel, nux graeca | 4.319 | Mandel | Prunus amygdalus Batsch (P. dulcis (Mill.) D.A.Webb) | -- | mandilboum | Prunus amygdalus Batsch (P. dulcis (Mill.) D.A.Webb) |
| mass blümlin, moss blUmen | 66 | Mass-Blüemli | Bellis perennis, Massliebchen | 5.83 | Maasblüemli; Massblume | Bellis perennis L. (1), Papaver sominferum L. ? (2) | 1.547, 3.562 | masslieben | Bellis perennis L. |
| mastix | 87 | Mastix-Pflaster | Emplastrum mastichum | 2.1261 | ? | -- | -- | mastix | Pistacia lentiscus L. |
| meister wurtz | 27 | Meisterwurz, Meisterwurtz | 1) Peucedanum (Imperatoria) ostruthium, 2) Astrantia major = Schwarze Meisterwurtz | 16.1742 | Meisterwurz, Meisterwurtz | Peucedanum ostruthium | 3.644 | meisterwurz | Peucedanum ostruthium (L.) W.D.J.Koch (Imperatoria ostruthium L.) |
| melissen | 127 | Melisse | Gartenmelisse, Mel. off. | 4.171 | Melisse | Melissa officinalis L. | 3.132; 3.312 | melise | Melissa officinalis L. |
| mer rettich | 150 | Mer-rättich, Mer-rätterich, Meerrettig | Armoracia, Meerrettich | 6.1630 | Meerrettich, merretich | Armoracia rusticana G.M.Sch. | 1.395 | meretich, meredich | Aromracia rusticana G.Gaertn. B.Mey et Scherb. (syn. Cochlearia armoracia L.) |
| miess | 46 | Mies | Various mosses or lichens | 4.467 | Mies, Moos, maisz | Musci and tree lichens such as Usnea florida Wiggers | 3.237, 4.927 | miess | Usnea spp. (Usnea arborum) |
| modelgeer | 182 | madelger | Enzian, spec. g. cruciata, mit kreuzweise durchstochener Wurzel; 'g. minor' | 2.402 | madelger | Gentiana cruciata | -- | [Not found] | -- |
| müllj stoub | 137 | mülistaub | flour dust | 10.1070 | [Not found] | -- | -- | [Not found] | -- |
| muscat nus, muscatt | 94 | muschkardnuss, muschgetnuss | Muskatnuss | 4.828 | Muskatnuss | Myristica fragrans Houtt. | 3.263 | muscat, nux muscata | Myristica fragrans Houtt. (M. moschata Thunb.) |

**SM – Table S2 – Assessment of RBH plant names, category A** (Extract from the RBH database on dermatological recipes / Reference list, see last page)

| Plant name RBH | ID | Plant name Idiotikon | Sci name Idiotikon | Reference Idiotikon | Plant name Marzell | Sci name Marzell | Reference Marzell | Plant name Fischer | Sci name Fischer |
|---|---|---|---|---|---|---|---|---|---|
| nacht schatt, nachtschatten, nach schatten | 60 | Nacht-Schatten | Primär Solanum nigrum oder Solanum dulcamara. Es werden zum Teil jedoch auch andere Pflanzen unter diesem Namen angesprochen | 8.1493 | Nachtschatten | Solanum nigrum L. | 4.363 | nacht schade; nahtsate | Solanum nigrum L.; Solanum dulcamara L. |
| näptenn krutt | 163 | Neptechrut = Chatzechrut | 1. gem. Baldrian, val. off. 2. gem. Katzenmünze, nep. cat. 3. Waldmünze, mentha silv. {Rossminze} 4. Katzen-Gamander, teucr. marum | 3, 898 | Katzenkraut | Nepeta cataria 1; Teucrium marum 1; Trifolium arvensis 1; Ruta graveolens 2; Satureja hortensis 2; Geranium sp. 2; Malva slyvestris 3; Meum athamanticum 4; Menta longifolium 4; Valeriana officinalis 4; Mentha arvensis 5; Mentha piperita 5; und weitere | Various | napt, nepeta, katzenkraut | Nepeta cataria L; Clinopodium nepeta ssp. spruneri (syn. Melissa calamintha L.); Valeriana officinalis L. |
| natter krutt, natterkrutt | 71 | Naterchrut | Heildistel, Cnicus benedictus | 3.903 | Natterkraut, Natterchrut | Centaurea benedicta (L.) L. (Cnicus benedictus), Lysimachia nummularia L., Echium vulgare L. (1); Scorzonera hispanica L. (2); Lycopodium clavatum L. (3). Alle Pflanzen gelten als Wundkräuter und haben einen Bezug zu Schlangenbissen. | 1.1064, 2.183, 1479, 1504, 4.183 | Natherwurtz | Polygonum bistorta L. |
| nëgellin, näglin | 6 | Nägeli | Gewürz-Nelke, 1) Caryophyllum aromaticum, 2) Dianthus spp. und 3) "wilde Nelken" | 4.692 | Näglein, Nelke, negelin | 1) Gewürznelke, Eugenia caryophyllata, 2) Ab 16. Jh., aufgrund ähnlichen Geruchs der Nelken auf | 2.101 | negely, negelyn; negelein, negelly | Syzygium aromaticum (L.) Merr. et L.M.Perry (Eugenia caryophyllata Thunbg.); Dianthus spp. |

**SM – Table S2 – Assessment of RBH plant names, category A** (Extract from the RBH database on dermatological recipes / Reference list, see last page)

| Plant name RBH | ID | Plant name Idiotikon | Sci name Idiotikon | Reference Idiotikon | Plant name Marzell | Sci name Marzell | Reference Marzell | Plant name Fischer | Sci name Fischer |
|---|---|---|---|---|---|---|---|---|---|
| | | | | | | Dianthus spp. übertragen, inbesondere Dianthus caryophyllus L.. | | | |
| nesslen | 43 | Nessle(n) | Brennessel, sowohl grosse (U. dioica) wie kleine (U. urens). | 4.805 | Nesslen | Urtica dioica L., U. urens L. (1); Lamium spp. (L. album L., L. amplexicaule L., L. galeobdolon (L.) L., L. maculatum L., L. purpureum L.), Stachys sylvatica L. (2) | 2.1154, 4.475, 914, 922, | nessel; cleyn brennende nessel | Urtica dioica L.; Urtica urens L. |
| niess wurtz | 166 | nieswurz | weisse n. = Germerwurz; schwarze n. = Christwurz | 16.1744 | Nieswurz; Schwarze/Weisse Nieswurz | Helleborus niger; Veratrum album | 2.799; 4.1021 | niesewurze; niswurz | Helleborus niger, viridis, foetidus; Veratrum album |
| olibanum, olibanj [see "weÿrauch"] | 120 | [Not found] | -- | -- | [Not found] | -- | -- | olibanum | Boswellia sacra Flueck. |
| papell, babell, bapellenn krutt | 35 | Pappele, papelen, bapel | Malve: Wegmalve, Malva vulg.; Rundblättrige Malve, Malva rotund. | 4.1415 | Pappel, papel | Malva sylvestris L., M. neglecta Wallr.; Taraxacum officinalis L. | 1.231, 3.33, 4.650 | papele, pappel; papele | Malva sylvestris L.; Alcea rosea L. (Althaea rosea Willd.) |
| paris körnlin | 129 | Paris-Chorn | Samenkorn von amomum granum paradisi {Paradieskorn} | 4.1445 | Pariskorn, Pariskörner | Semen Paradisi von Aframomum melegueta K.Schum. (Amomum melegueta Rosc.) | 1.248 | [Not found] | -- |
| petterlj | 135 | peterli | 1) Peterli, Petr. sat.; 2) Übertragungen auf andere Pflanzen, die mehr oder weniger Ähnlichkeit mit der Gartenpetersilie haben* | 4.1842 | Peterle | Petroselinum crispum (Mill.) Fuss (Petroselinum hortensis) | 3.630 | petersill, petersilien | Petroselinum crispum (Mill.) Fuss (Petroselinum sativum Mill.) |
| pfäffer krutt | 184 | Pfefferchrut | 1. Garten-Pfefferkraut, sat. hort., 2. breitblättrige Kresse, lep. lat. | 3.905 | Pfefferkraut | Artem. drac. 2, Asarum europ. 4, Dictamn. alb. 9, Gypsophila pan., Lepidium latif., | -- | pfefferkraut, pfeffercrut | Lepidium latifolium L., Satureja hortensis L. |

**SM – Table S2 – Assessment of RBH plant names, category A** (Extract from the RBH database on dermatological recipes / Reference list, see last page)

| Plant name RBH | ID | Plant name Idiotikon | Sci name Idiotikon | Reference Idiotikon | Plant name Marzell | Sci name Marzell | Reference Marzell | Plant name Fischer | Sci name Fischer |
|---|---|---|---|---|---|---|---|---|---|
| | | | | | | Polygonum hydropip. 1, Salvia off. 5, Satureja hort 1, Sedum acre 7, Spilanthes oleracea, Tropaeolum majus 6 | | | |
| pflumen | 17 | pflume | Pflaume, Frucht des Pflaumenbaums, Prun. dom. | 5.1247 | pflume | Prunus domestica L. | 3.1111 | pflawm | Prunus domestica L. |
| poleÿ, boleÿ, hertzen bleich, hertz bleÿ | 5 | Polei, Hërze-Bleiche, Herz-Polei | Polei-Minze, Mentha pulegium | 5.8, 4.1181 | Polei, Herze-bleiche, Herz-Polei | Mentha pulegium L. | 3.163 | polei | Mentha pulegium L. |
| reb krutt | 10 | Rebchrut | 1) Gundelrebe, Glechoma heder., 2) Trauben-Gamander, Teucr. botrys, 3) Edler Gamander, Teucr. cham. | 3.906 | Rebechrut | 1) Teucrium botrys L., Traubengamander | 4.665 | [Not found] | -- |
| rörlin krutt | 160 | Rörlichrut | 1. gem. Löwenzahn, tar. off. 2. eine Art Cichorie, {Hedypnois} | 3.907 | Röhrleinkraut, Röhrlkraut, Röhrkraut, Röhrlichrut | Taraxacum officinalis L. | 4.607 | [Not found] | -- |
| rosenn wurtzell | 47 | rose(n)wurz | Rosenwurz, Rhodiola rosea | 16.1750 | Rosenwurz | Rhodiola rosea L. (Sedum roseum (L.) Scop.) | 4.222 | [Not found] | -- |
| rosenn, ross | 36 | ros, ross | Rosa spp. | 6.1385, 1391 | Rose | Rosa gallica L., R. x damascena Herrm., R. x centifolia L. | 3.1394, 1432 | rosen (wys, edel, gefüllt, zam) | Rosa x centifolia L., R. x alba L., R. villosa L., |
| rot bugglenn, rott buglenn, rott bugenn | 11 | Rot-Buggelen | 1) Grüner Fuchsschwanz, Amar. blitum; 2) Stumpfblättriger Ampfer, Rum. obt.; 3) Gemeiner Portulak, Port. ol; 4) Rote Abart von Art(emisia). vulg. | 4.1091 | Rotbuckel, Rote Buck, Rother Bock, Rot Buggele | 1) Artemisia vulgaris L., 2) Amaranthus blitum L. (syn. A. viridis L.) | 1.243, 436 | buggela | Artemisia vulgaris L., A. campestris L., A. genipi Weber ex Stechm. (A. campestris L.) |
| rotten mangolt | 75 | Rotmangel- | Mangold, beta | 4.328 | Rot-mangolt, Roter Mangolt | Beta vulgaris var. cicla L. | 1.583 | Mangolt, Rotrueben | Beta vulgaris L. (B. cicla) |
| rottenn thannen, rott danj | 68 | Rottanne | Fichte, Picea excelsa (pinus abies usw.) | 13.72 | Rottanne | Picea abies (L.) H.Karst. (Picea excelsa) | 3.272 | [Not found] | -- |

**SM – Table S2 – Assessment of RBH plant names, category A** (Extract from the RBH database on dermatological recipes / Reference list, see last page)

| Plant name RBH | ID | Plant name Idiotikon | Sci name Idiotikon | Reference Idiotikon | Plant name Marzell | Sci name Marzell | Reference Marzell | Plant name Fischer | Sci name Fischer |
|---|---|---|---|---|---|---|---|---|---|
| rutten | 33 | Rute, Rut | Gartenraute, Ruta grav. | 6.1797 | Raute | Ruta graveolens L. | 3.1552 | rautten | Ruta graveolens L. |
| saffrann | 118 | Saffran | 1) Offizineller Safran, Crocus sat.; 2) Wilder Safran, a) Färberdistel, Carthamus tinctorius, b) Herbstzeitlose, Colchic. aut. | 7.333 | safran | Crocus sativus L. | 1.1248 | saffran; wilder saffran; | Crocus sativus L.; Carthamus tinctorius L.; |
| safft, so vs dem holtz flüst das jm fürr ligt | 147 | [Not found] | -- | -- | [Not found] | -- | -- | [Not found] | -- |
| salbinenn | 103 | Salbei, Salbine | Salbei, Salvia offic.., S. prat. (wildi Salbine) | 7.817 | Salbine, Salbei | 1) Salvia officinalis L., 2) Salvia pratensis | 4.42, 48 | salveye; wilder Salbei | Salvia officinalis L.; S. pratensis L. |
| Sanickel | 45 | Sanikel | Sanicula europ. | 7.999 | Sanikel, sanicula mas | Sanicula europaea L. | 4.100 | sanikel | Sanicula europaea L. |
| sanndelholtz | 38 | Sandel | Sandelholz, santalum | 7.1116 | Sandelholz, Sandel holtz | Sandalum spp. | 4.103 | sandelholz | Santalum album L. |
| Sannt petters krutt | 29 | St. Peters-Chrut | 1) Glaskraut, Parietariae; 2) Mauerrauten, Muralis herba | 3.905 | -- | Parietaria officinalis L.; Succisa pratensis Moench ? (Teufelsabiss), Gentiana cruciata L. ? (St. Peterswurz) | 2.622, 3.574, 4.529 | sant peterskrut | Parietaria officinalis L. (P. erecta), P. judaica L. (P. ramiflora Moench, P. diffusa) |
| sanntt johanns krutt | 117 | Johannischrut, Sankt Johannskraut | 1) Gemeines Hartheut, Hyp. perf.; 2) Wermut, Artem. vulg.; 3) Schmerrwurz, grosse Fetthenne, Sedum telephium; 4) die Blätter der schwarzen Johannisbeere | 3.896 | Johanniskraut, Sankt Johannskraut | Hypericum perforatum L. (Weitere Interpretationen: Achillea, Actaea, Artemisia, Aruncus, Arnica mont., Aspidium, Chrysanthemum, Epilobium, Geum, Orchis, Salvia, Scleranthus, Scrophularia, Sedum, Senecio, Solidago, Stachys, Thalictrum, Thesium, Verbascum) | 2.942 | sant johannskraut | Hypericum perforatum L. |

**Dermatological plants and their uses in the *Receptarium* of Burkhard III from Hallwyl (RBH) from 16th century Switzerland – Data mining a historical text and preliminary *in vitro* screening –** Jonas Stehlin, Ina Albert, Thomas Frei, Barbara Frei Haller, Andreas Lardos – February 2024.

**SM – Table S2 – Assessment of RBH plant names, category A** (Extract from the RBH database on dermatological recipes / Reference list, see last page)

| Plant name RBH | ID | Plant name Idiotikon | Sci name Idiotikon | Reference Idiotikon | Plant name Marzell | Sci name Marzell | Reference Marzell | Plant name Fischer | Sci name Fischer |
|---|---|---|---|---|---|---|---|---|---|
| sant barbel krutt | 179 | [Not found] | -- | -- | Sant Barbaren krut | Barbaraea vulgaris R. Br. | 1.538 | [Not found] | -- |
| schellkrutt, schelkrut, schell krutt | 74 | schellkraut, schälkraut | Primär: Sedum acre, S. tel., S. alb., S. ann., S. rep., S. vill.; Sekundär: Schöllkrautblätter (Chelidonium); grosses Schöllchrut: Thalictrum aquil. | 3.909 | Schellkrut, Schellchrut | Chelidonium majus L., Sedum acre L., S. album L., S. telephium L., Sedum spp. (1); Rhinanthus alectorolophus (Scop.) Pollich ?, R. minor L. ?, Rhinanthus spp. ? | 1.184, 924; 4.213, 218, 230 | schelchrawt | Chelidonium majus L. |
| schlechenn, schlehen dornn | 34 | Schlehe, Schleche | Schlehe, Schwarzdorn, Prunus spin. | 9.499 | Schlehe | Prunus spinosa L. | 3.1153 | slehe | Prunus spinosa L.; Prunus domestica subsp. insititia (L.) Bonnier & Layens (P. insititia L.) - meist Haferschlehe |
| schlüssell blUmen | 183 | Schlüsselblueme | 1. verschiedene Primelarten Prim. elat., Prim. off., Prim. far., Prim. aur., Prim. vill.2. Soldan. alp. 3. Orchis mac., masc, 3) Coryd. cav. 5) Polygala cham. 6) Lotus cornic. | 5.88 | Schlüsselblume | Anem. nem. 13, Artem. laxa, Cardam. prat. 12, Coryd. cava 7, Gentiana bavarica, G. verna 6, Helleb. purpur., Hyacinth. s orient. 4, Lotus corn. 10, Orchis 8, Polyg. chamaebux. 2, Prim. aur. 7, Prim. elatior, veris 1a, Pulmon. off 9, Syringa vulg. 1 | -- | schlüsselblumen | Primula veris L. (P. officinalis L.) |
| schwalmenn wurtzell | 171 | Schwalmenwurz | Schwalbenwurz, Vincetoxicum hirundinaria | 16.1753 | Schwalmenwurz(el), Schwalbenwurz(el) | Vincetoxicum hirundinaria Med. | 4.1152 | schwalbenwurz | Vincetoxicum hirundinaria Medik. (Vincetoxicum asclepiadea) |
| schwartzen nacht schatten | 25 | Schwarzer Nachtschatten | Solan. nigr. | 8.1493 | Nachtschatten, Schwarzer Nachtschatten | Solanum nigrum L. (1), Scrophularia umbrosa Dumort. (2) | 4.195, 363 | nachtschade | Solanum nigrum L. |

**Dermatological plants and their uses in the *Receptarium* of Burkhard III from Hallwyl (RBH) from 16th century Switzerland – Data mining a historical text and preliminary *in vitro* screening –** Jonas Stehlin, Ina Albert, Thomas Frei, Barbara Frei Haller, Andreas Lardos – February 2024.

**SM – Table S2 – Assessment of RBH plant names, category A** (Extract from the RBH database on dermatological recipes / Reference list, see last page)

| Plant name RBH | ID | Plant name Idiotikon | Sci name Idiotikon | Reference Idiotikon | Plant name Marzell | Sci name Marzell | Reference Marzell | Plant name Fischer | Sci name Fischer |
|---|---|---|---|---|---|---|---|---|---|
| scritten [see "stritten"] | 155 | [Not found] | -- | -- | [Not found] | -- | -- | [Not found] | -- |
| seffe[2] | n.a. | sefi, sefibaum | Juniperus sabina | 7.34 | seffen, sefi | Juniperus sabina L., Calluna vulgaris Hull?, Juniperus communis var. saxatilis Pall. (J. nana)? | 2.1094, 1.173 | [Not found] | -- |
| sennff krutt | 180 | Senfchrut | 1. breitblättrige Kresse, lep. lat., zsgeworfen mit dem Senf, sinapis 2. 'wild s.', Rainkohl, laps. comm. | 3.908 | Senfkraut | Lepidium latifolium L. (n x), Barbaraea vulgaris R. Br. (4x D); Satureja hortensis L. (2x D) | 4.127 | [Not found] | -- |
| serpentia | 107 | [Not found] | -- | -- | serpentaria; serpentina, serpentaria maior/minor | Bistorta officinalis Delarbre (Persicaria bistorta (L.) Samp., Persicaria bistorta (L.) Samp.); Dracunculus vulgaris Schott, Arum maculatum L., A. italicum L. | 3.907 | serpentin, serpentaria | Bistorta officinalis Delarbre (Persicaria bistorta (L.) Samp., Polygonum bistorta L.), P. amphibia (L.) Delarbre (syn. Polygonum amphibium L.) |
| seueboum | 152 | Sefibaum, seuenboum, sauina | Sabina, Sevenbaum, Juniperus sabina | 4.1245 | savina, sabina, seuenboum | Juniperus sabina L. | 2.1094 | seuvenbom, seuvin, savina | Juniperus sabina L. |
| sigilis solomonis | 165 | Salomonssiegel | Maiblume, Weisswurz, Polygonatum odoratum (Mill.) Druce (Convallaria polygonatum) | 7.496 | Salomonssiegel, sigilum salamonis | Polygonatum odoratum (Mill.) Druce (P. officinale Moench) | 4.xxx | sigilum salamonis | Polygonatum odoratum (Mill.) Druce (P. officinale Moench), P. verticillatum (L.) All. |
| sinauw | 44 | Sinau | Alchemilla vulg. | 7.1085 | Sinau | Alchemilla xanthochlora Rothm. (A. vulgaris L.) | 1.178 | sinau | Alchemilla xanthochlora Rothm. (A. vulgaris) |
| spickennardj | 131 | Spi(c)knard(en) | Lavandula spica oder Valeriana spica? | 1.475 | spic(a)nard(en), spikennard; spica nardus germanica, spicanard, Spitzenart; narrenspeik, nardi spica | Nardostachys jatamansi (D.Don) DC.; Lavandula latifolia Medik. (L. spica var. latifolia L.); Valeriana celtica L. | 3.291; 2.1212; 4.986 | spica nardi; nardus spica, spica | Nardostachys jatamansi DC.; Lavandula latifolia Medik. |

**SM – Table S2 – Assessment of RBH plant names, category A** (Extract from the RBH database on dermatological recipes / Reference list, see last page)

| Plant name RBH | ID | Plant name Idiotikon | Sci name Idiotikon | Reference Idiotikon | Plant name Marzell | Sci name Marzell | Reference Marzell | Plant name Fischer | Sci name Fischer |
|---|---|---|---|---|---|---|---|---|---|
| spitzwegrich, spitzen wegrich, spitzen wägrich | 58 | Spitz-Wegerich | Plantago lanceolata | 15.951 | Spitz-Wegerich | Plantago lanceolata L. | 3.806 | spitzer-wegerich | Plantago lanceolata L. |
| stinckende nesslen | 154 | stink-chrut | No species cited | 3.912 | stink-nessel | Stachys silvatica L. (in der CH häufig, Wälder, Hecken); Ballota nigra L. (wärmeliebend, in der CH nur regional) | 1.537; 4.475 | [Not found] | -- |
| stritten | 161 | Strit, Strite, Stritele | a) Immergrün, Vinca minor b) von andern (kriechenden, rankenden) Pflanzen α) «gëlʷi Str.-eⁿ», Pfennigkraut, Lysimachia numm. β) «Balm-Str.», gestutzte Weide, Salix ret. | 11.2409 | Strit, Streit | Vinca minor; Lysimachia numularia (=Gelwi Strite) | 4.1151 | [Not found] | -- |
| süss-holtz | 26 | Süessholz | No species cited | 2.1259 | Süssholz | Glycyrrhiza glabra L.; Solanum dulcamara L. ? | 2.725, 4.353 | suessholz | Glycyrrhiza glabra L. |
| terpenntin, terpetinum, terprini, terpetin, terpatin, Derpentin* | 88 | terpetin, terpentin | Terpentinharz, resina terebinthina; "wird aus Venedig geführt" | 8.1677 | Terpentin-baum; Terpentin-kiefer | Larix decidua; Pistacia terebinthus L. | 2.1178; 3.792 | terebinthina | Pistacia terebinthus L. |
| Tubenn kröpfflin | 126 | Tubenchropfwasser; Tubenchropf, Tubenchnopf | Arznei aus Taubenkropf-Leimkraut, Silene vulgaris; 1) Scilla bifolia, 2) Musc. botr., 3) Prim. elat., 4) Viola can., 5) Rubus caes., 6) Fumaria off., 7) Blasiges Leimkraut, Sil. infl., 8) Stellaria media, 9) Ajuga chamaepyt. ? | 16.1817; 3.848 | Taubenkropf, Tubenchropf, Taubenkröpflein | Fumaria officinalis L., Muscari spp., Primula veris L., Silene vulgaris L. (S. inflata), Viola canina L. (1); Corydalis cava (L.) Schweigg. & Körte, Phyteuma spicata L., Rubus caesius L., (2); Anthyllis vulneraria L., *(Fortsetzung siehe Kommentar) | 1.1192, 2.508, 1404, 3.228, 720, 1450, 4.162, 317, 507, 1166 | taubenkropf, taubenchropen | Anthyllis vulneraria L., Fumaria officinalis L., Verbena officinalis L. |
| venumgrecum | 167 | [Not found] | -- | -- | [Not found] | -- | -- | foenum graecum | Trigonella foenum-graecum L. |

**SM – Table S2 – Assessment of RBH plant names, category A** (Extract from the RBH database on dermatological recipes / Reference list, see last page)

| Plant name RBH | ID | Plant name Idiotikon | Sci name Idiotikon | Reference Idiotikon | Plant name Marzell | Sci name Marzell | Reference Marzell | Plant name Fischer | Sci name Fischer |
|---|---|---|---|---|---|---|---|---|---|
| verbena | 104 | verbena (unter Eisenkraut/Isechrut) | Verb. off., gemeines Eisenkraut | 3.887 | verbena | Verbena officinalis L. | 4.1045 | verbena | Verbena officinalis L., Lysimachia arvensis (L.) U.Manns & Anderb. |
| viöl | 143 | viole; vyolkrut, veielichrut, violarum | Viola odorata, and other similar plants | 1.633; 3.889 | viole, viönli, viola, veilchen | "Veilchen": 1) Viola odorata L., 2) other Viola spp., 3) Cheiranthus cheiri, Myosotis spp., Vinca minor, etc. | 4.1155 | viola, veyal; herba violaria | Viola odorata L.; Viola tricolor L. var. arvensis und var. maxima |
| viönlj, vigell [-öl] | 97 | violen-saft; viönli, vigeli; viöndliwurtz | Iris florentina rhizome | 7.365; 10.1721; 16.1726 | viönli; zams viönli, waldviönli; violenwurtz, viöliworzle | Erysimum × cheiri (L.) Crantz (Cheiranthus cheiri L.), Matthiola incana (L.) W.T.Aiton; Viola odorata L.; Iris x germanica L. (I. florentina L.) | 1.918; 4.1169; 2.1019, 1025 | uioln | Viola odorata L. |
| vnnserr frouwen mentle | 145 | Unser lieben Frauen mänteli | Gem. Frauenmantel, Alchem. vulg. | 4.342 | Vnser Frawen mantel, frauen mänteli | Alchemilla vulgaris L. | 1.175 | unser frouwen mantel | Alchemilla vulgaris L. |
| voglj krutt | 134 | Vogelchrut | Stellaria media, S. als.; Anagallis arv., A. caer., Veronica arv., Polygonum arv., Plantago major, Capsella bursa past., Senecio vulg., Prismatocarpus speculum, Viscum album, Geranium rob. | 3.899 | Vogelchrut, Vögelichrut | Anagallis arv. (AG), Geranium rob. (VS), Legousia speculum-veneris (1xCH), Plantago major (AG), Polygonum avic. (ZH et al.), Senecio vulgaris (viele), Stellaria media (viele), Veronica arvensis (AG), Vicia cracca (UR), Vicia sepium (SG), Viscum album (AG) | 1.258; 2.662, 1223; 3.828, 901; 4.280, 500, 1056, 1117, 1136, 1204 | vogelkrut | Stellaria media Dill. und andere Arten |
| wal wurtzen | 81 | walwurz | Beinwell | 16.1757 | Wallwurz | Symphytum officinale L. | 4.537 | walwurz; wallwurz | Symphytum officinale L. (1); Anagallis arvensis L. ? (2) |

**SM – Table S2 – Assessment of RBH plant names, category A** (Extract from the RBH database on dermatological recipes / Reference list, see last page)

| Plant name RBH | ID | Plant name Idiotikon | Sci name Idiotikon | Reference Idiotikon | Plant name Marzell | Sci name Marzell | Reference Marzell | Plant name Fischer | Sci name Fischer |
|---|---|---|---|---|---|---|---|---|---|
| wald farn [grosse] | 162 | Waldfarn | Rippenfarn = Blechnum spicant | 1, 1019 | Grosser Waldfarn; Waldfarn | "Grosser": Pteridium aquilinum (L.) Kuhn (hist. Texte, 2x D); Dryopteris filix-mas (L.) Schott (Aspidium filix-mas) (2x hist. Texte), Athyrium filix-femina (L.) Roth. (1x D), Blechnum spicant (L.) Roth. (1x CH), Polypodium vulgare L. (1x) | Diverse | [Not found] | -- |
| waldholunder[2] | n.a. | [Not found] | -- | -- | Waldholunder | Sambucus racemosus L. | 4.134 | [Not found] | -- |
| wald meister | 83 | Waldmeister | Asperula od. | 12.26 | Waldmeister | Galium odoratum (L.) Scop. (Asperula odorata L.) (1); Lonicera periclymenum L. (2); Adoxa moschatellina L. ? (3) | 1. 123, 469; 2.1384 | waltmeister | Galium odoratum (L.) Scop. (Asperula odorata L.) |
| weg grass | 99 | weggras | Knöterich, Polygonum | 2.797 | weggras | Polygonum aviculare L. | 3.893 | weggraz | Polygonum aviculare L. |
| wegrich | 4 | Wĕgrich (Wĕgerich) | Wegerich, 1) Plantago major, P. media; 2) P. lanceolata; 3) P. alpina | 15.952 | Wegerich | 1) Plantago major L., P. media L., 2) P. lanceolata L. | 3.806, 816 | wegerich | Plantago major L., P. media L. |
| wermUtt | 48 | wermuet | Artemisia absinthium (andere Arten kommen eventuell als lokale Substitute in Frage, z.B. A. pontica) | 16.1510 | wermut | Artemisia absinthium L. | 1.421 | wermut | Artemisia abstinthium L. |
| weÿrauch, wÿssen wieruch, -wierauch [see "olibanum"] | 89 | Weisen Wierauch, wiser wirhoch, olibanum (Weihrauch) | Weihrauch | 6.97 | [Not found] | -- | -- | weyrauch | Boswellia spp., B. serrata Roxb. ex Colebr. (B. thurifera) |
| weÿtzenn | 111 | Weizen, Weiss, Weiz, weitzen | Triticum aestivum | 16.1888 | Weizen | Triticum aestivum L. (T. vulgare L.) | 4.826 | weizze | Triticum aestivum L. (T. sativum L.) |

**SM – Table S2 – Assessment of RBH plant names, category A** (Extract from the RBH database on dermatological recipes / Reference list, see last page)

| Plant name RBH | ID | Plant name Idiotikon | Sci name Idiotikon | Reference Idiotikon | Plant name Marzell | Sci name Marzell | Reference Marzell | Plant name Fischer | Sci name Fischer |
|---|---|---|---|---|---|---|---|---|---|
| wilden boleÿ, kleinen costenntz | 79 | Wildpoley | Serpyllus; Mentastrum; Zopyron, Clinopodium dicitur | 4.1181 | Wilde Poley | Calamintha officinalis Moench (syn. C. montana, M. calamintha, S. calamintha) = Clinopodium nepeta (L.) Kuntze*, C. nepeta subsp. glandulosum (Req.) Govaerts [Plant List]*; Mentha arvensis L.; Thymus serpyllum L.** | 1.713; 3.146; 4.709 | Wild polai | Thymus serpyllum L. |
| wisse mülleblümlin | 76 | Müli-blüemli | Anem. hep., Leberblümchen | 5.83 | Mühlen-blume, Müli-blüemli | Hepatica nobilis Schreb. (syn. Anemone hepatica L.) ? (Blüten sind ist meist lila, selten weisslich) | 1.276 | [Not found] | -- |
| wull krutt (mitt denn gellen blUmen), wollen krut | 12 | Wullchrut, Wollenchrut | Wollkraut, Verb. spec., bes. kleinblumige / echte Königskerze, V. thaps., und die grossblumige / gemeine Königskerze, V. thapsiforme | 3.914 | Wollkraut, Wullchrut | Verbascum spp., mainly V. thapsiforme, V. phlomoides, V. thapsus, V. nigrum | 4.1033, 1.339, 3.1006, 4.858 | wullkraut, wullachchrawt | Verbascum thapsus L., V. densiflorum Bertol. (V. thapsiforme Schrad.), V. phlomoides L. |
| wund blümlin | 102 | Wunt-Blüemli | herba sanctae Mariae. Gemeint ist wohl Arnica mont., Marienkraut | 5.91 | Wundblümlein, Wundblüemli | Hypericum perforatum L. | 2.950 | [Not found] | -- |
| wÿssen nacht schatten | 22 | Braunwurz oder nachtschat, weiss und braun | Scrof. nod. | 8.1493 | Weisser Nachtschatten | Scrophularia oblongifolia Loisel. (S. umbrosa Dumort.) | 4.195 | [Not found] | -- |
| ÿbschenn | 108 | Ibische, Ibsche, Ybisch | Althaea officinalis L. (1); Ononis spinosa L. (incl. syn. O. repens L.) (2); Hyssopus officinalis L. (3); Taxus baccata L. (4) | 1.47 | ibisch, ybesch, ibsche | Althaea officinalis L. (1), Hyssopus officinalis L. (2), Ononis spinosa L., Taxus baccata L. (3) | 1.230, 2.967, 3.400, 4.656 | ybisch, ybische, ibische | Althaea officinalis L. |

**SM – Table S2 – Assessment of RBH plant names, category A** (Extract from the RBH database on dermatological recipes / Reference list, see last page)

| Plant name RBH | ID | Plant name Idiotikon | Sci name Idiotikon | Reference Idiotikon | Plant name Marzell | Sci name Marzell | Reference Marzell | Plant name Fischer | Sci name Fischer |
|---|---|---|---|---|---|---|---|---|---|
| zimett | 21 | Zimmet- | No species cited | 12.28 | Zimmetbaum; Wilder Zimmet, Mutterzimmet | C. verum J.Presl (C. ceylanicum Br.); Cinnamomum cassia (L.) J.Presl (C. cassia Bl.) | 2.1005 | zimetrinden | Cinnamomum cassia (L.) J.Presl (C. aromaticum Nees), C. verum J.Presl (C. zeylanicum Nees) |
| zitt lossen | 178 | Zitlose | 1. zur Unzeit blühende Pflanze, colch. aut. 2. zur Unzeit blühende Pflanze, im frühesten Frühjahr blühende Pflanzen a) tuss. farf. b) bell. per. c) gal. niv. d) croc. vern. e) leuc. vern. f) prim. el. g) prim. aur. h) prim. ac. i) an. nem. | 3.1437 | Zeitlose / Kleine Zeitlose | Colchicum autumnale L., Leucojum vernum L., Galanthus nivalis L., Crocus albiflorus Kit. (1); Narcissus spp., Primular veris L. (2); evtl. auch Bellis perennis L. (3) | 1.1071; 2.1259 | zeitloze | Colchicum autumnale L., C. variegatum L. |
| zittwan | 39 | Zitwen-wurtz | Rhizom der weissen Curcuma, Curcuma zedoaria | 16.1760 | Zitwar, zitwan | Curcuma zedoaria Roscoe | 1.1269 | zittvar; zitwan | Curcuma zedoaria (Christm.) Roscoe |

**Dermatological plants and their uses in the *Receptarium* of Burkhard III from Hallwyl (RBH) from 16th century Switzerland – Data mining a historical text and preliminary *in vitro* screening –** Jonas Stehlin, Ina Albert, Thomas Frei, Barbara Frei Haller, Andreas Lardos – February 2024.

**SM – Table S3 – Assessment of RBH plant names, categories B and C** (Extract from the RBH database on dermatological recipes / Reference list, last page)

| Plant name RBH | ID | Plant name Cat. B - Bock | Sci name Cat. B - Bock (Hoppe) | Plant name Cat. B - Fuchs | Sci name Cat. B - Fuchs (Dobat) | Plant name Cat. C - Schneider | Sci name Cat. C - Schneider |
|---|---|---|---|---|---|---|---|
| agrimonienn (klein vnnd gross) | 64 | Agrimonia | Agrimonia eupatoria L. | Agrimonia | Agrimonia eupatoria L. | agrimonia | Agrimonia eupatoria L. |
| alat, alatt | 125 | Alantwurzel | Inula helenium L. | Alantwurz | Inula helenium L. | Alant | Inula helenium L. |
| albunn grecum | 121 | [Not found] | -- | [Not found] | -- | Weisse Hirschwurz? | Lasterpitium latifolium L. (Cervariae albae, Gentianae albae) |
| äpfen distell | 112 | [Not found] | -- | [Not found] | -- | [Not found] | -- |
| Aristologia rottunda | 173 | Aristolochia rotunda | Artistolochia pallida Willd., A. rotunda L. | [Not found] | -- | aristologia rotunda | Artistolochia pallida Willd., A. rotunda L. |
| aronenn | 156 | Aron | Arum maculatum L. | Aron | Arum maculatum L. | Aron | Arum maculatum L. |
| attich | 19 | Attich | Sambucus ebulus L. | Attich | Sambucus ebulus L. | Attich | Sambucus ebulus L. |
| bach bumlen | 144 | Bach bungen | Veronica beccabunga L., Veronica anagallis-aquatica L. (als "ganz kleines Geschlecht") | Bachpungen | Veronica beccabunga L. | Bachbungen | 1) Veronica beccabunga L., 2) Veronica anagallis-aquatica L. |
| barbenn | 63 | [Not found] | -- | [Not found] | -- | Barbenkraut, Barbarakraut | Barbarea vulgaris L. |
| berÿ | 175 | [Not found] | -- | [Not found] | -- | [Not found] | -- |
| blauw Viönnlinn | 133 | Violen, blaw | Viola odorata L. | Blaw violen | Viola odorata L. | Blaue Violen | Viola odorata L. |
| bonnen | 172 | Welsch-bonen / Feld-bonen, Teutschbonen | Phaseolus vulgaris L. / Vicia faba L. | Welsch Bohnen | Phaseolus vulgaris L. / Vicia faba L. | Bohnen | Phaseolus vulgaris L., Vicia faba L. |
| boum öll, baum öll, boumöll | 67 | Ölbaum | Olea europaea L. | [Not found] | -- | Baumöhl, Baumöle | Oleum olivarum aus Olea europaea |
| boumwollen | 42 | [Not found] | -- | Baumwoll | Gossypium herbaceum L. | baumwolle | Gossypium herbaceum L. |
| breitt vnnd spitz wëgrich | 24 | Breiter Wegerich; Spitzer Wegerich | Plantago media L.; Plantago lanceolata L. | Breiter Wegerich; Spitzer Wegerich | Plantago media L.; Plantago lanceolata L. | Breiter Wegerich; Spitzer Wegerich | Plantago media L., P. major L.; Plantago lanceolata L. |

**SM – Table S3 – Assessment of RBH plant names, categories B and C** (Extract from the RBH database on dermatological recipes / Reference list, last page)

| Plant name RBH | ID | Plant name Cat. B - Bock | Sci name Cat. B - Bock (Hoppe) | Plant name Cat. B - Fuchs | Sci name Cat. B - Fuchs (Dobat) | Plant name Cat. C - Schneider | Sci name Cat. C - Schneider |
|---|---|---|---|---|---|---|---|
| breitten wëgrich | 59 | Wegerich, breiten | Plantago media L.; Plantago media L. | Breiter Wegerich | Plantago media L.; Plantago media L. | Breiter Wegerich | Plantago media L., P. major L. |
| Brun bethonienn, Brunne Betonicken | 18 | Bathonienkraut | Stachys officinalis (L.) Trevis. | Bethonien, braune | Betonica officinalis L | braune Bethonie, Betonica | Stachys officinalis (L.) Trevis. |
| brunen kressich, brunenn kressich, brun kressich | 28 | Brunn Cress | Nasturtium officinale R.Br. | Brunnenkress | Nasturtium officinale R.Br. | Brunnn-Cress, Brunnenkresse | Nasturtium officinale R.Br. |
| bUch spick (mitt gälbenn blUmen) | 151 | [Not found] | -- | [Not found] | -- | Grosses Mausöhrchen, fanzösisches Lungenkraut, Kostkraut, Mauer-Habichtskraut | Hieracium murorum L. |
| camillien, cammillien | 8 | Camillien | Matricaria chamomilla L. | Camillen | Matricaria chamomilla L. | chamillenblumen, camomilla | Matricaria chamomilla L. |
| cardobenedickten | 14 | Cardobenedict | Centaurea benedicta L. (Cnicus benedictus L.) | Cardobenedict | Centaurea benedicta L. (Cnicus benedictus L.) | Cardobenedikten, Cardo benedict | Centaurea benedicta L. (Cnicus benedictus L.) |
| deschel krutt | 41 | Teschelkraut | Capsella bursa-pastoris (L.) Medik. | Däschelkraut | Capsella bursa-pastoris (L.) Medik. | deschelkraut, teschelkraut | Capsella bursa-pastoris (L.) Medik. |
| diptannana | 174 | Dictam | Dictamnus albus L., | [Not found] | -- | diptan, diptamnum | Dictamnus albus L., Origanum dictamnus L. |
| dragantz | 170 | [Not found] | -- | [Not found] | -- | dragant | Astragalus gummifer Labill., A. microcephala Willd., Astragalus spp. |
| edle salbinen | 114 | Edel salbei | Salvia officinalis ssp. maior / ssp. minor | [Not found] | Salvia officinalis ssp. maior / ssp. minor | Edel-Salbey | Salvia officinalis L. |
| eerenpriss, erenbriss, erenn briss | 84 | Ehrenbreiss | Veronica officinalis L. (1. Geschlecht); V. serpyllifolia L. (2. Geschlecht) | Ehrenbreis | Veronica officinalis L. | Ehrenbreis | Veronica officinalis L. |
| egelkrutt | 109 | Egelkraut | Lysimachia nummularia L. | Egelkraut | Lysimachia nummularia L. | Egelkraut | Lysimachia nummularia L. |

**SM – Table S3 – Assessment of RBH plant names, categories B and C** (Extract from the RBH database on dermatological recipes / Reference list, last page)

| Plant name RBH | ID | Plant name Cat. B - Bock | Sci name Cat. B - Bock (Hoppe) | Plant name Cat. B - Fuchs | Sci name Cat. B - Fuchs (Dobat) | Plant name Cat. C - Schneider | Sci name Cat. C - Schneider |
|---|---|---|---|---|---|---|---|
| eichen, eichin | 136 | Eychbaum | Quercus robur L. | Eychbaum | Quercus robur L. | Eiche | Quercus robur L., Q. petraea (Matt.) Liebl. und andere mitteleuropäische Eichenarten |
| endiuien | 98 | Endiuia, grösst zam Endiuia; zam Wegwart, zam Cihcorea | Cichorium endivia L. | Endivia | Cichorium endivia L. | Endivie | Cichorium endivia L., Endivienwasser = Succus Endiviae, Intybi |
| epich (safft) | 23 | Epff; Ephew, Eppich | Apium graveolens L.; Hedera helix L. | Epff | Apium graveolens L.; Hedera helix L. | Eppichsaft, Eppichwurtz; Eppich; Eppich | Apium graveolens L. (A. palustre), Wildtyp; Hedera helix L. |
| erdberre | 77 | Erdberen | Fragaria vesca L. | Erdtbeerkraut | Fragaria vesca L. | Erdbeere | Fragaria vesca L. |
| erdrauch | 148 | Erdrauch | Fumaria offcinalis L. | Erdtrauch | Fumaria offcinalis L. | Erdrauch | Fumaria offcinalis L. |
| Eschinne | 101 | Eschernholtz | Fraxinus excelsior | [Not found] | Fraxinus excelsior | Esche | Fraxinus excelsior L. |
| firnnis | 119 | [Not found] | -- | [Not found] | -- | Resina sandaraca, Sandarak | Tetraclinis articulata (Vahl) Mast |
| flachs, lÿnn | 53 | Flachs | Linum usitatissimum L. | Flachs | Linum usitatissimum L. | flachs, lein, linum | Linum usitatissimum L. |
| full boümen | 149 | Fulbaum | Frangula alnus Mill. | [Not found] | -- | Faulbaum | Frangula alnus Mill. |
| fünnff fingerr krutt | 141 | Fünffingerkraut | Potentilla verna L. (das kleinst), P. argentea L. (das zweite), P. reptans L. (das dritt und gemein), P. alba L. (das viert und fremd), P. recta L. | Fünffingerkraut | Potentilla alba L., P. reptans L., P. pusilla Host. (P. neumanniana)? | Fünffingerkraut | Potentilla reptans L., andere Potentilla-Arten |
| für blUmen | 116 | [Not found] | -- | [Not found] | -- | [Not found] | -- |
| Gallöpffell | 49 | Gallöpffel | Quercus robur L. (Q. pedunculata) | Gallöpfel | Quercus robur L. (Q. pedunculata) | Galläpfel | Quercus infectoria G.Olivier und weitere Quercus spp. |

**SM – Table S3 – Assessment of RBH plant names, categories B and C** (Extract from the RBH database on dermatological recipes / Reference list, last page)

| Plant name RBH | ID | Plant name Cat. B - Bock | Sci name Cat. B - Bock (Hoppe) | Plant name Cat. B - Fuchs | Sci name Cat. B - Fuchs (Dobat) | Plant name Cat. C - Schneider | Sci name Cat. C - Schneider |
|---|---|---|---|---|---|---|---|
| ganfer, ganfferr | 95 | [Not found] | -- | [Not found] | -- | kampher | Cinnamomum camphora (L.) J.Presl (1); Dryobalanops aromatica C.F.Gaertn. (D. sumatrensis (J.F.Gmel.) Kosterm.) (2) |
| genist blUmen | 65 | [Not found] | -- | [Not found] | -- | Ginster, Genista | Cytisus scoparius (L.) Link; Spatium junceum L. |
| Gilge, wis gilge, wÿss Gilge | 86 | weiss Gilgen | Lilium candidum L. | Weiss Gilgen | Lilium candidum L. | Weiss Gilgen | Lilium candidum L. |
| glariett, gloria, glorÿ | 93 | [Not found] | -- | [Not found] | -- | Kirschenklar | Kirschengummi, Kirschenharz, Gummi cerasorum von Prunus avium L.. Wird auch von anderen Prunus Arten gewonnen. |
| glost krutt | 61 | [Not found] | Parietaria officinalis L. | Glaskraut | Parietaria officinalis L. | Glaskraut | Parietaria officinalis L., P. judaica L. |
| Gotz gnad, Gotz gnadt | 100 | Gottsgenad | Geranium robertianum L. | Gottes Gnad | Geranium pratense L. | Gottesgnad | Geranium robertianum L., Grattiola officinalis L. |
| grünne betonien | 106 | Bathonienkraut | Stachys officinalis (L.) Trevis. | Bethonien, weisse | Betonica officinalis L | betonien | Stachys officinalis (L.) Trevis. |
| gUtter heinrich | 82 | gut Heinrich | Chenopodium bonus-henricus L. | Guter Heinrich | Chenopodium bonus-henricus L. | gut Heinrich, guot henrich | Chenopodium bonus-henricus L. |
| haber neslen, kleinn weg neslenn, kleiner nesslen | 9 | Nesselen | Urtica urens L. | Habernessel (Urtica minor, Nessel kleine) | Urtica urens L. | Habernessel, kleine Brennessel | Urtica urens L. |
| haber, haberr | 85 | Habern (zamer) | Avena sativa L. | Habern | Avena sativa L. | Haber | Avena sativa L. |
| hannff | 55 | Hanf | Cannabis sativa L. | Hanff | Cannabis sativa L. | hanf | Cannabis sativa L. |
| hartz, wÿss büll hartz, rott büll hartz, gelüterett hartz* | 57 | Hartz, Kyfferbaum | Pinus cembra L. | [Not found] | -- | Harz | Pinus sylvestris L., P. strobus L., Pinus palustris Will., Picea abies (L.) |

**SM – Table S3 – Assessment of RBH plant names, categories B and C** (Extract from the RBH database on dermatological recipes / Reference list, last page)

| Plant name RBH | ID | Plant name Cat. B - Bock | Sci name Cat. B - Bock (Hoppe) | Plant name Cat. B - Fuchs | Sci name Cat. B - Fuchs (Dobat) | Plant name Cat. C - Schneider | Sci name Cat. C - Schneider |
|---|---|---|---|---|---|---|---|
| | | | | | | | H.Karst. (Abies excelsa DC) |
| haslen | 40 | Haselnüss | Corylus avellana L. | Hasel | Corylus avellana L. | Hasel | Corylus avellana L. |
| heidisch wund krutt | 157 | Heidnisch Wundkraut | Anthyllis vulneraria L.; | Heydnisch Wundkraut | Senecia ovatus (G.Gaertn., B.Mey. & Scherb.) Willd. | -- (keine Einträge für "heidnisch"); Wundkraut; gulden W.; w-cruth; w-chrawt | -- / Anthyllis vulneraria L.; Ajuga reptans L.; Prunella vulgaris L.; Silene dioica (L.) Clairv. |
| heidnisch wundkrutt (mitt den zer schnitnen bletterenn)[1] | 72 | [Not found] | -- | [Not found] | -- | [Not found] | -- |
| Heill aller wéltt | 70 | Heil aller Schaden | Veronica officinalis L. | [Not found] | -- | Heyl aller Schäden | Veronica officinalis L. |
| hirtzen zungen | 54 | Hirtzzung | Asplenium scolopendrium L. (Phyllitis scolopendrium (L.) Newman) | [Not found] | -- | hirtzzung, hischzunge | Asplenium scolopendrium L. (Phyllitis scolopendrium (L.) Newman) |
| holder | 13 | Holderbaum | Sambucus nigra L. | holder | Sambucus nigra L. | Holder | Sambucus nigra L. |
| holtz öpffell | 139 | [Not found] | -- | [Not found] | -- | Holzapfel | Malus sylvestris Mill.: Poma sylvestris (äusserlich als Adstringens verwendet) |
| hünner darm | 105 | Hünerdarm | Stellaria media (L.) Vill. | Hünerdarm | Stellaria media (L.) Vill. | Hünerdarm; Rother Hünerdarm; | Stellaria media (L.) Vill.; Anagallis avensis L. |
| hus wurtzen | 80 | Hauswurz | Sempervivum tectorum L., Sedum telephium L., Sedum album L. | Hauswurtz | Sempervivum tectorum L., Sedum telephium L., Sedum album L., Petrosedum pruinatum (Link ex Brot.) Grulich (Sedum reflexum Cutanda) | Hauswurz | Sempervivum tectorum L., Sedum telephium L. |

**SM – Table S3 – Assessment of RBH plant names, categories B and C** (Extract from the RBH database on dermatological recipes / Reference list, last page)

| Plant name RBH | ID | Plant name Cat. B - Bock | Sci name Cat. B - Bock (Hoppe) | Plant name Cat. B - Fuchs | Sci name Cat. B - Fuchs (Dobat) | Plant name Cat. C - Schneider | Sci name Cat. C - Schneider |
|---|---|---|---|---|---|---|---|
| jmber, ÿmberr | 69 | [Not found] | -- | [Not found] | -- | Ingwer, ymber | Zingiber officinale Roscoe |
| juden öpffell | 132 | Judenöpffel | Citrus medica L. | [Not found] | -- | Judenöpfel | Citrus medica L. |
| kärnenn | 110 | Kern | Triticum spelta L, Triticum aestivum L. (T. vulgare L.) | [Not found] | -- | Kern | Triticum spelta L, Triticum aestivum L. (T. vulgare L.) |
| katzen trübel | 62 | Katzentreubel | Sedum acre L. | Katzentreuble | Sedum acre L. | Katzenträublein | Sedum acre L. |
| kernn gertten | 115 | [Not found] | -- | [Not found] | -- | Kerngerten | Ligustrum vulgare L. |
| klein schwerttel wurtz | 176 | Klein blo Schwertel | Iris sibirica L. | [Not found] | -- | Klein blo Schwertel | Iris sibirica L. |
| klein winter grün | 73 | Wynter gruen | Pyrola rotundifolia Fern. | Wintergrün | Pyrola rotundifolia Fern. | wintergrün | Pyrola rotundifolia L., P. secunda L. |
| knaben krutt | 140 | Knabekraut | Orchis mascula L., O. militaris L., O. morio L., O. ustulata L., Anacamptis pyramidalis (L.) Rich.; Sedum telephium L. | Knabenkraut | Orchis mascula L., O. militaris L., O. morio L., O. ustulata L., Anacamptis pyramidalis (L.) Rich. | Knabenkraut | Orchis mascula L., O. militaris L., O. morio L., O. purpurea Huds., O. ustulata L., Orchis spp.; Anacamptis pyramidalis (L.). Rich.; Platanthera bifolia (L.) Rich.; und evtl. weitere Orchideen-Arten |
| knoblach | 153 | Knoblauch | Allium sativum L. | Knoblauch | Allium sativum L. | Knoblauch | Allium sativum L. |
| knoblach krutt | 146 | Wald knoblauch | Alliaria petiolata Cav. et Grande | Knoblochkraut | Alliaria petiolata Cav. et Grande | Knoblauchkraut; Knoblauch-Gamander, Lachen-Knoblauch | Alliaria petiolata (M.Bieb.) Cavara & Grande (A. officinalis Andrz.); Teucrium scordium L. |
| korble krutt, korblin krutt | 37 | Koerffel (garten oder zam Koerffel) | Anthriscus cerefolium (L.) Hoffm. | Körffel | Anthriscus cerefolium (L.) Hoffm. | Koerbelkraut | Anthriscus cerefolium (L.) Hoffm. |
| küngund krutt | 32 | Küngundkraut | Eupatorium cannabinum L. | Künigundkraut | Eupatorium cannabinum L. | Küngundkraut | Eupatorium cannabinum L. |
| kütenenn | 169 | Kütten | Cydonia oblonga Mill. | Kütten | Cydonia oblonga Mill. | chutten, kütten, quitten | Cydonia oblonga Mill. |

**SM – Table S3 – Assessment of RBH plant names, categories B and C** (Extract from the RBH database on dermatological recipes / Reference list, last page)

| Plant name RBH | ID | Plant name Cat. B - Bock | Sci name Cat. B - Bock (Hoppe) | Plant name Cat. B - Fuchs | Sci name Cat. B - Fuchs (Dobat) | Plant name Cat. C - Schneider | Sci name Cat. C - Schneider |
|---|---|---|---|---|---|---|---|
| lennger ie lieber | 16 | Jelenger je lieber | Keine Angaben, im Eintrag unter Solanum nigrum erwähnt. | Lenger ye lieber | [Not specified, cited chapter on Geissblatt] | je länger je lieber | Solanum dulcamara L.; Veronica chamaedrys L.; Lonicera caprifolium L. |
| lilienn (krutt) | 164 | Gilgen | [Not specified] | Gilgen, weiss | Lilium candidum L. | Lilie, Weiss-Lilie, lilium, lilium album | Lilium candidum L.: u.a. Blüten sind offizinell (Flores lilii albidi, meist als lilium bezeichnet), auch Aqua dest. aus frischen Blütenblättern |
| linnden | 168 | Lindenbaum | Tilia platyphyllus Scop. (die zame), T. cordata Mill. (der wild) | Linden | Tilia sp. | Linde | Tilia platyphyllus Scop., T. cordata Mill. |
| Lor-, lorberr | 7 | Loorbeerbaum | Laurus nobilis L. | [Not found] | -- | Lorbeeren | Laurus nobilis L. |
| lüss krutt | 159 | Leusskraut | Daphne mezereum L., Helleborus foetidus L., Pedicularis palustris L., | Leusskraut | Helleborus foetidus L. | Laus-kraut/wurz, Läuse ... | Daphne mezereum L., Delphinium staphisagria L., Helleborus foetidus L., Pedicularis palustris L., |
| macis | 130 | [Not found] | -- | [Not found] | -- | macis | Arillus (Samenmantel) von Myristica fragrans Houtt. |
| mandel | 96 | Mandelbaum | Prunus dulcis (Mill.) D.A.Webb) | [Not found] | -- | Mandel | Prunus amygdalus Batsch (P. dulcis (Mill.) D.A.Webb) |
| mass blümlin, moss blUmen | 66 | Masslieblin | Bellis perennis L. | [Not found] | -- | Masslieben | Bellis perennis L. |
| mastix | 87 | [Not found] | -- | [Not found] | -- | Mastix | Pistacia lentsicus L. |
| meister wurtz | 27 | Meisterwurtz | Peucedanum ostruthium (L.) W.D.J.Koch | Meisterwurtz | Peucedanum ostruthium (L.) W.D.J.Koch | Meisterwurz | Peucedanum ostruthium (L.) W.D.J.Koch |
| melissen | 127 | Melisse | Melissa officinalis L. | Melisse | Melissa officinalis L. | Melisse | Melissa officinalis L. |

**SM – Table S3 – Assessment of RBH plant names, categories B and C** (Extract from the RBH database on dermatological recipes / Reference list, last page)

| Plant name RBH | ID | Plant name Cat. B - Bock | Sci name Cat. B - Bock (Hoppe) | Plant name Cat. B - Fuchs | Sci name Cat. B - Fuchs (Dobat) | Plant name Cat. C - Schneider | Sci name Cat. C - Schneider |
|---|---|---|---|---|---|---|---|
| mer rettich | 150 | Meerrhetich | Aromracia rusticana G.Gaertn. B.Mey et Scherb. (syn. Armoracia lapathifolia Gilib.) | Meerrhettich | Aromracia rusticana G.Gaertn. B.Mey et Scherb. (syn. Armoracia lapathifolia Gilib.) | Mer-rettich, meerrätich | Aromracia rusticana G.Gaertn. B.Mey et Scherb. |
| miess | 46 | Moos | Usnea spp. | [Not found] | -- | miess, musci | Usnea spp., and other moss or lichen species |
| modelgeer | 182 | Modelgeer | Gentiana cruciata L. | [Not found] | -- | Modelgeer | Gentiana cruciata L. |
| müllj stoub | 137 | [Not found] | -- | [Not found] | -- | Mehl, Amylum tritici | Flour of various Triticum spp. |
| muscat nus, muscatt | 94 | [Not found] | -- | [Not found] | -- | Muskatnuss | Myristica fragrans Houtt. |
| nacht schatt, nachtschatten, nach schatten | 60 | Nachtschadt | Solanum nigrum L.; Solanum dulcamara L. | Nachtschatt | Solanum nigrum L. | Nachtschatten; Bittersüsser Nachtschatten | Solanum nigrum L.; Solanum dulcamara L. |
| näptenn krutt | 163 | Nepten; Nept, zam katzenkraut | Nepeta cataria L. | Nepeta | Nepeta cataria L. | nepeta, katzenkraut; katzenkraut | Nepeta cataria L.; Valeriana officinalis L. |
| natter krutt, natterkrutt | 71 | [Not found] | -- | [Not found] | -- | Natterkopf; Nattergras | Echium vulgare L.; Scorzonera hispanica L. |
| nëgellin, näglin | 6 | [Not found] | -- | [Not found] | -- | negely, Näglen, Gewürz-Nägelein | Syzygium aromaticum (L.) Merr. et L.M.Perry |
| nesslen | 43 | Nesselen | Urtica dioica L., U. urens L., Lamium album L., L. galeobdolon (L.) L., L. maculatum L., L. purpureum L., Stachys sylvatica L. | Nesseln | Urtica dioica L., U. urens L. | Nessel, gemein brennend Nessel; Eiter Nessel | Urtica dioica L.; Urtica urens L. |
| niess wurtz | 166 | Schwarze Niesswurz; Weiss Niesswurz | Helleborus foetidus; Veratrum album | Schwarze Niesswurz; Weiss Niesswurz | Helleborus foetidus; Veratrum album | Niesswurz (Schwarze/ Weisse) | Helleborus niger, viridis, foetidus; Veratrum album, nigrum |

**SM – Table S3 – Assessment of RBH plant names, categories B and C** (Extract from the RBH database on dermatological recipes / Reference list, last page)

| Plant name RBH | ID | Plant name Cat. B - Bock | Sci name Cat. B - Bock (Hoppe) | Plant name Cat. B - Fuchs | Sci name Cat. B - Fuchs (Dobat) | Plant name Cat. C - Schneider | Sci name Cat. C - Schneider |
|---|---|---|---|---|---|---|---|
| olibanum, olibanj [see "weÿrauch"] | 120 | [Not found] | -- | [Not found] | -- | olibanum | Boswellia sacra Flueck. |
| papell, babell, bapellenn krutt | 35 | Rosspappel; Gemeine kess pappel | Malva sylvestris L.; M. neglecta Wallr. | Pappeln | Malva sylvestris L.; M. neglecta Wallr. | Pappel, Rosspappel, Käsepappel; Pappenkraut | Malva sylvestris L., M. neglecta Wallr., M. pusilla Sm. (M. rotundifolia); Taraxacum officinale Wiggers. |
| paris körnlin | 129 | [Not found] | -- | [Not found] | -- | Parisskorn, Paradieskörner | Aframomum melegueta K.Schum. |
| petterlj | 135 | Peterlin | Petroselinum crispum (Mill.) Fuss (Petroselinum hortense Hoffm.) | Gemein Peterlin | Petroselinum crispum (Mill.) Fuss (Petroselinum hortense Hoffm.) | Peterlin | Petroselinum crispum (Mill.) Fuss |
| pfäffer krutt | 184 | Pfefferkraut | Lepidium latifolium L. | Pfefferkraut | Lepidium latifolium L. | Pfefferkraut | Lepidium latifolium L., Satureja hortensis L. |
| pflumen | 17 | Pflumen | Prunus domestica L. | Pflaumen | Prunus domestica L. | Pflaumen | Prunus domestica L. |
| poleÿ, boleÿ, hertzen bleich, hertz bleÿ | 5 | Poleien | Mentha pulegium L. | Poley | Mentha pulegium L. | Poley, Hertzpoley | Mentha pulegium L. |
| reb krutt | 10 | [Not found] | -- | [Not found] | -- | [Not found] | -- |
| rörlin krutt | 160 | Pfaffenroerlin | Taraxacum officinale L. | Pfaffenröhrlin | Taraxacum officinale L. | Rhoerlenkraut, Rohrenkraut | Taraxacum officinalie L. |
| rosenn wurtzell | 47 | Rosenwurz | Rhodiola rosea L. (Sedum roseum (L.) Scop.) | Rosenwurtz | Rhodiola rosea L. (Sedum roseum (L.) Scop.) | Rosenwurz | Rhodiola rosea L. (Sedum roseum (L.) Scop.) |
| rosenn, ross | 36 | Zam garten Rosen; Hannrosen Feldrosen, Haberrosen | Rosa x alba L., R. x damascena Herrm., Rosa x centifolia L.; Rosa gallica L. | Rosen, zam-, wild- | Rosa gallica L., Rosa canina L. | Rose, Rosenöl, Rosenwasser | Rosa gallica L., R. x alba L., R. x damascena Herrm., R. x centifolia L. |
| rot bugglenn, rott buglenn, rott bugenn | 11 | Rother Buck | Artemisia vulgaris L. | Rot Bucken | Artemisia vulgaris L. | Rother Buck | Artemisia vulgaris L. |

**SM – Table S3 – Assessment of RBH plant names, categories B and C** (Extract from the RBH database on dermatological recipes / Reference list, last page)

| Plant name RBH | ID | Plant name Cat. B - Bock | Sci name Cat. B - Bock (Hoppe) | Plant name Cat. B - Fuchs | Sci name Cat. B - Fuchs (Dobat) | Plant name Cat. C - Schneider | Sci name Cat. C - Schneider |
|---|---|---|---|---|---|---|---|
| rotten mangolt | 75 | Mangolt, rote rüben | Beta vulgaris var. cicla L. | Rot Mangolt | Beta vulgaris var. cicla L. | rother Mangold | Beta vulgaris L., Beta rubrae |
| rottenn thannen, rott danj | 68 | [Not found] | -- | [Not found] | -- | Rottanne | Picea abies (L.) H.Karst. |
| rutten | 33 | Raut | Ruta graveolens L. | Raut | Ruta graveolens L. | rautten, Raut | Ruta graveolens L. |
| saffrann | 118 | Saffran | Crocus sativus L.; Carthamus tinctorius L. | Saffran; Wilder garten Saffran | Crocus sativus L.; Carthamus tinctorius L. | Safran; Wilder Saffran; Wiesen Saffran | Crocus sativus L.; Carthamus tinctorius L.; Colchicum autumnale L. |
| safft, so vs dem holtz flüst das jm fürr ligt | 147 | [Not found] | -- | [Not found] | -- | [Not found] | -- |
| salbinenn | 103 | Salbei | Salvia officinalis L.; S. pratensis L. | Salveye; Wild Salbey | Salvia officinalis L.; S. pratensis L. | Salbei; Wilder Salbei; Wild Salbey | Salvia officinalis L.; S. pratensis L.; Teucrium scorodonia L. |
| Sanickel | 45 | Sanickel | Sanicula europaea L. | Sanickel | Sanicula europaea L. | sanikel | Sanicula europaea L. |
| sanndelholtz | 38 | [Not found] | -- | [Not found] | -- | Sandelholz, Sandel holtz; Rotes Sandelholz | Santalum album L. (1); Pterocarpus santalinus L.f. (2) |
| Sannt petters krutt | 29 | Gross St. Peterskraut; Klein St. Peterskraut | Parietaria officinalis L. (P. erecta Mert. et Koch); P. judaica L. (P. ramiflora Moench) | Sant Peterskraut | Parietaria officinalis L. | sant peterskrut, S. Peterskraut, Mauerkraut | Parietaria officinalis L. (P. erecta), P. judaica L. (P. ramiflora Moench, P. diffusa) |
| sanntt johanns krutt | 117 | S. Johanskraut | Hypericum perforatum L. | S. Johannskraut | Hypericum perforatum L. | Johanniskraut, Sanct Johannskraut | Hypericum perforatum L. |
| sant barbel krutt | 179 | S. Barbelkraut | Barbarea vulgaris W.T.Aiton (B. vulgaris L.) | Barbarakraut | Barbarea vulgaris L. | Barbarakraut | Barbarea vulgaris L. |
| schellkrutt, schelkrut, schell krutt | 74 | Schellenkraut | Chelidonium majus L. | Schoelkraut | Chelidonium majus L. | Schellkraut, schelkraut | Chelidonium majus L. |
| schlechenn, schlehen dornn | 34 | Schlehen; Haferschlehe | Prunus spinosa L.; Prunus domestica | Schlehen | Prunus spinosa L.; Prunus domestica | Schlehen | Prunus spinosa L.; Prunus domestica subsp. insititia (L.) Bonnier & Layens (P. |

**SM – Table S3 – Assessment of RBH plant names, categories B and C** (Extract from the RBH database on dermatological recipes / Reference list, last page)

| Plant name RBH | ID | Plant name Cat. B - Bock | Sci name Cat. B - Bock (Hoppe) | Plant name Cat. B - Fuchs | Sci name Cat. B - Fuchs (Dobat) | Plant name Cat. C - Schneider | Sci name Cat. C - Schneider |
|---|---|---|---|---|---|---|---|
| | | | subsp. insititia Poir. ? - var. Juliana | | subsp. insititia Poir. ? - var. Juliana | | insititia L.) - meist Haferschlehe |
| schlüssell blUmen | 183 | Schlüsselblümen | Primular veris L., P. elatior L. | Schlüsselblümen, gel-, weiss- | Primular veris L., P. elatior L. | Schlüsselblumen | Primular veris L., P. elatior L. |
| schwalmenn wurtzell | 171 | Schwalbenwurtz | Vincetoxicum hirundinaria Medik. (Vincetoxicum officinale) | Schwalbenwurtz | Vincetoxicum hirundinaria Medik. (Vincetoxicum officinale) | Schwalbenwurz(el) | Vincetoxicum hirundinaria Medik. (Cynanchum vincetoxicum) |
| schwartzen nacht schatten | 25 | Gemein Nachtschadt | Solanum nigrum L. | Schwarzer Nachtschatt | Solanum nigrum L. | Schwarzer Nachtschatten | Solanum nigrum L. |
| scritten [see "stritten"] | 155 | [Not found] | -- | [Not found] | -- | [Not found] | -- |
| seffe[2] | n.a. | [Not found] | -- | [Not found] | -- | [Not found] | -- |
| sennff krutt | 180 | Senffkräuter | Lepidium latifolium L., Barbaraea vulgaris R. Br. | Gartensenff, gel-, weiss-; Wilder Senf | Sinapis alba L., Eruca sativa L.; Sinapis arvensis L. | Senf, Gartensenf, Wilder Senf | Lepidium latifolium L., Sinapis alba L., Sinapis arvensis L. (1), Satureja hortensis L. (2); Barbaraea vulgaris L., Eruca sativa L. |
| serpentia | 107 | Serpentaria | Bistorta officinalis Delarbre (Persicaria bistorta (L.) Samp.) | [Not found] | -- | serpentaria, radix serpentariae, serpentaria maior/minor | 1) Bistorta officinalis Delarbre (Persicaria bistorta (L.) Samp., Polygonum bistorta L.); 2) Dracunculus vulgaris Schott, Arum maculatum L., A. italicum L. |
| seueboum | 152 | Sevenbaum | Juniperus sabina L. | Sevenbaum | Juniperus sabina L. | Sevina, Sabina, Sevenbaum, Sadebaum | Juniperus sabina L. |
| sigilis solomonis | 165 | Sigillum solomonis | Polygonatum multiflorum (L.) All., P. verticillatum (L.) All. | Sigillum solomonis: Weisswurz / Schmale Weisswurz | Polygonatum multiflorum (L.) All., P. verticillatum (L.) All. | Sigillum salomonis, Salomonssiegel | Polygonatum odoratum (Mill.) Druce (P. officinale Moench), P. verticillatum (L.) All., P. multiflorum (L.) All. |

**SM – Table S3 – Assessment of RBH plant names, categories B and C** (Extract from the RBH database on dermatological recipes / Reference list, last page)

| Plant name RBH | ID | Plant name Cat. B - Bock | Sci name Cat. B - Bock (Hoppe) | Plant name Cat. B - Fuchs | Sci name Cat. B - Fuchs (Dobat) | Plant name Cat. C - Schneider | Sci name Cat. C - Schneider |
|---|---|---|---|---|---|---|---|
| sinauw | 44 | Sinnaw | Alchemilla xanthochlora Rothm. (A. vulgaris L.); Alchemilla xanthochlora Rothm. (A. vulgaris L.) | Synaw | Alchemilla xanthochlora Rothm. (A. vulgaris L.); Alchemilla xanthochlora Rothm. (A. vulgaris L.) | sinau | Alchemilla xanthochlora Rothm. (A. vulgaris L.) |
| spickennardj | 131 | Nardus, spica | Lavandula latifolia Medik. | spicanardi | Lavandula latifolia Medik. | spicanardi (indianisch, vera); Spik-Lavendel, Narde; spica celtica, Nardenkraut | Nardostachys jatamansi DC.; Lavandula latifolia Medik.; Valeriana celtica L. |
| spitzwegrich, spitzen wegrich, spitzen wägrich | 58 | Spitzer Wegerich | Plantago lanceolata L. | Spitzer Wegerich | Plantago lanceolata L. | Spitzer-Wegerich | Plantago lanceolata L. |
| stinckende nesslen | 154 | Gross und Schwartz Andorn; Waldt-Nesseln | Stachys silvaticus L. | Prasium foetidum | Ballota nigra L. | 1) Lamii sylvatici foetidi, Urticae inertis magnae foetidissime; 2) Ballota foetida | 1) Stachys sylvatica L.; 2) Ballota nigra L. |
| stritten | 161 | [Not found] | -- | [Not found] | -- | [Not found] | -- |
| süss-holtz | 26 | Süssholtz | Glycyrrhiza glabra L. | Süessholtz | Glycyrrhiza glabra L. | Süssholtz | Glycyrrhiza glabra L. |
| terpenntin, terpetinum, terprini, terpetin, terpatin, Derpentin* | 88 | [Not found] | -- | [Not found] | -- | Terebinthina vera/cypria/veneta; T. veneta/communis; T. communis/vulgaris | Pistacia terebinthus L. ("...bei uns höchst selten, meistens mit venetianischem Terpentin verfälscht"); Larix decidua Mill. (Lärchenterpentin); Pinus sylvestris L., Pinus pinaster Aiton, Picea abies (L.) H.Karst. (Abies excelsa) |
| Tubenn kröpfflin | 126 | Dubenkropff | Silene vulgaris L. | [Not found] | -- | Taubenkropf, Dubenkropff | Fumaria officinalis L., Silene vulgaris L. |

**SM – Table S3 – Assessment of RBH plant names, categories B and C** (Extract from the RBH database on dermatological recipes / Reference list, last page)

| Plant name RBH | ID | Plant name Cat. B - Bock | Sci name Cat. B - Bock (Hoppe) | Plant name Cat. B - Fuchs | Sci name Cat. B - Fuchs (Dobat) | Plant name Cat. C - Schneider | Sci name Cat. C - Schneider |
|---|---|---|---|---|---|---|---|
| venumgrecum | 167 | Foenograecum | Trigonella foenum-graecum L. | [Not found] | -- | foenum graecum | Trigonella foenum-graecum L. |
| verbena | 104 | Verbena | Verbena officinalis L. | Verbene | Verbena officinalis L. | Verbena | Verbena officinalis L. |
| viöl | 143 | Mertzenviolen: die eldste und zame; Weiss-; Wilden- | Viola odorata L.; Viola albiflora Ob.; Viola canina L. | Violen, Mertzen-, blaw- | Viola odorata L. | Veielkraut, Herba Violaria; Veieln (-kraut?) | 1) Viola odorata L.; 2) Viola tricolor L. |
| viönlj, vigell [-öl] | 97 | Violwutz | Iris germanica L.; Viola odorata L., Matthiola incana (L.) R.Br. (weisse, rote und violette Gartenformen); Erysimum cheiri (L.) Crantz, Hesperis matronalis L. | Violwutz; Violen, blau-, Metzen-; Welsch Veiel; Veiel, gel-, braun-, weiss- | Iris germanica L.; Viola odorata L., Matthiola incana (L.) R.Br. (weisse, rote und violette Gartenformen); Erysimum cheiri (L.) Crantz, Hesperis matronalis L. | Veielöle; violenwurtz; weyse fiolen; gelfiolen | Viola odorata L.; Iris germanica L., I. florentina L.; Matthiola incana L., M. annua Sw.; Erysimum cheiri (L.) Crantz (Cheiranthus cheiri L.) |
| vnnserr frouwen mentle | 145 | Frawenmantel | Alchemilla vulgaris L., A. xanthochlora Rothm. | Unser Frawen Mantel | Alchemilla xanthochlora Rothm. (A. vulgaris L.) | Unser Frauen Mantel | Alchemilla vulgaris L. |
| voglj krutt | 134 | Vogelkraut | Stellaria media Vill. | Vogelkraut | Stellaria media Vill. | Vogelkraut, vogelkrut: gelbes Vogelkraut | Anagallis arvensis L., Stellaria media (L.) Vill.; Senecio vulgaris L. |
| wal wurtzen | 81 | Walwurtz | Symphytum officinale L. | Walwurtz | Symphytum officinale L. | walwurtz | Symphytum officinale L., S. bulbosum Schmer. (letztere Art in der CH nur im südlichsten Tessin) |
| wald farn [grosse] | 162 | Waltfarn; Gross Farnkraut | Dryopteris filix-mas (L.) Schott; Pteridium aquilinum (L.) Kuhn | Waldtfarn | Dryopteris filix-mas (L.) Schott; Pteridium aquilinum (L.) Kuhn | Waldfarn; Waldfarn (weiblicher); Wald-Frauenfarn, Wald-Wurmfarn; | Dryopteris filix-mas (L.) Schott; Pteridium aquilinum (L.) Kuhn; Athyrium filix-femina (L.) Roth |

**SM – Table S3 – Assessment of RBH plant names, categories B and C** (Extract from the RBH database on dermatological recipes / Reference list, last page)

| Plant name RBH | ID | Plant name Cat. B - Bock | Sci name Cat. B - Bock (Hoppe) | Plant name Cat. B - Fuchs | Sci name Cat. B - Fuchs (Dobat) | Plant name Cat. C - Schneider | Sci name Cat. C - Schneider |
|---|---|---|---|---|---|---|---|
| waldholunder[2] | n.a. | Waldholunder | Sambucus racemosus L. | [Not found] | -- | Waldholunder | Sambucus racemosus L. |
| wald meister | 83 | Waldmeister | Galium odoratum (L.) Scop. (Asperula odorata L.) | [Not found] | -- | Waldmeister | Galium odoratum (L.) Scop. (Asperula odorata L.) |
| weg grass | 99 | weggrass | Polygonum aviculare L. | Weggras | Polygonum aviculare L. | weggrass | Polygonum aviculare L. |
| wegrich | 4 | Wegerich: breiter, roter, spitzer | Plantago major L., P. media L., P. lanceolata L. | Wegerich: gross/rot; mittelst/breit; spitzig | Plantago major L., P. media L., P. lanceolata L. | wegerich | Plantago major L. (1), P. media L. (2), P. lanceolata L. (2) |
| wermUtt | 48 | Wermutt | Artemisia abstinthium L. | Wermut | Artemisia abstinthium L. | wermut | Artemisia absinthium L. |
| weÿrauch, wÿssen wieruch, -wierauch [see "olibanum"] | 89 | [Not found] | -- | [Not found] | -- | Weisser Weyrauch, Weihrauch | Boswellia sacra Flueck. |
| weÿtzenn | 111 | Weyssen | Triticum aestivum L. (T. vulgare Vill.) | Weytzen | Triticum aestivum L. | Waizen, Weizen, Weyssen | Triticum aestivum L. |
| wilden boleÿ, kleinen costenntz | 79 | Wild Poley, Klein Bachmünz | Mentha arvensis L. | Wilder Poley | Mentha arvensis L. | Wild Poley, Costentz; Wilder Poley | Thymus serpyllus L. (Serpyllum); Mentha arvensis L., Calamintha officinalis Moench |
| wisse mülleblümlin | 76 | [Not found] | -- | [Not found] | -- | [Not found] | -- |
| wull krutt (mitt denn gellen blUmen), wollen krut | 12 | Wullkräuter | Verbascum thapsus L., V. lychnitis L., V. phlomoides L., V. nigrum L. | Wullkraut (weiss-mennle, weiss-weible, wild, schwarz) | Verbascum thapsus L., V. lychnitis L., V. phlomoides L., V. nigrum L. | wulkrut, Wullkraut | Verbascum thapsus L., V. densiflorum Bertol. (V. thapsiforme Schrad.), V. phlomoides L. |
| wund blümlin | 102 | [Not found] | -- | [Not found] | -- | [Not found] | -- |
| wÿssen nacht schatten | 22 | [Not found] | -- | [Not found] | -- | [Not found] | -- |
| ÿbschenn | 108 | Jbischwurz | Althaea officinalis L. | Ybisch | Althaea officinalis L. | ibisch; ispen | Althaea officinalis L.; Hyssopus officinalis L |

**SM – Table S3 – Assessment of RBH plant names, categories B and C** (Extract from the RBH database on dermatological recipes / Reference list, last page)

| Plant name RBH | ID | Plant name Cat. B - Bock | Sci name Cat. B - Bock (Hoppe) | Plant name Cat. B - Fuchs | Sci name Cat. B - Fuchs (Dobat) | Plant name Cat. C - Schneider | Sci name Cat. C - Schneider |
|---|---|---|---|---|---|---|---|
| zimett | 21 | [Not found] | -- | [Not found] | -- | zimmet | Cinnamomum cassia (L.) J.Presl, C. verum J.Presl (C. zeylanicum Nees) |
| zitt lossen | 178 | Zeitlöslin | Colchicum autumnale L. | Zeitlosen | Colchicum autumnale L. | Zeitlosen | Colchicum autumnale L., C. variegatum L. |
| zittwan | 39 | Gelber Zittwan | Zingiber zerumbet (L.) Roscoe ex Sm. | [Not found] | -- | zittwerwurtz; zittwan | zittwerwurtz: Curcuma zedoaria (Christm.) Roscoe; zittwan (pseudozedoaria): Zingiber zerumbet (L.) Roscoe ex Sm. |

**Dermatological plants and their uses in the *Receptarium* of Burkhard III from Hallwyl (RBH) from 16th century Switzerland – Data mining a historical text and preliminary *in vitro* screening –** Jonas Stehlin, Ina Albert, Thomas Frei, Barbara Frei Haller, Andreas Lardos – February 2024.

**SM – Table S4 – Medicinal plant use reports** (Extract from the RBH database on dermatological recipes)

| txtPlantName_ABvH | txtDrugName_ABvH | txtPlantpart | txtCitedUse | txtCombinedUse | txtRecSig | txtRecTyp |
|---|---|---|---|---|---|---|
| agrimonienn (klein vnnd gross) | agrimonienn (klein vnnd gross) | Not mentioned | zU villen presten | presten, zu villen bresten [äusserlich behandelt] | 103b,98r.04.z01 | Simple herbal |
| agrimonienn (klein vnnd gross) | agrimonienn (klein vnnd gross) | Not mentioned | Wunnd salb | wunden | 105a,99v.03 | Composed mixed |
| agrimonienn (klein vnnd gross) | agrimonienn (klein vnnd gross) safft | Preparation | zU villen presten | presten, zu villen bresten [äusserlich behandelt] | 103b,98r.04 | Composed mixed |
| alat, alatt | alat wurtz, alatt | Root | einn angesicht als obs vss setzig sige | vs satz, vssetzigkeitt | 128b,123r.01 | Composed mixed |
| albunn grecum | albunn grecum | Not mentioned | zU altenn scheden welchs etzt vnnd züch zU eitter reinigt | zU alltenn schadenn | 114b,109r.02.v01 | Composed mixed |
| äpfen distell | äpfen distell | Not mentioned | gUtt blatterenn | Blattern | 107a,101v.08.v02 | Composed mixed |
| äpfen distell | äpfen distell | Not mentioned | eÿssenn | eÿssenn | 107a,101v.08.v01 | Composed mixed |
| Aristologia rottunda | Aristologia rottunda | Not mentioned | Dornn vnnd pfill vss zU ziechenn | Dornn vnnd pfill vss zU ziechenn | 156a,150v.01 | Composed herbal |
| aronen | aronen safft | Leaf | wurm am finngerr | wurm am finnger | 131b,126r.02 | Composed mixed |
| attich | attich wurtz | Root | Für nüw erhabne geschwulst | geschwulst | 051b,45r.04 | Composed mixed |
| bach bumlen | bach bumlen | Not mentioned | Man sol wüssenn, das drierlein wertzen sinnd, das manns nitt mitt einerlei vertribt, ... die annderen vertribt man...: wen ein mensch da gschwollen das er nitt kan zU stUll gang | figwertzenn, fÿg wertzenn, fÿg Blatterenn | 121b,116r.03.z01 | Composed mixed |
| barbenn safft | barbenn safft | Preparation | zU villen presten | presten, zu villen bresten [äusserlich behandelt] | 103b,98r.04 | Composed mixed |
| berÿ | berÿ | Fruit | Dornn vnnd pfill vss zU ziechenn | Dornn vnnd pfill vss zU ziechenn | 156a,150v.02 | Composed herbal |
| blauw viönlin, blauw viönnlinn | blauw viönlin wasserr | Preparation | fig wertzenn | figwertzenn, fÿg wertzenn, fÿg Blatterenn | 122a,116v.01 | Simple herbal |
| blauw viönlin, blauw viönnlinn | blauw Viönnlinn | Not mentioned | figwertzenn | figwertzenn, fÿg wertzenn, fÿg Blatterenn | 120b,115r.01 | Composed herbal |
| bonnen | bonnen blUst | Flower | hitziger schadenn | Branndt | 136b,131r.04 | Simple herbal |
| boum öll, baum öll, boumöll | boum öll, baum öll, boumöll | Fatty oil | heilett brannd vnnd erfrores | Branndt | 104a,98v.02 | Composed mixed |
| boum öll, baum öll, boumöll | boum öll, baum öll, boumöll | Fatty oil | figwertzenn die zU vertribenn | figwertzenn, fÿg wertzenn, fÿg Blatterenn | 120b,115r.03.v01 | Composed mixed |
| boum öll, baum öll, boumöll | boum öll, baum öll, boumöll | Fatty oil | Uber alle geschwärr | geschwërr, geschwärr, geschwerr | 106a,100v.02 | Composed mixed |
| boum öll, baum öll, boumöll | boum öll, baum öll, boumöll | Fatty oil | das eim das glid Wasser Verstadt | glid Wasser Verstadt | 161b,156r.01 | Composed mixed |
| boum öll, baum öll, boumöll | boum öll, baum öll, boumöll | Fatty oil | zU villen presten | presten, zu villen bresten [äusserlich behandelt] | 103b,98r.04 | Composed mixed |
| boum öll, baum öll, boumöll | boum öll, baum öll, boumöll | Fatty oil | Für denn bössenn magerr | rud vnnd magerr | 127a,121v.03 | Composed mixed |
| boum öll, baum öll, boumöll | boum öll, baum öll, boumöll | Fatty oil | Schwartz blatteren | Schwartz blatteren | 107a,101v.01 | Composed mixed |
| boum öll, baum öll, boumöll | boum öll, baum öll, boumöll | Fatty oil | so die kinndt bösse füslin hannd | so die kinndt bösse füslin hannd | 127b,122r.07 | Composed mixed |
| boum öll, baum öll, boumöll | boum öll, baum öll, boumöll | Fatty oil | wen ein mensch das harnn hatt | wen ein mensch das harnn hatt | 120b,115r.03.v02 | Composed mixed |

**SM – Table S4 – Medicinal plant use reports** (Extract from the RBH database on dermatological recipes)

| txtPlantName_ABvH | txtDrugName_ABvH | txtPlantpart | txtCitedUse | txtCombinedUse | txtRecSig | txtRecTyp |
|---|---|---|---|---|---|---|
| boum öll, baum öll, boumöll | boum öll, baum öll, boumöll | Fatty oil | zU frischen wunndenn | wunden | 106a,100v.01 | Composed mixed |
| boum öll, baum öll, boumöll | boum öll, baum öll, boumöll | Fatty oil | zU alltenn schadenn | zU alltenn schadenn | 105a,99v.05 | Composed mixed |
| boumwollen | boumwollen zäpfflin | Reparation | so aber das blUtt selber kompt von der nasen | Nasën blUtten | 079a,72v.06.v01 | Composed herbal |
| breitt vnnd spitz wëgrich | breitt vnnd spitz wëgrich | Not mentioned | Für nüw erhabne geschwulst | geschwulst | 051b,45r.06 | Composed herbal |
| breitten wägrich | breitten wëgrich safft | Preparation | zU villen presten | presten, zu villen bresten [äusserlich behandelt] | 103b,98r.04 | Composed mixed |
| breitten wägrich | breitten wägrich | Not mentioned | für denn wurm | wurm am finnger | 131b,126r.08 | Composed herbal |
| Brun bethonienn, brunne betonicken | Brun bethonienn saft | Preparation | Für nüw erhabne geschwulst | geschwulst | 051b,45r.04 | Composed mixed |
| Brun bethonienn, brunne betonicken | Brunne Betonicken | Aerial part | zU allerlei Wunnden vnnd annderem | wunden | 106b,101r.02 | Composed mixed |
| brunen kressich | Brunenn kressich, die blUmen vnnd die bletter | Flowering herb | für alle gschwulst gUtt | geschwulst | 052a,45v.03 | Simple herbal |
| brunen kressich | brunen kressich | Aerial part | Man sol wüssenn, das drierlein wertzen sinnd, das manns nitt mitt einerlei vertribt, ... die annderen vertribt man...: wen ein mensch da gschwollen das er nitt kan zU stUll gang | figwertzenn, fÿg wertzenn, fÿg Blatterenn | 121b,116r.03.z01 | Composed mixed |
| brunen kressich | brunen kressich | Aerial part | gschwärr | geschwërr, geschwärr, geschwerr | 107a,101v.05 | Simple herbal |
| bUch spick (mitt gälbenn blUmen) | bUch spick (mitt gälbenn blUmen) | Not mentioned | wurm am finngerr | wurm am finnger | 131a,125v.09 | Simple herbal |
| camillien | camillien blUmmen | Flower | Für die geschwulst | geschwulst | 051b,45r.02 | Composed herbal |
| camillien | cammillien oll | Preparation | zU offnenn schaden Vnnd flüssenn | zU offnenn schaden Vnnd flüssenn | 105b,100r.03 | Composed mixed |
| cardobenedickten | cardobenedickten wasser | Preparation | Für die geschwulst | geschwulst | 051b,45r.02.z01 | Composed mixed |
| deschel krutt | deschel krutt safft | Aerial part | so aber das blUtt selber kompt von der nasen | Nasën blUtten | 079a,72v.06.v01 | Composed herbal |
| diptannana | diptannana | Not mentioned | Dornn vnnd pfill vss zU ziechenn | Dornn vnnd pfill vss zU ziechenn | 156a,150v.01 | Composed herbal |
| dragantz | dragantz | Gum, resin | grosse hitz zU eim schadenn schlecht die der hitz hefftig werdt | Branndt | 136b,131r.01 | Composed mixed |
| edle salbinen | edle salbinen bletter | Leaf | für fläcken jnn angesicht | fleckenn | 117b,112r.02.v05 | Composed mixed |
| edle salbinen | edle salbinen bletter | Leaf | verthribt alles geschwerr | geschwërr, geschwärr, geschwerr | 117b,112r.02.v12 | Composed mixed |
| edle salbinen | edle salbinen bletter | Leaf | büllen, vnnd geschwulst | geschwulst | 117b,112r.02.v15 | Composed mixed |
| edle salbinen | edle salbinen bletter | Leaf | Uber allen prestenn gelegt | presten, zu villen bresten [äusserlich behandelt] | 117b,112r.02.v02 | Composed mixed |
| edle salbinen | edle salbinen bletter | Leaf | es thribt den vssatzt | vs satz, vssetzigkeitt | 117b,112r.02.v17 | Composed mixed |
| edle salbinen | edle salbinen bletter | Leaf | zU fullen wunden | zU fullen wunden, bewertt vor fullem fleisch | 117b,112r.02.v03 | Composed mixed |
| edle salbinen | edle salbinen | Not mentioned | figwertzenn die zU vertribenn | figwertzenn, fÿg wertzenn, fÿg Blatterenn | 120b,115r.03.v01 | Composed mixed |
| edle salbinen | edle salbinen | Not mentioned | wen ein mensch das harnn hatt | wen ein mensch das harnn hatt | 120b,115r.03.v02 | Composed mixed |
| edle salbinen | edle salbinen | Not mentioned | wunnden | wunden | 107b,102r.01 | Composed herbal |
| eerenpriss, erenbriss, erenn briss | eerenpriss wasser, erenbriss waserr | Preparation | bössenn schaden | bössenn schaden | 118a,112v.02.v08 | Simple herbal |
| eerenpriss, erenbriss, erenn briss | eerenpriss wasser, erenbriss waserr | Preparation | alle geschwärr heilett vnnd weicht es | geschwërr, geschwärr, geschwerr | 118a,112v.02.v18 | Simple herbal |

**SM – Table S4 – Medicinal plant use reports** (Extract from the RBH database on dermatological recipes)

| txtPlantName_ABvH | txtDrugName_ABvH | txtPlantpart | txtCitedUse | txtCombinedUse | txtRecSig | txtRecTyp |
|---|---|---|---|---|---|---|
| eerenpriss, erenbriss, erenn briss | eerenpriss wasser, erenbriss waserr | Preparation | verthribt all ... gifftig stich der spinen oder würmenn | gifftig stich der spinen oder würmenn | 134a,128v.08.v03 | Composed mixed |
| eerenpriss, erenbriss, erenn briss | eerenpriss wasser, erenbriss waserr | Preparation | rud | rud | 134a,128v.08.v02 | Composed mixed |
| eerenpriss, erenbriss, erenn briss | eerenpriss wasser, erenbriss waserr | Preparation | zitter möller vnnd fläckenn | zitter mall (mossen vnnd flecken) | 134a,128v.08.v01 | Composed mixed |
| eerenpriss, erenbriss, erenn briss | eerenpriss wasser, erenbriss waserr | Preparation | zU allen wunden | zU fullen wunden, bewertt vor fullem fleisch | 118a,112v.02.v07 | Simple herbal |
| eerenpriss, erenbriss, erenn briss | eerenpriss wasser, erenbriss waserr | Preparation | zU offnenn schaden Vnnd flüssenn | zU offnenn schaden Vnnd flüssenn | 105b,100r.03 | Composed mixed |
| eerenpriss, erenbriss, erenn briss | eerenpriss, erenbriss, erenn briss | Not mentioned | das eim das glid Wasser Verstadt | glid Wasser Verstadt | 161b,156r.02 | Composed mixed |
| eerenpriss, erenbriss, erenn briss | eerenpriss, erenbriss, erenn briss | Not mentioned | wund salben | wunden | 104b,99r.01 | Composed mixed |
| egelkrutt | egelkrutt | Aerial part | heillett wunder-lichen, auch fistalam | fistel, fistalam | 106b,101r.06.v02 | Composed herbal |
| egelkrutt | egelkrutt | Aerial part | Altt schadenn, heillett wunderlichen | zU alltenn schadenn | 106b,101r.06.v01 | Composed herbal |
| eichen, eichin | eichin wurm mëll | Other | wurm | wurm [allgemein] | 132a,126v.02.v01 | Composed mixed |
| eichen, eichin | eich öpffell | Gall apple | zitter mall | zitter mall (mossen vnnd flecken) | 134a,128v.06 | Composed mixed |
| eichen, eichin | eichen loub, eichin laub | Leaf | hitziger schadenn | Branndt | 136b,131r.05 | Simple herbal |
| eichen, eichin | eichen loub, eichin laub | Leaf | fÿg wertzenn | figwertzenn, fÿg wertzenn, fÿg Blatterenn | 120b,115r.05.v03 | Composed mixed |
| endiuien | endiuienwasserr | Preparation | zU offnenn schaden Vnnd flüssenn | zU offnenn schaden Vnnd flüssenn | 105b,100r.03 | Composed mixed |
| epich | epich safft | Preparation | Für nüw erhabne geschwulst, Einn annders | geschwulst | 051b,45r.05 | Composed mixed |
| erdberre | erdberre krutt | Aerial part | wund salben | wunden | 104b,99r.01 | Composed mixed |
| erdrauch | erdrauch wasserr | Preparation | für die rud Vnnd denn magerr | rud vnnd magerr | 127a,121v.01 | Composed mixed |
| eschinne, weid eschenn | weid eschenn | Not mentioned | fÿg blatterenn | figwertzenn, fÿg wertzenn, fÿg Blatterenn | 121b,116r.01 | Composed mixed |
| eschinne, weid eschenn | Eschinne rinden | Bark | Wunnden zU samen ziechen | wunden | 106a,100v.03 | Simple herbal |
| firnnis | firnnis | Gum, resin | zU alltenn schaden, es reiniget zücht fleisch trücknett vnnd rimpfft zU samen | zU alltenn schadenn | 114b,109r.01 | Composed mixed |
| flachs, lÿnn | lÿnn öll | Fatty oil | bruch die zum schaden, fürr den Branndt | Branndt | 104a,98v.03 | Composed mixed |
| flachs, lÿnn | flachs samen, flachs samenn | Seed | geschwërr | geschwërr, geschwärr, geschwerr | 107a,101v.07 | Composed mixed |
| flachs, lÿnn | flachs samen, flachs samenn | Seed | Für denn Stich | stich | 094b,89r.05 | Composed herbal |
| full boümen | full boümen holtz, die mitlest rinnden | Bark | Für die rud | rud vnnd magerr | 127a,121v.05 | Composed mixed |
| fünff finngerr krutt, fünnff fingerr krutt | fünnff fingerr krutt | Not mentioned | fÿg Blatterenn | figwertzenn, fÿg wertzenn, fÿg Blatterenn | 121a,115v.05 | Simple herbal |
| fünff finngerr krutt, fünnff fingerr krutt | fünff finngerr krutt, wurtzenn von | Root | fÿg Blatterenn | figwertzenn, fÿg wertzenn, fÿg Blatterenn | 121a,115v.06 | Simple herbal |
| für blUmen | für blUmen wasserr | Preparation | wunden welche full fleisch erzügt, vertribt auch die hitz | zU fullen wunden, bewertt vor fullem fleisch | 107b,102r.02.v01 | Composed herbal |
| für blUmen | für blUmen | Not mentioned | figwertzenn | figwertzenn, fÿg wertzenn, fÿg Blatterenn | 120b,115r.01 | Composed herbal |

**SM – Table S4 – Medicinal plant use reports** (Extract from the RBH database on dermatological recipes)

| txtPlantName_ABvH | txtDrugName_ABvH | txtPlantpart | txtCitedUse | txtCombinedUse | txtRecSig | txtRecTyp |
|---|---|---|---|---|---|---|
| gallöpffell | Gallöpffell | Gall apple | blUtt stellenn | blUtt stellunng | 079b,73r.05 | Simple herbal |
| ganfer, ganfferr | ganfer, ganfferr | Gum, resin | grosse hitz zU eim schadenn schlecht die der hitz hefftig werdt | Branndt | 136b,131r.01 | Composed mixed |
| ganfer, ganfferr | ganfer, ganfferr | Gum, resin | für alerleÿ flüs im anngesicht | flüs jm angsicht | 105b,100r.01 | Composed mixed |
| ganfer, ganfferr | ganfer, ganfferr | Gum, resin | einn angesicht als obs vss setzig sige | vs satz, vssetzigkeitt | 128b,123r.01 | Composed mixed |
| genist blUmen | genist blUmen safft | Preparation | zU villen presten | presten, zu villen bresten [äusserlich behandelt] | 103b,98r.04 | Composed mixed |
| gilgen, wis gilgen, wÿss gilgen | wÿss Giligenn bletterr (wenn sÿ grünn sinndt) | Leaf | Sannt anthonis fürr | Sannt anthonis fürr | 136a,130v.02.v01 | Simple herbal |
| gilgen, wis gilgen, wÿss gilgen | wÿss Giligenn bletterr (wenn sÿ grünn sinndt) | Leaf | schwartzen blatteren | Schwartz blatteren | 136a,130v.02.v02 | Simple herbal |
| gilgen, wis gilgen, wÿss gilgen | wÿss Giligenn bletterr (wenn sÿ grünn sinndt) | Leaf | heillet schlanngen vnnd scorpionn biss | Würm bÿss, schlanngen biss, nater biss | 136a,130v.02.v03 | Simple herbal |
| gilgen, wis gilgen, wÿss gilgen | wiss Gilgen wasserr | Preparation | wunden welche full fleisch erzügt, vertribt auch die hitz | zU fullen wunden, bewertt vor fullem fleisch | 107b,102r.02.v01 | Composed herbal |
| gilgen, wis gilgen, wÿss gilgen | Gilgen öll, wis gilgen öll, wÿss Gilgen öll | Preparation | zU offnenn schaden Vnnd flüssenn | zU offnenn schaden Vnnd flüssenn | 105b,100r.03 | Composed mixed |
| gilgen, wis gilgen, wÿss gilgen | wiss Gilgenn wurtzen, wÿss Giligenn wurtzen | Root | Branndt Löschenn | Branndt | 136a,130v.01 | Composed herbal |
| gilgen, wis gilgen, wÿss gilgen | wiss Gilgenn wurtzen, wÿss Giligenn wurtzen | Root | wurm am finnger | wurm am finnger | 131a,125v.01 | Composed herbal |
| glariett, gloria, glorÿ | glariett, gloria, glorÿ | Gum, resin | Blattern | Blattern | 105a,99v.02 | Composed mixed |
| glariett, gloria, glorÿ | glariett, gloria, glorÿ | Gum, resin | das eim das glid Wasser Verstadt | glid Wasser Verstadt | 161b,156r.06 | Composed mixed |
| glariett, gloria, glorÿ | glariett, gloria, glorÿ | Gum, resin | Für die rud | rud | 127b,122r.06 | Composed mixed |
| glariett, gloria, glorÿ | glariett, gloria, glorÿ | Gum, resin | zU offnenn schaden Vnnd flüssenn | zU offnenn schaden Vnnd flüssenn | 105b,100r.03 | Composed mixed |
| glost krutt | glost krutt safft | Preparation | zU villen presten | presten, zu villen bresten [äusserlich behandelt] | 103b,98r.04 | Composed mixed |
| gotz gnad, gotz gnadt | Gotz gnad wasserr | Preparation | zU offnenn schaden Vnnd flüssenn | zU offnenn schaden Vnnd flüssenn | 105b,100r.03 | Composed mixed |
| gotz gnad, gotz gnadt | gotz gnad wasserr gebrennt | Preparation | fig wertzen | figwertzenn, fÿg wertzenn, fÿg Blatterenn | 121b,116r.06 | Simple herbal |
| gotz gnad, gotz gnadt | gotz gnad wasserr gebrennt | Preparation | das eim das glid Wasser Verstadt | glid Wasser Verstadt | 161b,156r.03.v02 | Composed herbal |
| gotz gnad, gotz gnadt | gotz gnad, Gotz gnadt krutt | Aerial part | fig wertzen | figwertzenn, fÿg wertzenn, fÿg Blatterenn | 121b,116r.05 | Simple herbal |
| gotz gnad, gotz gnadt | gotz gnad, Gotz gnadt krutt | Aerial part | das eim das glid Wasser | glid Wasser Verstadt | 161b,156r.03.v01 | Composed mixed |
| grünne betonien | grünne betonien | Aerial part | haupt wunden, ist nitt also heillsam. sonnder hefftett die selben zU samen jnn kurtzerr zitt, vnnd zücht auch vs die sprisenn der hirnn schallen | haupt wunden, Houbt wunnden | 106b,101r.03 | Simple herbal |
| gUtter heinrich | gUtter heinrich | Not mentioned | wund salben | wunden | 104b,99r.01 | Composed mixed |
| haber neslen, kleinn weg neslenn, kleiner nesslen | haber neslen, oder kleinn weg neslenn | Not mentioned | geschwulsten vnnd wie die zU vertribenn, Erstlich ann eim schenckel | geschwulst | 051b,45r.01 | Simple herbal |
| haber neslen, kleinn weg neslenn, kleiner nesslen | kleiner nesslen wurtzen | Root | gestellt das blUtt gar wunder barlichen | blUtt stellunng | 079a,72v.09 | Simple herbal |
| haber, haberr | haber, haberr | Seed | Einn annders für denn magerr | mager, magerr | 127b,122r.03 | Simple herbal |

**SM – Table S4 – Medicinal plant use reports** (Extract from the RBH database on dermatological recipes)

| txtPlantName_ABvH | txtDrugName_ABvH | txtPlantpart | txtCitedUse | txtCombinedUse | txtRecSig | txtRecTyp |
|---|---|---|---|---|---|---|
| hannff | hannff samenn öll | Fatty oil | bruch die zum schaden, fürr den Branndt | Branndt | 104a,98v.03 | Composed mixed |
| hartz, wÿss büll hartz, rott büll hartz, gelüterett hartz* | hartz, wÿss büll hartz, rott büll hartz | Gum, resin | so hast ein gUtte brannd salben | Branndt | 105b,100r.02 | Composed mixed |
| hartz, wÿss büll hartz, rott büll hartz, gelüterett hartz* | hartz, wÿss büll hartz, rott büll hartz | Gum, resin | Erbgrinndt | erbgrinndt | 160a,154v.01 | Composed herbal |
| hartz, wÿss büll hartz, rott büll hartz, gelüterett hartz* | hartz, wÿss büll hartz, rott büll hartz | Gum, resin | für denn magerr | mager, magerr | 127b,122r.01 | Composed mixed |
| hartz, wÿss büll hartz, rott büll hartz, gelüterett hartz* | hartz, wÿss büll hartz, rott büll hartz | Gum, resin | zU villen presten | presten, zu villen bresten [äusserlich behandelt] | 103b,98r.04 | Composed mixed |
| hartz, wÿss büll hartz, rott büll hartz, gelüterett hartz* | hartz, wÿss büll hartz, rott büll hartz | Gum, resin | Für denn magerr | rud vnnd magerr | 127a,121v.02 | Composed mixed |
| hartz, wÿss büll hartz, rott büll hartz, gelüterett hartz* | hartz, wÿss büll hartz, rott büll hartz | Gum, resin | Wunnd salb | wunden | 105a,99v.03 | Composed mixed |
| hartz, wÿss büll hartz, rott büll hartz, gelüterett hartz* | büll hartz (wÿss / rott) | Gum, resin | zU allerlei Wunnden vnnd annderem | wunden | 106b,101r.02 | Composed mixed |
| hartz, wÿss büll hartz, rott büll hartz, gelüterett hartz* | hartz, wÿss büll hartz, rott büll hartz | Gum, resin | zU altenn scheden welchs etzt vnnd züch zU eitter reinigt | zU alltenn schadenn | 114b,109r.02.v01 | Composed mixed |
| haslen, hassen | haslen balg [hassen balg] | Fruit peel | thUn das vnder ein anderen thU das buluer in die wunden es stadt für warr [blUtt ze stellen] | blUtt stellunng | 079a,72v.05 | Composed herbal |
| heidnisch wundkrutt - mitt den zer schnitnen bletterenn | heidnisch wundkrutt - mitt den zer schnitnen bletterenn | Not mentioned | blUtt ze stellen | blUtt stellunng | 079a,72v.08 | Composed herbal |
| heidnisch wundkrutt - mitt den zer schnitnen bletterenn | heidnisch wundkrutt - mitt den zer schnitnen bletterenn | Not mentioned | wund salben | wunden | 104b,99r.01 | Composed mixed |
| heidnisch wundkrutt, heidisch wund krutt | Heidisch wund krutt | Not mentioned | für denn wurm | wurm am finnger | 131b,126r.06 | Simple herbal |
| heidnisch wundkrutt, heidisch wund krutt | heidnisch wundkrutt öll | Preparation | zU offnenn schaden Vnnd flüssenn | zU offnenn schaden Vnnd flüssenn | 105b,100r.03 | Composed mixed |
| Heill aller wêltt | Heill aller wêltt | Not mentioned | wund salben | wunden | 104b,99r.01 | Composed mixed |
| hirtzen zungen | hirtzen zungen bletter | Leaf | Für denn Stich | stich | 094b,89r.05 | Composed herbal |
| holder | holder bletter, holder | Leaf | gUtt blatterenn | Blattern | 107a,101v.08.v02 | Composed mixed |
| holder | holder bletter, holder | Leaf | eÿssenn | eÿssenn | 107a,101v.08.v01 | Composed mixed |
| holder | holder bletter, holder | Leaf | Für die geschwulst | geschwulst | 051b,45r.02 | Composed herbal |

**SM – Table S4 – Medicinal plant use reports** (Extract from the RBH database on dermatological recipes)

| txtPlantName_ABvH | txtDrugName_ABvH | txtPlantpart | txtCitedUse | txtCombinedUse | txtRecSig | txtRecTyp |
|---|---|---|---|---|---|---|
| holder | holder bletter, holder | Leaf | Altt schadenn, heillett wunderlichen | zU alltenn schadenn | 106b,101r.06.v01 | Composed herbal |
| holder | Holder schützlig, holder schösslÿ | Tip of shoot | wunnd salben | wunden | 106a,100v.05 | Composed mixed |
| holder | holder lattwergen | Preparation | Für die geschwulst | geschwulst | 051b,45r.02.z01 | Composed mixed |
| holtz öpffell | holtz öpffell safft | Preparation | Für fÿgwertzenn | figwertzenn, fÿg wertzenn, fÿg Blatterenn | 121a,115v.03 | Composed mixed |
| hünner darm | hünner darm | Aerial part | zU allerlei Wunnden vnnd annderem | wunden | 106b,101r.02 | Composed mixed |
| hus wurtzen | hus wurtzen safft | Preparation | grosse hitz zU eim schadenn schlecht die der hitz hefftig werdt | Branndt | 136b,131r.01 | Composed mixed |
| hus wurtzen | hus wurtzen | Not mentioned | wund salben | wunden | 104b,99r.01 | Composed mixed |
| jmber, ÿmberr | jmber, ÿmberr | Root | für fläcken jnn angesicht | fleckenn | 117b,112r.02.v05 | Composed mixed |
| jmber, ÿmberr | jmber, ÿmberr | Root | für alerleÿ flüs im anngesicht | flüs jm angsicht | 105b,100r.01 | Composed mixed |
| jmber, ÿmberr | jmber, ÿmberr | Root | verthribt alles geschwerr | geschwërr, geschwärr, geschwerr | 117b,112r.02.v12 | Composed mixed |
| jmber, ÿmberr | jmber, ÿmberr | Root | büllen, vnnd geschwulst | geschwulst | 117b,112r.02.v15 | Composed mixed |
| jmber, ÿmberr | jmber, ÿmberr | Root | Uber allen prestenn gelegt | presten, zu villen bresten [äusserlich behandelt] | 117b,112r.02.v02 | Composed mixed |
| jmber, ÿmberr | jmber, ÿmberr | Root | es thribt den vssatzt | vs satz, vssetzigkeitt | 117b,112r.02.v17 | Composed mixed |
| jmber, ÿmberr | jmber, ÿmberr | Root | für denn wurm | wurm [allgemein] | 131b,126r.11 | Composed mixed |
| jmber, ÿmberr | jmber, ÿmberr | Root | zU fullen wunden | zU fullen wunden, bewertt vor fullem fleisch | 117b,112r.02.v03 | Composed mixed |
| juden öpffell | juden öpffell | Fruit | für fläcken jnn angesicht | fleckenn | 117b,112r.02.v05 | Composed mixed |
| juden öpffell | juden öpffell | Fruit | verthribt alles geschwerr | geschwërr, geschwärr, geschwerr | 117b,112r.02.v12 | Composed mixed |
| juden öpffell | juden öpffell | Fruit | büllen, vnnd geschwulst | geschwulst | 117b,112r.02.v15 | Composed mixed |
| juden öpffell | juden öpffell | Fruit | Uber allen prestenn gelegt | presten, zu villen bresten [äusserlich behandelt] | 117b,112r.02.v02 | Composed mixed |
| juden öpffell | juden öpffell | Fruit | es thribt den vssatzt | vs satz, vssetzigkeitt | 117b,112r.02.v17 | Composed mixed |
| juden öpffell | juden öpffell | Fruit | zU fullen wunden | zU fullen wunden, bewertt vor fullem fleisch | 117b,112r.02.v03 | Composed mixed |
| kärnenn | kärnenn | Seed | geschwerr | geschwërr, geschwärr, geschwerr | 107a,101v.06.v01 | Composed herbal |
| katzen trübel | katzen trübel safft | Preparation | zU villen presten | presten, zu villen bresten [äusserlich behandelt] | 103b,98r.04 | Composed mixed |
| kern-gertten | kern-gertten blUst wasserr | Preparation | wunden welche full fleisch erzügt, vertribt auch die hitz | zU fullen wunden, bewertt vor fullem fleisch | 107b,102r.02.v01 | Composed herbal |
| klein schwerttel wurtz | klein schwerttel wurtz | Root | Dornn vnnd pfill vss zU ziechenn | Dornn vnnd pfill vss zU ziechenn | 156a,150v.03 | Composed herbal |
| klein winter grün | klein winter grün | Not mentioned | wund salben | wunden | 104b,99r.01 | Composed mixed |
| knaben krutt | knaben krutt | Root | heilett gruntlich die fÿgwërtzen | figwertzenn, fÿg wertzenn, fÿg Blatterenn | 121a,115v.04 | Simple herbal |
| knoblach | knoblach | Not mentioned | für denn wurm | wurm am finnger | 131b,126r.08 | Composed herbal |
| knoblach krutt | knoblach krutt | Not mentioned | Für fig wertzenn | figwertzenn, fÿg wertzenn, fÿg Blatterenn | 122a,116v.02.z01 | Composed mixed |
| korble krutt, korblin krutt | korble krutt wasserr | Preparation | Gestockt blUtt vs trÿben | Gestockt blUtt vs trÿben | 078b,72r.06 | Composed mixed |
| korble krutt, korblin krutt | korblin krutt, krutt vnnd wurtzen | Aerial part and root | wurm am finngerr | wurm am finnger | 131b,126r.04 | Simple herbal |
| küngund krutt | küngund krutt | Aerial part | die bosen schenckell, vertribt die gschwulst, so vonn keltte vnnd müde kompt | geschwulst | 052b,46r.02 | Simple herbal |
| kütenenn | kütenenn kernenn | Seed | branndt | Branndt | 136a,130v.05 | Composed herbal |
| lennger ie lieber holtz | lennger ie lieber holtz | Twig | Für alle geschwulst | geschwulst | 051b,45r.03.v01 | Simple herbal |
| lilienn | lilienn krutt blUmen wasserr | Preparation | zitter mall | zitter mall (mossen vnnd flecken) | 134a,128v.05 | Simple herbal |

**SM – Table S4 – Medicinal plant use reports** (Extract from the RBH database on dermatological recipes)

| txtPlantName_ABvH | txtDrugName_ABvH | txtPlantpart | txtCitedUse | txtCombinedUse | txtRecSig | txtRecTyp |
|---|---|---|---|---|---|---|
| linnden | rinnden (glatt mitlest) des linnden boums | Bark | branndt | Branndt | 136a,130v.03 | Simple herbal |
| lorberr, lorbonen, lorbeeri | Lorbonen, lorbeeri, Grüne lorberr | Fruit | gschwulst | geschwulst | 052b,46r.03 | Composed herbal |
| lorberr, lorbonen, lorbeeri | loröll | Preparation | für fläcken jnn angesicht | fleckenn | 117b,112r.02.v05 | Composed mixed |
| lorberr, lorbonen, lorbeeri | loröll | Preparation | verthribt alles geschwerr | geschwërr, geschwärr, geschwerr | 117b,112r.02.v12 | Composed mixed |
| lorberr, lorbonen, lorbeeri | loröll | Preparation | büllen, vnnd geschwulst | geschwulst | 117b,112r.02.v15 | Composed mixed |
| lorberr, lorbonen, lorbeeri | loröll | Preparation | Uber allen prestenn gelegt | presten, zu villen bresten [äusserlich behandelt] | 117b,112r.02.v02 | Composed mixed |
| lorberr, lorbonen, lorbeeri | loröll | Preparation | es thribt den vssatzt | vs satz, vssetzigkeitt | 117b,112r.02.v17 | Composed mixed |
| lorberr, lorbonen, lorbeeri | loröll | Preparation | wurm | wurm [allgemein] | 132a,126v.02.v01 | Composed mixed |
| lorberr, lorbonen, lorbeeri | loröll | Preparation | zU fullen wunden | zU fullen wunden, bewertt vor fullem fleisch | 117b,112r.02.v03 | Composed mixed |
| lorberr, lorbonen, lorbeeri | lorbonen: geschelltt | Seed | für alerleÿ flüs im anngesicht | flüs jm angsicht | 105b,100r.01 | Composed mixed |
| lüss krutt | lüss krutt | Not mentioned | für denn wurm | wurm am finnger | 131b,126r.09 | Composed herbal |
| macis | macis | Other | für fläcken jnn angesicht | fleckenn | 117b,112r.02.v05 | Composed mixed |
| macis | macis | Other | verthribt alles geschwerr | geschwërr, geschwärr, geschwerr | 117b,112r.02.v12 | Composed mixed |
| macis | macis | Other | büllen, vnnd geschwulst | geschwulst | 117b,112r.02.v15 | Composed mixed |
| macis | macis | Other | Uber allen prestenn gelegt | presten, zu villen bresten [äusserlich behandelt] | 117b,112r.02.v02 | Composed mixed |
| macis | macis | Other | es thribt den vssatzt | vs satz, vssetzigkeitt | 117b,112r.02.v17 | Composed mixed |
| macis | macis | Other | zU fullen wunden | zU fullen wunden, bewertt vor fullem fleisch | 117b,112r.02.v03 | Composed mixed |
| mandel | mandel (gescheltter) | Seed | für alerleÿ flüs im anngesicht | flüs jm angsicht | 105b,100r.01 | Composed mixed |
| mass blümlin, moss blumen | mass blümlin safft | Preparation | zU villen presten | presten, zu villen bresten [äusserlich behandelt] | 103b,98r.04 | Composed mixed |
| mass blümlin, moss blumen | mass blümlin, moss blUmen wasserr gebränndt | Preparation | das eim das glid Wasser Verstadt | glid Wasser Verstadt | 161b,156r.03.v02 | Composed herbal |
| mass blümlin, moss blumen | mass blümlin, moss blUmen | Not mentioned | das eim das glid Wasser | glid Wasser Verstadt | 161b,156r.03.v01 | Composed mixed |
| mastix | mastix | Gum, resin | grosse hitz zU eim schadenn schlecht die der hitz hefftig werdt | Branndt | 136b,131r.01 | Composed mixed |
| mastix | mastix | Gum, resin | das eim das glid Wasser Verstadt | glid Wasser Verstadt | 161b,156r.01 | Composed mixed |
| mastix | mastix | Gum, resin | zU allerlei Wunnden vnnd annderem | wunden | 106b,101r.02 | Composed mixed |
| meister wurtz | meister wurtz safft | Preparation | zertheillt vnnd nider geleidt werden, vnnd zur heillunng komen alle geschwulst vnnd knollen | geschwulst | 052a,45v.01 | Simple herbal |
| meister wurtz | meisterr wurtzenn bletterr samt dem safft | Leaf | für denn wurm | wurm [allgemein] | 131b,126r.11 | Composed mixed |

**SM – Table S4 – Medicinal plant use reports** (Extract from the RBH database on dermatological recipes)

| txtPlantName_ABvH | txtDrugName_ABvH | txtPlantpart | txtCitedUse | txtCombinedUse | txtRecSig | txtRecTyp |
|---|---|---|---|---|---|---|
| melissen | melissen saft | Preparation | fistell | fistel, fistalam | 117a,111v.02.v02 | Simple herbal |
| melissen | melissen saft | Preparation | werr den vs satz mitt dissem wasserr wescht offt vnnd dick es hinndertt vnnd verhütett den lanng | vs satz, vssetzigkeitt | 117a,111v.02.v04 | Simple herbal |
| melissen | melissen saft | Preparation | alle wunden, vnnd schaden, mitt dissem wasser gewäschen reinigett vnnd bewertt das vor fullem fleisch, vnnd ist gUtt für rünende wunden | zU fullen wunden, bewertt vor fullem fleisch | 117a,111v.02.v01 | Simple herbal |
| melissen | melissen Wasser | Preparation | vor alle apostema vnnd geschwer | apostema vnnd geschwer | 117b,112r.01.v03 | Simple herbal |
| melissen | melissen Wasser | Preparation | fistell | fistel, fistalam | 117a,111v.01.v02 | Simple herbal |
| melissen | melissen Wasser | Preparation | reinigett alle bösse füchtigkeitt | füchtigkeitt - reinigett füchtigkeitt | 117b,112r.01.v02 | Simple herbal |
| melissen | melissen Wasser | Preparation | gschwulst alls büllen | geschwulst | 117b,112r.01.v05 | Simple herbal |
| melissen | melissen Wasser | Preparation | werr den vs satz mitt dissem wasserr wescht offt vnnd dick es hinndertt vnnd ver-hütett den lanng | vs satz, vssetzigkeitt | 117a,111v.01.v04 | Simple herbal |
| melissen | melissen Wasser | Preparation | wer lanng sich be sorgett vor vssetzigkeitt der bruch des wasserrs | vssetzigkeitt, wer lanng sich be sorgett vor ... | 117a,111v.01.v03 | Simple herbal |
| melissen | melissen Wasser | Preparation | alle wunden, vnnd schaden, mitt dissem wasser gewäschen reinigett vnnd bewertt das vor fullem fleisch, vnnd ist gUtt für rünende wunden | zU fullen wunden, bewertt vor fullem fleisch | 117a,111v.01.v01 | Simple herbal |
| melissen | melissen krutt | Aerial part | wer lanng sich be sorgett vor vssetzigkeitt | vssetzigkeitt, wer lanng sich be sorgett vor ... | 117a,111v.02.v03 | Simple herbal |
| mer rettich | mer rettich | Not mentioned | wurm am finnger | wurm am finnger | 131a,125v.01 | Composed herbal |
| miess | miess | Not mentioned | blUtt verstellen | blUtt stellunng | 079a,72v.01 | Simple herbal |
| modelgeer | modelgeer krutt | Aerial part | ein herrlich atraitium dörnn sprisen, spinlen spitz vs ze züchen | Dornn vnnd pfill vss zU ziechenn | 156a,150v.09 | Simple herbal |
| müllj stoub (des höchstenn) | müllj stoub (des höchstenn) | Seed | Für fÿgwertzenn | figwertzenn, fÿg wertzenn, fÿg Blatterenn | 121a,115v.01 | Composed herbal |
| muscat nus, muscatt | muscat nus, muscatt | Seed | für fläcken jnn angesicht | fleckenn | 117b,112r.02.v05 | Composed mixed |
| muscat nus, muscatt | muscat nus, muscatt | Seed | für alerleÿ flüs im anngesicht | flüs jm angsicht | 105b,100r.01 | Composed mixed |
| muscat nus, muscatt | muscat nus, muscatt | Seed | verthribt alles geschwerr | geschwërr, geschwärr, geschwerr | 117b,112r.02.v12 | Composed mixed |
| muscat nus, muscatt | muscat nus, muscatt | Seed | büllen, vnnd geschwulst | geschwulst | 117b,112r.02.v15 | Composed mixed |
| muscat nus, muscatt | muscat nus, muscatt | Seed | Uber allen prestenn gelegt | presten, zu villen bresten [äusserlich behandelt] | 117b,112r.02.v02 | Composed mixed |
| muscat nus, muscatt | muscat nus, muscatt | Seed | es thribt den vssatzt | vs satz, vssetzigkeitt | 117b,112r.02.v17 | Composed mixed |
| muscat nus, muscatt | muscat nus, muscatt | Seed | zU fullen wunden | zU fullen wunden, bewertt vor fullem fleisch | 117b,112r.02.v03 | Composed mixed |
| nacht schatt, nachtschatten | nacht schatt safft, nachtschatten safft | Preparation | Für fÿgwertzenn | figwertzenn, fÿg wertzenn, fÿg Blatterenn | 121a,115v.03 | Composed mixed |
| nacht schatt, nachtschatten | nacht schatt safft, nachtschatten safft | Preparation | zU villen presten | presten, zu villen bresten [äusserlich behandelt] | 103b,98r.04 | Composed mixed |
| nacht schatt, nachtschatten | nacht schatten, nach schatten | Not mentioned | für denn wurm | wurm am finnger | 131b,126r.09 | Composed herbal |
| nacht schatt, nachtschatten | nacht schatten, nach schatten | Not mentioned | Böss fläckenn zU vertriben | zitter mall (mossen vnnd flecken) | 134a,128v.11 | Composed mixed |
| nacht schatt, nachtschatten | nacht schatten wurtzen | Root | Man sol wüssenn, das drierlein wertzen sinnd, das manns nitt mitt einerlei vertribt, die ersten vertribt man mitt ... | figwertzenn, fÿg wertzenn, fÿg Blatterenn | 121b,116r.02 | Simple herbal |

**SM – Table S4 – Medicinal plant use reports** (Extract from the RBH database on dermatological recipes)

| txtPlantName_ABvH | txtDrugName_ABvH | txtPlantpart | txtCitedUse | txtCombinedUse | txtRecSig | txtRecTyp |
|---|---|---|---|---|---|---|
| näptenn krutt | näptenn krutt | Not mentioned | Würm býss, nater biss | Würm býss, schlanngen biss, nater biss | 132a,126v.05 | Simple herbal |
| natter krutt, natterkrutt | natter krutt, natterkrutt | Not mentioned | dornn vnnden in fUss | Dornn vnnd pfill vss zU ziechenn | 156a,150v.08 | Composed mixed |
| natter krutt, natterkrutt | natter krutt, natterkrutt | Not mentioned | wund salben | wunden | 104b,99r.01 | Composed mixed |
| nëgellin, näglin | nëgellin, näglin | Flower | für fläcken jnn angesicht | fleckenn | 117b,112r.02.v05 | Composed mixed |
| nëgellin, näglin | nëgellin, näglin | Flower | verthribt alles geschwerr | geschwërr, geschwärr, geschwerr | 117b,112r.02.v12 | Composed mixed |
| nëgellin, näglin | nëgellin, näglin | Flower | büllen, vnnd geschwulst | geschwulst | 117b,112r.02.v15 | Composed mixed |
| nëgellin, näglin | nëgellin, näglin | Flower | Uber allen prestenn gelegt | presten, zu villen bresten [äusserlich behandelt] | 117b,112r.02.v02 | Composed mixed |
| nëgellin, näglin | nëgellin, näglin | Flower | es thribt den vssatzt | vs satz, vssetzigkeitt | 117b,112r.02.v17 | Composed mixed |
| nëgellin, näglin | nëgellin, näglin | Flower | zU fullen wunden | zU fullen wunden, bewertt vor fullem fleisch | 117b,112r.02.v03 | Composed mixed |
| Nesslen | Nesslen safft | Aerial part | nasën blUtten | Nasën blUtten | 079a,72v.07 | Simple herbal |
| niess wurtz | niess wurtz | Not mentioned | rott louffennde grinnd | grindt | 160a,154v.02 | Composed mixed |
| olibanum, olibanj | olibanum, olibanj | Gum, resin | zU alttenn schaden, es reiniget zücht fleisch trücknett vnnd rimpfft zU samen | zU alltenn schadenn | 114b,109r.01 | Composed mixed |
| papell, bapellenn | bapellenn krutt | Aerial part | geschwërr | geschwërr, geschwärr, geschwerr | 107a,101v.07 | Composed mixed |
| papell, bapellenn | papell wurtzen | Root | blUtt stellunng | blUtt stellunng | 078b,72r.01 | Simple herbal |
| paris körnlin | paris körnlin | Seed | für fläcken jnn angesicht | fleckenn | 117b,112r.02.v05 | Composed mixed |
| paris körnlin | paris körnlin | Seed | verthribt alles geschwerr | geschwërr, geschwärr, geschwerr | 117b,112r.02.v12 | Composed mixed |
| paris körnlin | paris körnlin | Seed | büllen, vnnd geschwulst | geschwulst | 117b,112r.02.v15 | Composed mixed |
| paris körnlin | paris körnlin | Seed | Uber allen prestenn gelegt | presten, zu villen bresten [äusserlich behandelt] | 117b,112r.02.v02 | Composed mixed |
| paris körnlin | paris körnlin | Seed | es thribt den vssatzt | vs satz, vssetzigkeitt | 117b,112r.02.v17 | Composed mixed |
| paris körnlin | paris körnlin | Seed | zU fullen wunden | zU fullen wunden, bewertt vor fullem fleisch | 117b,112r.02.v03 | Composed mixed |
| petterlj | petterlj | Not mentioned | figwertzenn die zU vertribenn | figwertzenn, fÿg wertzenn, fÿg Blatterenn | 120b,115r.03.v01 | Composed mixed |
| petterlj | petterlj | Not mentioned | wen ein mensch das harnn hatt | wen ein mensch das harnn hatt | 120b,115r.03.v02 | Composed mixed |
| pfäffer krutt | pfäffer krutt | Aerial part | Ob einner an einn ÿssenn trädten, es zücht herus | Dornn vnnd pfill vss zU ziechenn | 156b,151r.03 | Simple herbal |
| pflumen | pflumen | Fruit | Für alle geschwulst | geschwulst | 051b,45r.03.v02 | Composed herbal |
| pflumen | pflumen bletter | Leaf | Für alle geschwulst | geschwulst | 051b,45r.03.v02 | Composed herbal |
| poleÿ, boley, hertzen bleich, hertz bleÿ | poleÿ, boley, hertzen bleich, hertz bleÿ | Not mentioned | Für das juckenn | juckenn | 127a,121v.04 | Simple herbal |
| poleÿ, boley, hertzen bleich, hertz bleÿ | poleÿ, boley, hertzen bleich, hertz bleÿ | Not mentioned | für denn wurm | wurm am finnger | 131b,126r.08 | Composed herbal |
| poleÿ, boley, hertzen bleich, hertz bleÿ | Boleÿ krutt vnnd wurtz | Aerial part and root | Für nüw erhabne geschwulst | geschwulst | 051b,45r.06 | Composed herbal |
| reb krutt | reb krutt | Aerial part | Für die geschwulst | geschwulst | 051b,45r.02 | Composed herbal |
| rörlin krutt | rörlin krutt | Aerial part | für denn wurm | wurm am finnger | 131b,126r.09 | Composed herbal |
| rosenn wurtzell | rosenn wurtzell | Root | edle blUtt stellung | blUtt stellunng | 079b,73r.01 | Composed herbal |
| rosenn, ross | ross essich | Preparation | grosse hitz zU eim schadenn schlecht die der hitz hefftig werdt | Branndt | 136b,131r.01 | Composed mixed |
| rosenn, ross | rosenn blüett | Flower | Wertzen | Wertzenn, wartzen | 122a,116v.04 | Simple herbal |
| rosenn, ross | ross wasser | Preparation | branndt | Branndt | 136a,130v.05 | Composed herbal |
| rosenn, ross | ross wasser | Preparation | einn angesicht als obs vss setzig sige | vs satz, vssetzigkeitt | 128b,123r.01 | Composed mixed |
| rosenn, ross | ross wasser | Preparation | zU offnenn schaden Vnnd flüssenn | zU offnenn schaden Vnnd flüssenn | 105b,100r.03 | Composed mixed |

**SM – Table S4 – Medicinal plant use reports** (Extract from the RBH database on dermatological recipes)

| txtPlantName_ABvH | txtDrugName_ABvH | txtPlantpart | txtCitedUse | txtCombinedUse | txtRecSig | txtRecTyp |
|---|---|---|---|---|---|---|
| rosenn, ross | ross öll | Preparation | gUtt blatterenn | Blattern | 107a,101v.08.v02 | Composed mixed |
| rosenn, ross | ross öll | Preparation | grosse hitz zU eim schadenn schlecht die der hitz hefftig werdt | Branndt | 136b,131r.01 | Composed mixed |
| rosenn, ross | ross öll | Preparation | Branndt Löschenn | Branndt | 136a,130v.01 | Composed herbal |
| rosenn, ross | ross öll | Preparation | eÿssenn | eÿssenn | 107a,101v.08.v01 | Composed mixed |
| rosenn, ross | ross öll | Preparation | Für gschwulst so vonn überiger hitz kombt es sig vonn wundern. törnn. stëchen schlahen oder gfallen | geschwulst vonn überiger hitz kombt, es sig vonn wundern, törnn, stëchen, schlahen oder gfallen | 052a,45v.02 | Composed mixed |
| rosenn, ross | ross öll | Preparation | zU offnenn schaden Vnnd flüssenn | zU offnenn schaden Vnnd flüssenn | 105b,100r.03 | Composed mixed |
| rot bugglenn | rott bugenn die oberistenn dölderlin | Tip of shoot | figwertzenn die zU vertribenn | figwertzenn, fÿg wertzenn, fÿg Blatterenn | 120b,115r.03.v01 | Composed mixed |
| rot bugglenn | rott bugenn die oberistenn dölderlin | Tip of shoot | wen ein mensch das harnn hatt | wen ein mensch das harnn hatt | 120b,115r.03.v02 | Composed mixed |
| rot bugglenn | rot bugglenn | Not mentioned | Für die geschwulst | geschwulst | 051b,45r.02 | Composed herbal |
| rott danj | rott danj hartz | Gum, resin | Houwt, oder sticht oder brent oder böss eissenn am Lÿb Hatt | Houwt, oder sticht oder brent oder böss eissenn am Lÿb Hatt | 106b,101r.01 | Composed mixed |
| rotten mangolt | rotten mangolt | Not mentioned | gUtt blatterenn | Blattern | 107a,101v.08.v02 | Composed mixed |
| rotten mangolt | rotten mangolt | Not mentioned | eÿssenn | eÿssenn | 107a,101v.08.v01 | Composed mixed |
| rotten mangolt | rotten mangolt | Not mentioned | wund salben | wunden | 104b,99r.01 | Composed mixed |
| ruten, rutten | rutten | Not mentioned | Wem blUtt zwüschen hutt vnnd fleisch | blUtt zwüschen hutt vnnd fleisch | 079b,73r.04 | Composed herbal |
| ruten, rutten | rutten | Not mentioned | für fläcken jnn angesicht | fleckenn | 117b,112r.02.v05 | Composed mixed |
| ruten, rutten | rutten | Not mentioned | verthribt alles geschwerr | geschwërr, geschwärr, geschwerr | 117b,112r.02.v12 | Composed mixed |
| ruten, rutten | rutten | Not mentioned | gschwulst | geschwulst | 052b,46r.03 | Composed herbal |
| ruten, rutten | rutten | Not mentioned | Uber allen prestenn gelegt | presten, zu villen bresten [äusserlich behandelt] | 117b,112r.02.v02 | Composed mixed |
| ruten, rutten | rutten | Not mentioned | es thribt den vssatzt | vs satz, vssetzigkeitt | 117b,112r.02.v17 | Composed mixed |
| ruten, rutten | rutten | Not mentioned | zum wurm | wurm [allgemein] | 131b,126r.10 | Composed herbal |
| ruten, rutten | rutten | Not mentioned | zU fullen wunden | zU fullen wunden, bewertt vor fullem fleisch | 117b,112r.02.v03 | Composed mixed |
| ruten, rutten | ruten krutt | Aerial part | Wurms annfanng; wurm am finnger | wurm am finnger | 131a,125v.02 | Composed mixed |
| ruten, rutten | rutten öll | Preparation | Schrunnden an hennden | Schrunnden an hennden | 107a,101v.09 | Composed mixed |
| ruten, rutten | rutten körnnerr | Seed | zitter mall: mossen vnnd flecken | zitter mall (mossen vnnd flecken) | 134a,128v.03 | Composed mixed |
| saffrann | saffrann | Not mentioned | grosse hitz zU eim schadenn schlecht die der hitz hefftig werdt | Branndt | 136b,131r.01 | Composed mixed |
| saffrann | saffrann | Not mentioned | wunden | wunden | 108a,102v.01.v01 | Composed mixed |
| safft, so vs dem holtz flüst das jm fürr ligt | safft, so vs dem holtz flüst das jm fürr ligt | Other | Wertzen | Wertzenn, wartzen | 122a,116v.05 | Simple herbal |
| salbinenn | salbinenn | Aerial part | wunnd salben | wunden | 106a,100v.05 | Composed mixed |
| sanickel | Sanickel | Aerial part | blUtt ze stellen | blUtt stellunng | 079a,72v.08 | Composed herbal |
| sanickel | Sanickel | Aerial part | wund salben | wunden | 104b,99r.01 | Composed mixed |
| sanndelholtz | sanndelholtz | Wood | thUn das vnder ein anderen thU das buluer in die wunden es stadt für warr [blUtt ze stellen] | blUtt stellunng | 079a,72v.05 | Composed herbal |
| sannt petters krutt | Sannt petters krutt | Not mentioned | für die gschwulst, der beinenn vnnd schëncklen | geschwulst | 052a,45v.05.v01 | Simple herbal |

**SM – Table S4 – Medicinal plant use reports** (Extract from the RBH database on dermatological recipes)

| txtPlantName_ABvH | txtDrugName_ABvH | txtPlantpart | txtCitedUse | txtCombinedUse | txtRecSig | txtRecTyp |
|---|---|---|---|---|---|---|
| sanntt johanns krutt | sanntt johanns krutt vnnd der selbenn blUmen | Flowering herb | wunden | wunden | 108a,102v.01.v01 | Composed mixed |
| sant barbel krutt | sant barbel krutt | Aerial part | dornn vnnden in fUss | Dornn vnnd pfill vss zU ziechenn | 156a,150v.08 | Composed mixed |
| schellkrutt, schelkrutt, schell krutt | schellkrutt, schelkrutt, schell krutt | Not mentioned | wund salben | wunden | 104b,99r.01 | Composed mixed |
| schlechenn, schlehen dornn | Laub vonn schlechenn puschenn | Leaf | Für alle geschwulst | geschwulst | 052b,46r.05 | Simple herbal |
| schlechenn, schlehen dornn | blUst vonn eim schlehen dornn | Flower | so aber das blUtt selber kompt von der nasen | Nasën blUtten | 079a,72v.03 | Simple herbal |
| schlüssell blumen | schlüssell blUmen wurtzen | Root | an einn ÿssenn trädten, es zücht herus | Dornn vnnd pfill vss zU ziechenn | 156b,151r.02 | Simple herbal |
| schwalmenn wurzell | Schwalmenn wurzell | Aerial part and root | Branndt | Branndt | 136b,131r.02 | Simple herbal |
| schwalmenn wurzell | Schwalmenn wurzell | Root | für alle gschwulst | geschwulst | 136b,131r.03 | Simple herbal |
| schwartze nacht schatten | schwartze nacht schatten beerr | Fruit | Für nüw erhabne geschwulst | geschwulst | 051b,45r.06 | Composed herbal |
| scritten | scritten | Not mentioned | wurm am finngerr | wurm am finnger | 131b,126r.01 | Composed herbal |
| sennff krutt | sennff krutt | Aerial part | dornn vnnden in fUss | Dornn vnnd pfill vss zU ziechenn | 156a,150v.08 | Composed mixed |
| serpentia wurtzen | serpentia wurtzen | Root | verthribt auch fleckenn | fleckenn | 106b,101r.04.v03 | Composed herbal |
| serpentia wurtzen | serpentia wurtzen | Root | sÿ süberett auch wunden | wunden | 106b,101r.04.v02 | Composed herbal |
| seueboum | seueboum, denn obersten dolder | Tip of shoot | wurm am finngerr | wurm am finnger | 131a,125v.10 | Composed herbal |
| seueboum | seueboum | Not mentioned | für denn wurm | wurm am finnger | 131b,126r.08 | Composed herbal |
| sigilis solomonis | sigilis solomonis wurtzell | Root | zitter mall | zitter mall (mossen vnnd flecken) | 134a,128v.06 | Composed mixed |
| sinauw | sinauw | Aerial part | blUtt ze stellen | blUtt stellunng | 079a,72v.08 | Composed herbal |
| sinauw | sinauw | Aerial part | figwertzenn die zU vertribenn | figwertzenn, fÿg wertzenn, fÿg Blatterenn | 120b,115r.02 | Composed mixed |
| spickennardj | spickennardj | Other | für fläcken jnn angesicht | fleckenn | 117b,112r.02.v05 | Composed mixed |
| spickennardj | spickennardj | Other | verthribt alles geschwerr | geschwërr, geschwärr, geschwerr | 117b,112r.02.v12 | Composed mixed |
| spickennardj | spickennardj | Other | büllen, vnnd geschwulst | geschwulst | 117b,112r.02.v15 | Composed mixed |
| spickennardj | spickennardj | Other | Uber allen prestenn gelegt | presten, zu villen bresten [äusserlich behandelt] | 117b,112r.02.v02 | Composed mixed |
| spickennardj | spickennardj | Other | es thribt den vssatzt | vs satz, vssetzigkeitt | 117b,112r.02.v17 | Composed mixed |
| spickennardj | spickennardj | Other | zU fullen wunden | zU fullen wunden, bewertt vor fullem fleisch | 117b,112r.02.v03 | Composed mixed |
| spickennardj | oleum spice | Preparation | für die rud | rud | 127b,122r.05 | Composed mixed |
| spitzwegrich, spitzen wägrich | spitzwegrich safft, safft vonn spitzen wägrich | Preparation | als kräps, wolff, nolime tangere vnnd des glich | kräps, wolff, noli me tangere, vnnd des glich | 114b,109r.05.v02 | Composed mixed |
| spitzwegrich, spitzen wägrich | spitzwegrich safft, safft vonn spitzen wägrich | Preparation | zU villen presten | presten, zu villen bresten [äusserlich behandelt] | 103b,98r.04 | Composed mixed |
| spitzwegrich, spitzen wägrich | spitzwegrich safft, safft vonn spitzen wägrich | Preparation | rünende bein | rünende bein | 114b,109r.05.v03 | Composed mixed |
| spitzwegrich, spitzen wägrich | spitzwegrich, spitzen wegrich, spitzen wägrich | Not mentioned | wund salben | wunden | 104b,99r.01 | Composed mixed |

**SM – Table S4 – Medicinal plant use reports** (Extract from the RBH database on dermatological recipes)

| txtPlantName_ABvH | txtDrugName_ABvH | txtPlantpart | txtCitedUse | txtCombinedUse | txtRecSig | txtRecTyp |
|---|---|---|---|---|---|---|
| spitzwegrich, spitzen wägrich | spitzwegrich, spitzen wegrich, spitzen wägrich | Not mentioned | allt schaden, allte ruden | zU alltenn schadenn | 114b,109r.05.v01 | Composed mixed |
| stinckende nesslen | stinckende nesslen krutt | Aerial part | wurm am finngerr | wurm am finnger | 131b,126r.01 | Composed herbal |
| stritten | stritten | Not mentioned | zum wurm | wurm [allgemein] | 131b,126r.10 | Composed herbal |
| süss-holtz | süss-holtz | Not mentioned | Für nüw erhabne geschwulst | geschwulst | 051b,45r.06 | Composed herbal |
| terpenntin, terpetinum, terprini, terpetin, terpatin, Derpentin | terpenntin, terpetinum, terprini, terpetin, terpatin, Derpentin | Gum, resin | das eim das glid Wasser Verstadt | glid Wasser Verstadt | 161b,156r.01 | Composed mixed |
| terpenntin, terpetinum, terprini, terpetin, terpatin, Derpentin | terpenntin, terpetinum, terprini, terpetin, terpatin, Derpentin | Gum, resin | für die rud | rud | 127b,122r.05 | Composed mixed |
| terpenntin, terpetinum, terprini, terpetin, terpatin, Derpentin | terpenntin, terpetinum, terprini, terpetin, terpatin, Derpentin | Gum, resin | zU frischen wunndenn | wunden | 106a,100v.01 | Composed mixed |
| terpenntin, terpetinum, terprini, terpetin, terpatin, Derpentin | terpenntin, terpetinum, terprini, terpetin, terpatin, Derpentin | Gum, resin | zU alltenn schaden, es reiniget zücht fleisch trücknett vnnd rimpfft zU samen | zU alltenn schadenn | 114b,109r.01 | Composed mixed |
| tubenn kröpfflin | Tubenn kröpfflin wasser | Preparation | so einner grunenn blUtt bÿ jm hatt | blUtt grunnen | 116b,111r.01.v06 | Composed herbal |
| tubenn kröpfflin | Tubenn kröpfflin wasser | Preparation | flüs jm angsicht | flüs jm angsicht | 116b,111r.01.v05 | Composed herbal |
| tubenn kröpfflin | Tubenn kröpfflin wasser | Preparation | für alle kretz vnd rüdigkeitt | kretz vnd rüdigkeitt | 116b,111r.01.v02 | Composed herbal |
| tubenn kröpfflin | Tubenn kröpfflin wasser | Preparation | für die rud Vnnd denn magerr | rud vnnd magerr | 127a,121v.01 | Simple herbal |
| tubenn kröpfflin | Tubenn kröpfflin wasser | Preparation | gUtt den menschen, so sich förchten vor der vss setzig keitt | vssetzigkeitt, wer lanng sich be sorgett vor | 116b,111r.01.v01 | Composed herbal |
| tubenn kröpfflin | Tubenn kröpfflin gebrannt wie rossenn | Preparation | Flüss jm anngesicht | flüs jm angsicht | 128b,123r.02 | Simple herbal |
| venumgrecum | venumgrecum | Seed | das eim das glid Wasser Verstadt | glid Wasser Verstadt | 161b,156r.07 | Composed mixed |
| verbena | verbena | Aerial part | zU allerlei Wunnden vnnd annderem | wunden | 106b,101r.02 | Composed mixed |
| viöl | viöl krutt | Aerial part | Man sol wüssenn, das drierlein wertzen sinnd, das manns nitt mitt einerlei vertribt, ... die anderen vertribt man mitt ... | figwertzenn, fÿg wertzenn, fÿg Blatterenn | 121b,116r.03 | Simple herbal |
| viönlj, vigell | viönlj öll, vigell öll | Preparation | gUtt blatterenn | Blattern | 107a,101v.08.v02 | Composed mixed |
| viönlj, vigell | viönlj öll, vigell öll | Preparation | eÿssenn | eÿssenn | 107a,101v.08.v01 | Composed mixed |
| viönlj, vigell | viönlj öll, vigell öll | Preparation | Für fÿgwertzenn | figwertzenn, fÿg wertzenn, fÿg Blatterenn | 121a,115v.02 | Simple herbal |
| viönlj, vigell | viönlj öll, vigell öll | Preparation | zU offnenn schaden Vnnd flüssenn | zU offnenn schaden Vnnd flüssenn | 105b,100r.03 | Composed mixed |
| vnnserr frouwen mentle | vnnserr frouwen mentle | Not mentioned | Für fig wertzenn | figwertzenn, fÿg wertzenn, fÿg Blatterenn | 122a,116v.02 | Simple herbal |
| voglj krutt | voglj krutt | Not mentioned | figwertzenn die zU vertribenn | figwertzenn, fÿg wertzenn, fÿg Blatterenn | 120b,115r.02 | Composed mixed |
| wal wurtzen | wal wurtzen | Not mentioned | wund salben | wunden | 104b,99r.01 | Composed mixed |
| wald farn [grosse] | wald farn wurtzen [grosse] | Root | zum wurm | wurm [allgemein] | 131b,126r.10 | Composed herbal |
| wald meister | wald meister | Not mentioned | wund salben | wunden | 104b,99r.01 | Composed mixed |

**SM – Table S4 – Medicinal plant use reports** (Extract from the RBH database on dermatological recipes)

| txtPlantName_ABvH | txtDrugName_ABvH | txtPlantpart | txtCitedUse | txtCombinedUse | txtRecSig | txtRecTyp |
|---|---|---|---|---|---|---|
| weg gras | weg gras wasserr | Preparation | zU offnenn schaden Vnnd flüssenn | zU offnenn schaden Vnnd flüssenn | 105b,100r.03 | Composed mixed |
| wegrich, wëgrich | wegrich samen | Seed | thUn das vnder ein anderen thU das buluer in die wunden es stadt für warr [blUtt ze stellen] | blUtt stellunng | 079a,72v.05 | Composed herbal |
| wermUtt | wermUtt | Flowering herb | Wem blUtt zwüschen hutt vnnd fleisch | blUtt zwüschen hutt vnnd fleisch | 079b,73r.04 | Composed herbal |
| weÿrauch, wÿssen wieruch, -wierauch | weÿrauch | Gum, resin | Dornn vnnd pfill vss zU ziechenn | Dornn vnnd pfill vss zU ziechenn | 156a,150v.03 | Composed herbal |
| weÿtzenn | weÿtzenn | Seed | geschwërr | geschwërr, geschwärr, geschwerr | 107a,101v.06.v02 | Composed herbal |
| wilden boleÿ | wilden boleÿ | Not mentioned | wund salben | wunden | 104b,99r.01 | Composed mixed |
| wisse mülleblümlin | wisse mülleblümlin, sampt dem krutt | Aerial part and root | wund salben | wunden | 104b,99r.01 | Composed mixed |
| wollen krut, wull krutt (mitt denn gellen blUmen) | wull krutt (mitt denn gellen blUmen) | Not mentioned | Für die geschwulst | geschwulst | 051b,45r.02 | Composed herbal |
| wollen krut, wull krutt (mitt denn gellen blUmen) | wollen krut wurtz | Root | branndt, es sige mitt wasserr, oder mitt fürr | Branndt | 136a,130v.04 | Composed mixed |
| wund blümlin | wund blümlin krutt | Aerial part | wunnd salben | wunden | 106a,100v.05 | Composed mixed |
| wÿssen nacht schatten | wÿssen nacht schatten | Not mentioned | bösse hitzige gesch wulsten schennckel | geschwulst | 052a,45v.04.v01 | Composed mixed |
| wÿssen nacht schatten | wÿssen nacht schatten | Not mentioned | schonn röttj | schonn röttj | 052a,45v.04.v02 | Composed mixed |
| ÿbschenn | ÿbschenn safft | Preparation | allerleÿ geschwulst | geschwulst | 106b,101r.05.v02 | Composed mixed |
| ÿbschenn | ÿbschenn safft | Preparation | wunden | wunden | 106b,101r.05.v01 | Composed mixed |
| ÿbschenn | ÿbschenn bletter | Leaf | Würm bÿss zeheillenn | Würm bÿss, schlanngen biss, nater biss | 132a,126v.04 | Simple herbal |
| ÿbschenn | ÿbschenn | Not mentioned | Drüssenn vertribenn | Drüssenn | 122a,116v.06 | Composed herbal |
| ÿbschenn | ÿbschenn | Not mentioned | ist gUtt für fläcken | fleckenn | 122a,116v.07 | Composed mixed |
| ÿbschenn | ÿbschenn | Not mentioned | wunden die full ist, die macht sÿ rein | zU fullen wunden, bewertt vor fullem fleisch | 134b,129r.02 | Simple herbal |
| ÿbschenn | ÿbschenn wurtzenn | Root | geschwërr | geschwërr, geschwärr, geschwerr | 107a,101v.07 | Composed mixed |
| zimett | zimett | Bark | für alerleÿ flüs im anngesicht | flüs jm angsicht | 105b,100r.01 | Composed mixed |
| zimett | zimett | Bark | Für nüw erhabne geschwulst | geschwulst | 051b,45r.04 | Composed mixed |
| zitt lossen | zitt lossen mitt sampt der wurtzen | Aerial part and root | Pfill vs Ziechenn | Dornn vnnd pfill vss zU ziechenn | 156a,150v.06 | Composed herbal |
| zittwan | zittwan holtz | Root | thUn das vnder ein anderen thU das buluer in die wunden es stadt für warr [blUtt ze stellen] | blUtt stellunng | 079a,72v.05 | Composed herbal |

**Dermatological plants and their uses in the *Receptarium* of Burkhard III from Hallwyl (RBH) from 16th century Switzerland – Data mining a historical text and preliminary *in vitro* screening –** Jonas Stehlin, Ina Albert, Thomas Frei, Barbara Frei Haller, Andreas Lardos – February 2024.

**SM – Table S5 – Details of the Hallwyl wound potion on folio 251a in *FA von Hallwyl A 814*.** The text describes one basic remedy and three supplementary variants, representing four individual recipes of herbal compound preparations.

| Recipe | Recipe text | RBH plant name | Candidate plant | Plant part | RBH use | Use interpretation | Pharmacological properties |
|---|---|---|---|---|---|---|---|
| Basic remedy | Also nimm Rote Mangolt, Heydnisch Wundkrut, Klein Wintergrün, Rot Buggen, Sinouw, Sanickel. Die Kräuter sind, alle in gleicher Quantität, an der Luft getrocknet, so dass keine Sonne dazukommt, dann zu Pulver gemacht (gestossen, gerieben), untereinander gemischt. Alle diese Kräuter soll man im August sammeln. Und wenn sich jemand schneidet oder sticht, dass es offen ist, soll man von diesem Pulver soviel nehmen wie eine Walnuss, tue es in einen glasierten neuen Topf (mit Ausguss) und ein gutes altes Mass weissen Wein darüber, gut zugedeckt, so dass kein Dampf entweichen kann, dann allmählich, ohne dass es überläuft, kochen lassen so lange wie ein hartes Ei gekocht werden soll, dann vom Feuer genommen, gut zugedeckt und dem Menschen alle Tage ein gutes Tränkchen eingeben, nämlich am Morgen nüchtern, zu Mittag nach dem Essen und zu Nacht zwei Stunden nach dem Nachtessen, wenn er nichts mehr essen noch trinken will. So soll er am Morgen und am Mittag stets zwei Stunden nüchtern bleiben und der Trank soll stets in einem Gläslein gewärmt und warm getrunken werden. | rotten mangolt | Beta vulgaris L. (cultivar) (1) | Not specified | schneidet oder sticht, dass es offen ist | Open stab or cut wounds | antibiotic, anti-inflammatory, analgesic, haemostatic[2] |
| | | heydnisch wundkrut | Anthyllis vulneraria L., Senecio ovatus Willd. (1); Actaea spicata L.?, Hieracium murorum aggr.?, Lactuca muralis (L.) E.Mey.?, Saxifraga rotundifolia L.?, Scrophularia nodosa L.?, Senecio hercynicus Herborg?, Solidago virgaurea L.? | Not specified | | | |
| | | klein wintergrün | Pyrola minor L., P. rotundifolia Fern., P. secunda L. (2); Vinca minor L.?, Polygala chamaebuxus L.?, Lycopodium clavatum L.? | Not specified | | | |
| | | rot buggen | Artemisia vulgaris L. (1); Artemisia campestris L.?, Amaranthus blitum L.?, Rumex obtusifolius L.?, Portulaca oleracea L.? | Not specified | | | |
| | | sinouw | Alchemilla xanthochlora agg. (A. vulgaris L., A. xanthochlora Rothm.) (1) | Not specified | | | |
| | | sanickel | Sanicula europaea L. (1) | Not specified | | | |
| Variant 1 | Wenn aber jemand eine Spindelspitze, einen Dorn, einen Büchsenstein oder etwas anderes in sich hat, so soll man unter das Pulver einen Zweig Sevenbaum (seuebalmen) tun und ihn darin kochen lassen, | seue balmen[1] | Juniperus sabina L. (1) | Twig | eine Spindelspitze, einen Dorn, einen Büchsenstein oder etwas anderes in sich | Injury from a mandrel, thorn, shotgun bullet or from another object | antibiotic, anti-inflammatory, analgesic, haemostatic[2] |

**Dermatological plants and their uses in the *Receptarium* of Burkhard III from Hallwyl (RBH) from 16th century Switzerland – Data mining a historical text and preliminary *in vitro* screening –** Jonas Stehlin, Ina Albert, Thomas Frei, Barbara Frei Haller, Andreas Lardos – February 2024.

| Recipe | Recipe text | RBH plant name | Candidate plant | Plant part | RBH use | Use interpretation | Pharmacological properties |
|---|---|---|---|---|---|---|---|
| Variant 2 | … oder wenn der Eiter an einer Verwundung nicht gehen will, so soll man seffe reintun. | seffe[1] | Juniperus sabina L. (3); Calluna vulgaris Hull?, Juniperus communis var. saxatilis Pall. (J. nana)? | Not specified | Eiter an einer Verwundung nicht gehen will | Purulent poorly healing wound | antibiotic, anti-inflammatory, immunomodulatory[2] |
| Variant 3 | Wenn es aber schnell heilen soll, so soll man Waldholunder reintun. | waldholunder[1] | Sambucus racemosus L. (1) | Not specified | Verwundung, schnell heilen soll | To speed the healing of a wound or injury | antibiotic, anti-inflammatory, immunomodulatory[3] |

**Recipe text:** Original text of the recipe on folio 251a in *FA von Hallwyl A 814* transcribed and translated from early New High German into modern Standard German.
**Candidate plants:** Data was adopted from **Table 2 – Candidate species**.
**RBH use:** Data was adopted from **Table 1 – Medicinal uses**.
**Use interpretation:** Data was adopted from **Table 1 – Medicinal uses**.
**Pharmacological properties:** If, according to Fitzpatrick's Dermatology (Kang et al., 2019), a pharmacological intervention is available for the treatment of the medical complaint(s) listed in the column "interpreted uses", the pharmacological properties of the medication mentioned in the textbook are listed. Data was adopted from **Table 1 – Medicinal uses**.

**Footnotes:**
1: The plant is added to the herbs of the basic remedy and prepared together with them.
2: The Hallwyl wound potion is an orally administered preparation. If this preparation were to have any beneficial effect on the mentioned dermatological conditions, the effect in question would have to be achieved by internal application. The pharmacological intervention in severe traumatic injuries with open or infected wounds, the main indication of the wound potion, includes not only a topical but often also an internal application of medication with antibiotic, anti-inflammatory, analgesic or haemostatic properties (Kang et al., 2019).
3: The Hallwyl wound potion is an orally administered preparation. If this preparation were to have any beneficial effect on wound healing as purported by the historical use, the effect in question would have to be achieved by internal application. Medication with antibiotic, anti-inflammatory or immunomodulatory properties is generally considered to promote wound healing. In some dermatological conditions also an internal application of such medication is recommended (Kang et al. (2019).

**Dermatological plants and their uses in the *Receptarium* of Burkhard III from Hallwyl (RBH) from 16th century Switzerland – Data mining a historical text and preliminary *in vitro* screening –** Jonas Stehlin, Ina Albert, Thomas Frei, Barbara Frei Haller, Andreas Lardos – February 2024.

### SM – Table S6 – Details of the four dermatological recipes from the RBH database that contain the plants selected for preliminary *in vitro* screening.

| Resource and recipe type | Recipe text | Plant name | Candidate plant [Family] | Plant part | RBH use | Use interpretation | Pharmacological properties |
|---|---|---|---|---|---|---|---|
| RBH database, recipe 051b,45r.03.v01<br>Herbal simple | Für alle geschwulst - nim lennger ie lieber holtz, in wÿnn gesotten wol in. vnnd den selbigen als warm man mag, so man schlaffen wÿll gan jn nemen, zetrinckenn | lennger ie lieber | *Solanum dulcamara* L. (1) [Solanaceae]; *Lonicera periclymenum* L., *L. caprifolium* L. (2) [Caprifoliaceae] | Twig | geschwulst<br>(für alle geschwulst) | Swelling of the tissue resulting from inflammation or infection (1); Dropsy (e.g. lymphedema) (2); Tumour (3) | antibiotic, anticancer, anti-inflammatory, antineoplastic |
| RBH database, recipe 121b,116r.02<br>Herbal simple | Einn gUtt stuck wie man alerleÿ fig wertzenn vertribenn soll - man sol wüssenn, das drierlein wertzen sinnd, das manns nitt mitt einerlei vertribt, die ersten vertribt man mitt nacht schatten dauon soll man nen die wurtzen, vnnd die woll in einem mörsell stossen, vnnd das safft durch ein thüchlin trucken, vnnd ein lind thüchlin darin nass machen, vnnd die fig wertzen dar mitt bestrichen, vnnd alls tick wider netzen, so offt es trochen wirtt, | nacht schatten | *Solanum nigrum* L. (1); *S. dulcamara* L.? [Solanaceae] | Root | fig wertzenn<br>(fig wertzenn, die ersten) | Genital or syphilitic warts (Condylomata acuminata or lata) (1); Haemorrhoids (2); Neoplastic skin chance, swelling or eczema on the anus (3) | antibiotic, anti-inflammatory, antineoplastic, antiproliferative, antiviral, immunomodulatory |
| RBH database, recipe 136b,131r.02<br>Herbal simple | Branndt, Einn annders – schwalmennwurtzell, jnn win gelegt, vnnd den branndt darmitt gewäschenn | schwalmenn-wurzell | *Vincetoxicum hirundinaria* Medik. [Apocynaceae] | Not specified | branndt<br>(branndt) | Burn wound (1); Severe inflammation or ulcer (2); Gangrenous ergotism (Saint Anthony's fire) (3) | antibiotic, anti-inflammatory, analgesic, antipruritic |
| RBH database, recipe 136b,131r.03<br>Herbal simple | Für alle gschwulst - glicher gstallt die wurtzell jnn winn gesottenn vnnd über bunden | schwalmenn-wurzell | *Vincetoxicum hirundinaria* Medik. [Apocynaceae] | Root | geschwulst<br>(für alle geschwulst) | Swelling (topical or internal) of the tissue resulting from inflammation or infectious disease (1); Tumour or dropsy (2) | antibiotic, anticancer, anti-inflammatory, antineoplastic |

**Recipe text:** Transcription of the original text by Fankhauser (2012) of the recipe written by *Hand A* in *FA von Hallwyl A 814*.
**Candidate plants:** Data was adopted from **Table 2 – Candidate species**.
**RBH use:** Data was adopted from **Table 1 – Medicinal uses**.
**Use interpretation:** Data was adopted from **Table 1 – Medicinal uses**.
**Pharmacological properties:** If, according to Fitzpatrick's Dermatology (Kang et al., 2019), a pharmacological intervention is available for the treatment of the medical complaint(s) listed in the column "interpreted uses", the pharmacological properties of the medication mentioned in the textbook are listed. Data was adopted from **Table 1 – Medicinal uses**.
